# Using network control theory to study the dynamics of the structural connectome

**DOI:** 10.1101/2023.08.23.554519

**Authors:** Linden Parkes, Jason Z. Kim, Jennifer Stiso, Julia K. Brynildsen, Matthew Cieslak, Sydney Covitz, Raquel E. Gur, Ruben C. Gur, Fabio Pasqualetti, Russell T. Shinohara, Dale Zhou, Theodore D. Satterthwaite, Dani S. Bassett

## Abstract

Network control theory (NCT) is a simple and powerful tool for studying how network topology informs and constrains dynamics. Compared to other structure-function coupling approaches, the strength of NCT lies in its capacity to predict the patterns of external control signals that may alter dynamics in a desired way. We have extensively developed and validated the application of NCT to the human structural connectome. Through these efforts, we have studied (i) how different aspects of connectome topology affect neural dynamics, (ii) whether NCT outputs cohere with empirical data on brain function and stimulation, and (iii) how NCT outputs vary across development and correlate with behavior and mental health symptoms. In this protocol, we introduce a framework for applying NCT to structural connectomes following two main pathways. Our primary pathway focuses on computing the *control energy* associated with transitioning between specific neural activity states. Our second pathway focuses on computing *average controllability*, which indexes nodes’ general capacity to control dynamics. We also provide recommendations for comparing NCT outputs against null network models. Finally, we support this protocol with a Python-based software package called *network control theory for python (nctpy)*.

## I. INTRODUCTION

Network neuroscience is principally concerned with studying the connectome [1], the description of whole brain connectivity. This connectome is often encoded as a graph of nodes interconnected by edges that can be defined across multiple scales, species, and data modalities [2, 3]. In any case, these descriptions of brain connectivity give rise to complex topology—including hubs, modules, small-worldness, and core-periphery structure [4]—and understanding how this topology shapes and constrains the brain’s rich repertoire of dynamics is a central goal of network neuroscience.

Network control theory (NCT) provides an approach to studying these dynamics that yields insights into how they emerge from the topology of the underlying structural connectome [5–8]. The application of NCT has revolutionized both the understanding and design of complex networks in contexts as diverse as space and terrestrial exploration, as well as modeling of financial markets, airline networks, and fire-control systems. Briefly, NCT assumes that inter-nodal communication follows a linear model of diffusion, where activity from one set of nodes (i.e., an initial *state*) spreads across the network over time along a series of fronts [4, 9]. Then, upon this dynamical system, NCT models a set of external control signals that are designed to guide these diffusing activity patterns towards a chosen target *state*. This choice can be informed by a measurement of activity evoked by behavior, spontaneous activity, or the type of brain system. These control signals are found by minimizing the total magnitude of their input over a given time horizon; that is, they are designed to achieve a desired state transition with the lowest amount of *control energy*. Once modeled, these control signals can be examined to determine to what extent, and how, they were constrained by topology, thus allowing researchers to study how the connectome might be leveraged to control dynamics.

Recently, we have developed and tested the application of NCT to brain network data across multiple contexts, scales, and definitions of connectivity [10–20]. Here, we present a protocol for applying NCT to two different structural connectomes: one defined using undirected connectivity estimated in the human brain [21, 22] and the other using directed connectivity estimated in the mouse brain [23–25]. Briefly, we detail two common applications of NCT that we—as well as other groups [26–32]—have deployed that focus on (i) quantifying the amount of energy that is required to complete transitions between specific brain states (Figure 1) and (ii) examining regions’ general capacity to control dynamics (Figure 2). The former approach is useful for researchers interested in examining how dynamics can be controlled to move from one place on the network to another, while the latter is useful for researchers interested in analyzing topographic maps of control. Additionally, we provide recommendations for visualization of model outputs, and discuss the use of null network models to examine which topological features affect model outputs.

**FIG. 1.**
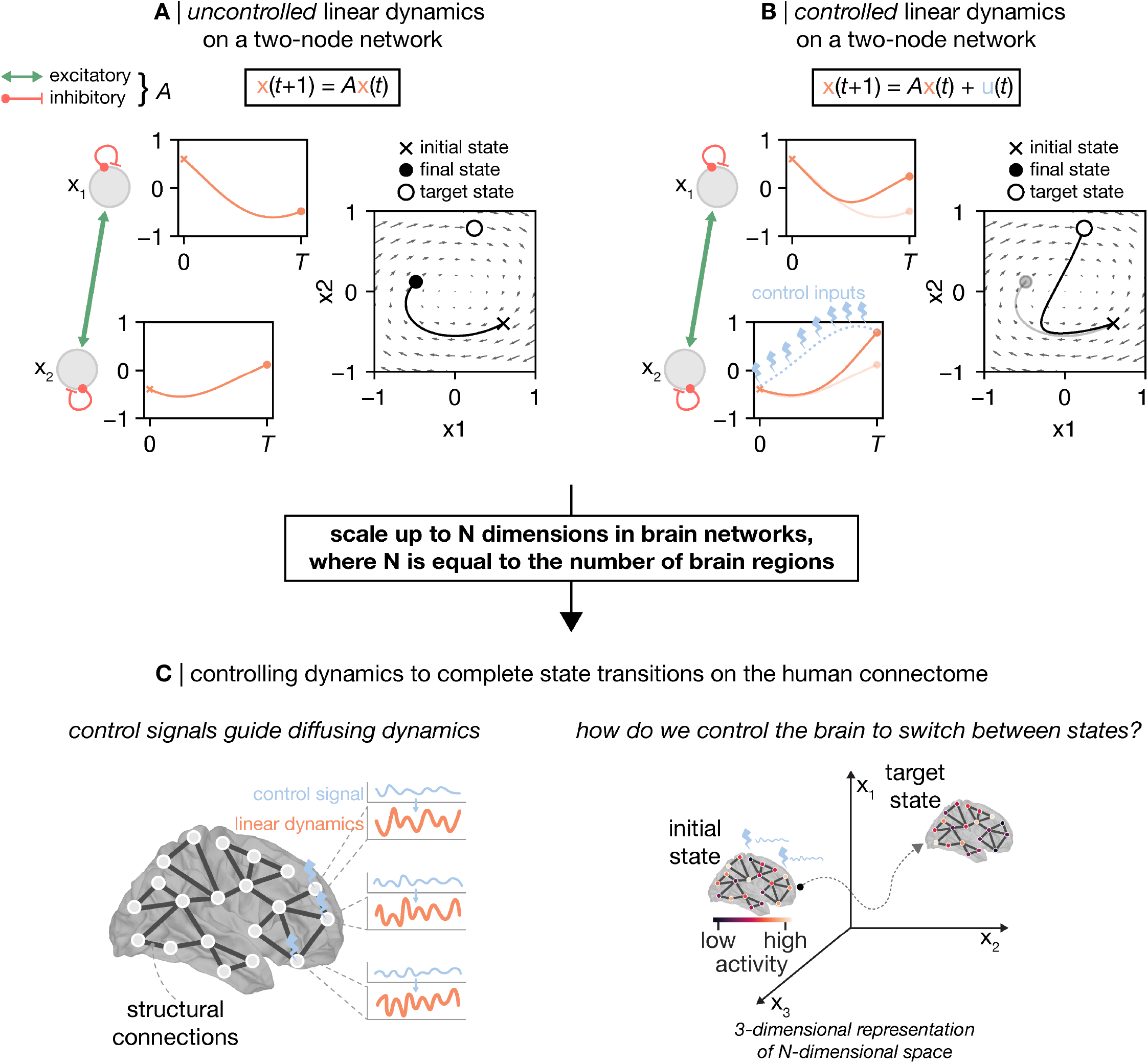
Modeling the *control energy* required to complete a state transition. Network control theory (NCT) finds the control signals that, when injected into a networked system, will guide simulated neural activity from an *initial state* to a *target state*. Here, we show a two-node toy network (*x*_1_, *x*_2_) that illustrates the difference between neural activity (solid orange lines) in the absence (**A**) and presence (**B**) of a control signal (dashed blue line). **A**, Uncontrolled linear dynamics on a two-node network. Given an initial state where *x*_1_ = 0.3 and *x*_2_ = −0.2, as well as coupling between nodes encoded by *A*, uncontrolled neural activity unfolds as shown on the left. These dynamic trajectories can also be represented in two dimensions as a vector field as shown on the right. Under this uncontrolled regime, the state of the system culminates in *x*_1_ = −0.24 and *x*_2_ = 0.06 at time *T*. **B**, Controlled linear dynamics on a two-node network. By contrast, when we introduce a control signal to *x*_2_, the trajectory changes to now culminate in *x*_1_ = 0.12 and *x*_2_ = 0.39 at time *T*. Thus, NCT has found the control signal that drove our system from our *initial state* [*x*_1_ = 0.3, *x*_2_ = −0.2] to our desired *target state* [*x*_1_ = 0.12, *x*_2_ = 0.39]. **C**, NCT applied to the human connectome. The above model can be extended to the scale of *N* brain regions that constitute a whole-brain connectome (left). In doing so, we can model and examine the control signals required to transition the brain between various states of interest (right).

**FIG. 2.**
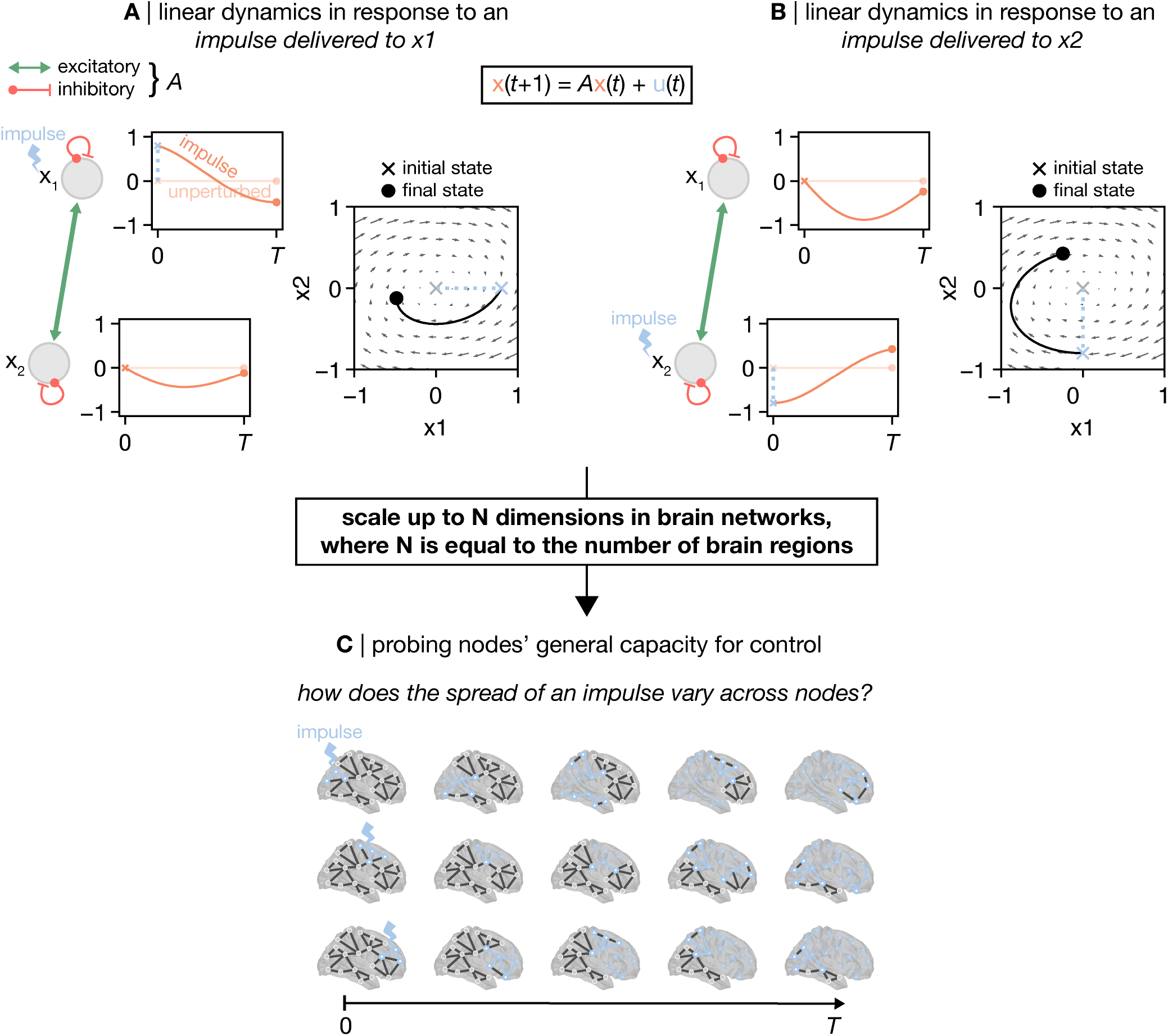
Average controllability: modeling the impulse response of the system from each node. Network control theory (NCT) can be used to probe regions’ general capacity to control dynamics. This is achieved by studying how the system responds to an impulse delivered to each node. Here, we show a two-node toy network (*x*_1_, *x*_2_) coupled by *A*. Upon this network, we demonstrate how neural activity (solid orange lines) unfolds when an impulse (dashed blue line) is delivered to *x*_1_ (**A**) and *x*_2_ (**B**). **A**, An impulse is delivered to *x*_1_ that sets the initial state of the system to [*x*_1_ = 0.4, *x*_2_ = 0]. **B**, An impulse is delivered to *x*_2_ that sets the initial state of the system to [*x*_1_ = 0, *x*_2_ = −0.4]. In each case, the impulse response of the system is quantified as the area under the squared curves of the two orange traces. Intuitively, this measurement corresponds to the amount of activity propagated throughout the system over time. We refer to this measure as the *average controllability*. Thus, for a given time horizon (*T*), a region with higher *average controllability* is better able to broadcast an impulse. **C**, Impulse response modeled for the human connectome. The above model can be extended to the scale of *N* brain regions that constitute a whole-brain connectome. In doing so, we can compare each region’s capacity to broadcast an impulse across the whole brain.

## II. DEVELOPMENT OF THE PROTOCOL

The methods that underlie NCT are based on the established fields of control theory and dynamical systems theory. Dating back to at least the 19th century [33], control theory is primarily concerned with engineering perturbations to achieve desired behaviors in the states of a system, and specifically the evolution of such states over *time*. Hence, one of the most natural ways to formulate theories of control is through *differential* and *difference* equations that mathematically define the next state of a system given its current state. A common example of a control system is an inverted pendulum on a cart: the system states are the positions and velocities of the cart and pendulum, the differential equations are determined through the governing Newtonian physics, and the control task is to perturb the cart so that the cart and pendulum end up in a desired state. For example, one might want to push the cart back and forth in such a way as to stabilize the pendulum so that it remains upright [34].

From one perspective, the inverted pendulum is not unlike the brain, where the system states are the activities of neural units (e.g., brain regions), the differential equations are determined through the diffusion of activity through structural connections between those units, and the control task is to perturb the brain to steer it to a desired state. There is a rich history of such modeling of the brain as a dynamical system using differential equations, ranging from biophysical models of single neurons [35], to phenomenological [36] and coarse-grained [37] models of neural populations. In tandem, there is a very practical translational need to understand how to control brain dynamics [38], for example to compensate for abnormal dynamics that may be present in neurological and neuropsychiatric disorders.

However, there are also many ways in which the brain is not like an inverted pendulum. First is the dimensionality and complexity of the brain. Understanding how the topology of the structural connectome gives rise to brain function is a difficult task that has motivated a large body of work in the last two decades. This research has revealed that structure-function coupling is not one-to-one, varies spatially across the cortex [39–43], and is stronger when indirect structural pathways are accounted for under multiple models of network communication [44, 45]. Second is the distributed nature of brain states for human function. While some brain regions may be thought of as supporting specific functions (e.g., the fusiform face area), carrying out complex human functions typically requires the recruitment of a network of brain regions to a distributed brain state [46]. Finally, biology imposes relatively tight operating constraints. To support complex human function, the brain needs to optimize for efficient signaling while balancing the need to minimize wiring cost within the spatial constraints of the cranial cavity. Hence, there is a need to express the unique complexities and constraints of controlling brain structure-function coupling in the quantitative formalism of dynamics and control.

NCT emerges as a flexible framework that is methodologically based in optimal control theory [47], and can accommodate a wide range of theoretical and experimental hypotheses and constraints about structure-function coupling through a consistent mathematical framework [20, 48, 49]. Because NCT posits a model of neural dynamics at the level of individual neural interactions, it allows us to probe the role of the complex structural connectome on brain function at the level of those interactions [14, 50, 51]. Additionally, because NCT parameterizes which regions to control and how, as well as the precise patterns of initial and target neural activity, it can answer questions ranging from the importance of a single region for propagating activity [10] to the cost of transitioning between specific brain states [11]. Hence, the development of NCT has largely served to provide a simple, first-order biophysical model with the flexibility and power to study more advanced hypotheses of function.

The modeling framework of NCT comprises *N* nodes (e.g., neurons, brain regions) and *m* inputs, and stipulates that the state of each node, *x*_*i*_(*t*), evolves in time as a weighted sum of the state of all upstream nodes, *x*_*j*_(*t*), and any inputs, *u*_*m*_(*t*). If the evolution of the states can be framed in terms of *states*—where the activity of upstream nodes determines the state of downstream nodes at the next *discrete* point in time—then the model takes the form of a difference equation,

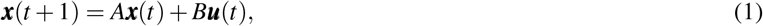

where ***x***(*t*) = [*x*_1_(*t*), *x*_2_(*t*), · · ·, *x*_*n*_(*t*)]^⊤^ is the vector of neural states, *A* is the *N*×*N* connectome, ***u***(*t*) = [*u*_1_(*t*), · · · *u*_2_(*t*),, *u*_*m*_(*t*)]^⊤^ *i*s the vector of independent control signals, and *B* is the *N*×*m* matrix that quantifies how each input affects the nodes. If instead the evolution of the states can be framed in terms of *rates*—where the activity of upstream nodes affects the *continuous* rate at which the state of downstream nodes change—then the model takes the form of the differential equation

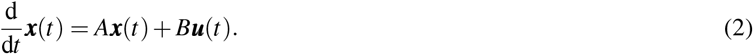

While these two models appear similar at first glance, their definition, properties, and behavior differ substantially. In turn, the interpretation of the model parameters and outputs can vary dramatically between them.

This protocol will discuss two common operationalizations of NCT that can be derived from either of these models. The first employs a time-varying perturbation, ***u***(*t*), to drive the neural activity, ***x***(*t*), from an initial state, *x*0, to a target state, *xf*, given a balance of constraints on the magnitude of both the neural states and the perturbations. The magnitude of these perturbations are summarized as the *control energy*, which we interpret as the amount of effort that the model system must exert to complete a given state transition. The second is *average controllability*, which is the magnitude of the neural activity, ***x***(*t*), in response to an impulse stimulus delivered to a single node; a node with higher *average controllability* is better able to leverage graph topology to spread an impulse throughout the system. Note, *average controllability* is only one example of a node-level NCT metric that falls within the broader category of *controllability statistics*. This category encompasses NCT outputs that describe different ways in which the nodes of the system may control its dynamics. While we have used other controllability statistics in our previous work (e.g., *modal controllability*, see section III), we focus on *average controllability* in this protocol owing to its simple intuition and broad appeal.

We focus on these two operationalizations—*control energy* and *average controllability*—because they encompass two common sets of questions about the brain. The first set of questions stems from advances in neuroimaging that allow us to empirically measure neural states in many forms such as functional Magnetic Resonance Imaging (fMRI), electrophysiology, and calcium imaging [52]. Given these state-level empirical data, a natural question is “how does the brain reach or switch between these states using regimes of internal or external control?” Optimal control theory provides a powerful and flexible set of tools to explore these questions under various constraints and at different spatio-temporal scales. The second set of questions stems from empirical evidence demonstrating that individual and groups of brain regions may be important for (i) enabling specific functions, such as visual processing [53], motor processing [54], and cognition [55, 56]; (ii) supporting critical functional processes in the brain, such as segregation and integration [57, 58]; and (iii) may be disproportionately impacted by diseases processes [59, 60]. Given these data, a natural question to ask is “what is the contribution of these sets of regions to the control of brain activity?” Average controllability measures the magnitude of propagation of stimulation along neural tracts, thereby providing a coarse-grained understanding of this contribution.

## III. APPLICATIONS OF THE METHOD

Analyzing brain data using a network representation is increasingly popular in neuroscience, and researchers have used a wide range of connectomic data to perform NCT analysis. For example, the availability of multimodal neuroimaging data in large cohorts accompanied by clinical and cognitive data [21, 22, 61–63]—as well as indices of neurobiology not measurable *in vivo* (e.g., high-resolution histology [64], gene expression [65], etc.)—enable researchers to validate NCT against brain function and biology as well as examine individual differences. Indeed, we have applied NCT in our research with a view towards achieving these goals. Here, we briefly review selections of this work to show how our protocol may be applied to study the brain. Specifically, we discuss how model outputs from NCT (A) link to network topology; (B) explain individual differences in mental health symptoms, cognition, and age; (C) predict effects of neurostimulation; (D) explain switching between functional task states; and (E) link to neuroanatomy.

### A. Understanding the Influence of Topology

In our early work, we began by contextualizing nodal controllability statistics against what we know about connectome topology from graph theory. Specifically, Gu *et al*. [10] examined how nodal control properties–specifically *average controllability* (see Pathway B, Section IX B) and *modal controllability*–correlated with nodes’ strength (the sum of a node’s edge weights). Gu *et al*. [10] found that nodes’ strength correlated strongly positively and negatively with average and modal controllability, respectively. These relations were conserved across both humans and macaques. Collectively, these results indicate that a node’s local topological importance predicts its capacity to control the dynamics of a system.

We have also examined how connectome topology influences the *control energy* associated with specific state transitions. Betzel *et al*. [49] found that nodes’ topological importance predicted their capacity to facilitate transitions between eight canonical brain states (seven resting-state cortical networks as well as a subcortical network [66]). Specifically, Betzel *et al*. found that target states that intersected with the brain’s rich club [58]—a set of highly interconnected nodes that form the connectome’s core—exhibited low transition energy. This result demonstrates that the rich club is well positioned in the network to act as an efficient target state to which a diverse set of initial states can transition with low *control energy*. Thus, the topology of the human connectome may be optimized to guide dynamics toward the rich club, bolstering the idea that these nodes support functional integration [57, 67, 68].

Given these advances in understanding how connectome topology contributes to control, we subsequently analyzed what the underlying control equations could tell us about network topology. Starting from the NCT equations, Kim *et al*. [50] derived the features of network architecture that were the most important to determining *control energy*. Kim *et al*. [50] discovered that a strong and diverse set of connections from stimulated nodes to unstimulated nodes were the leading-order contributors to the control cost. Using this discovery, the authors reduced the cost of controlling connectomes in the *Drosophila*, mouse, and human by virtually resecting edges, and developed a method to meaningfully compare the control cost between different species and connectomes. These results provide simple and quantitative knowledge about the most important features of topology according to NCT.

### B. Individual Differences

While the strong correlation between *average controllability* and strength reported by Gu *et al*. [10] may seem to imply redundancy between nodal controllability statistics and measures from graph theory incorporating weighted degree, we note that this correlation was spatial (i.e., across the brain) rather than between subjects. In subsequent work examining individual differences, Parkes *et al*. [69] compared the capacity of *average controllability* and strength to predict psychosis spectrum symptoms using out-of-sample testing. Parkes *et al*. [69] found that *average controllability* significantly outperformed strength in this predictive task, and demonstrated that this improved performance was concentrated in higher-order default mode cortex [70]. These results show that while high *average controllability* may depend upon high strength, there exists unique inter-individual variation between the metrics, and that this variance in *average controllability* couples more tightly to mental health symptoms.

We have also shown that *average controllability* exhibits robust developmental and sex effects. *Average controllability* increases between the ages of 8 and 22 years [12] and is higher in females in the cortex while higher in males in subcortex [19]. Furthermore, Tang *et al*. [12] showed that age effects were strongest in nodes with higher controllability, underscoring the developmental importance of nodes that are well positioned in the network to control dynamics. When examining *control energy*, Cui *et al*. [71] demonstrated that the amount of energy required to activate the fronto-parietal system—a brain network thought to support executive function [72]—was negatively correlated with both age and executive function in the same sample. This result suggests that the developmental emergence of executive function is associated with increased efficiency of neural signaling within the human connectome.

### C. Predicting Stimulation Effects

An application of NCT that has clear translational impact is modeling the relationship between brain structure and function. To this end, we have examined whether NCT can predict the brain’s functional response to neurostimulation from its structural connectome. For example, in patients with epilepsy, Stiso *et al*. [14] found that NCT was able to predict electrophysiological neuronal responses (measured with electrocorticography) following direct electrical stimulation. This result shows that our model, wherein neural activity is simulated upon the structural connectome, explains variance in experimentally-manipulated empirical changes in brain state.

We have also examined NCT in the context of non-invasive neurostimulation techniques. In a pair of studies, Medaglia *et al*. [16, 17] delivered transcranial magnetic stimulation (TMS) to the left inferior frontal gyrus (LIFG) in between repeated sessions of a set of language tasks. Across both studies, the authors found that NCT metrics extracted from the LIFG explained variance in changes to task performance before and after TMS. These results demonstrate that NCT can be used to probe the network mechanisms that underpin how neurostimulation elicits changes in behavior.

### D. Modeling Switches Between Functional Brain States

In addition to predicting the effects of neurostimulation, NCT can be used to investigate how the topology of the structural connectome supports transitions between empirically-observed functional brain states. Our group has studied this process using brain states derived from fMRI. Cornblath *et al*. [20] clustered resting-state fMRI (rs-fMRI) into brain states representing instantaneous co-activations among canonical brain networks, and used NCT to model the energy required to transition between those states. Using a series of null network models (see section X), Cornblath *et al*. found that the topology of the structural connectome was wired to support efficient switching between brain states. This result demonstrates that the topology of the connectome is optimized to support dynamic fluctuations in resting-state activity.

Subsequent work by Braun *et al*. [73] examined transitions between brain states elicited by a working memory task. Braun *et al*. [73] found that transitioning from a 0-back brain state to the more cognitively demanding 2-back brain state required more energy than the reverse transition, demonstrating an asymmetry in *control energy*. Braun *et al*. also found that this energy asymmetry was more pronounced in patients with schizophrenia compared to healthy controls. Thus, while connectome topology may be setup to enable low-cost fluctuations in resting-state [20], activating cognitively demanding brain states may require more control effort. Furthermore, this increased control effort appears to scale with within-task differences in cognitive demand and is further elevated in psychopathology.

### E. Biologically informed NCT

Neuroscience is increasingly moving toward a multi-scale approach that seeks to understand how features of the brain observed at one scale link to properties observed at another, and *vice versa* [3, 74–82]. Recently, we have applied this multi-scale approach to NCT by examining how dynamics within the model are influenced by variations to regions’ cellular composition. Specifically, we [51] examined how regions’ profiles of cytoarchitecture impacted the energy associated with state transitions that spanned the cortical hierarchy (i.e., the sensory-fugal axis [83]). We found that state transitions traversing bottom-up along the cortical hierarchy of cytoarchitecture required lower *control energy* to complete compared to their top-down counterparts, and observed that nodes’ position along this hierarchy predicted their importance in facilitating these transitions. This result shows that spatial variations in cortical microstructure constrains macroscopic connectome topology; this effect is consistent with work from neuroanatomy that describes a precise relationship between regions’ profiles of cytoarchitecture and their extrinsic connectivity [84].

In recent work from outside our group, Luppi *et al*. [32] characterized the *control energy* associated with a large set of activity maps derived from NeuroSynth [85] related to cognition. Additionally, the authors examined how these transition energies varied when they utilized a broad range of neurotransmitter density maps to modify the control weights. This work ties together switching between functional brain states and biologically informed connectome analysis to provide the field with a comprehensive “look-up table” of how regions’ diverse biology impacts *control energy*.

## IV. COMPARISON WITH OTHER METHODS

We consider NCT with respect to other models that also seek to understand how communication unfolds within a structurally interconnected complex system. Specifically, for practicing neuroscientists, we view NCT as complementing both more complex biophysical models of dynamics as well as graph-theoretic measures of inter-nodal communication. While both of these approaches model communication, they differ in their biological plausability and complexity.

Biophysical models aim to capture neuronal communication by distilling the various biophysical processes necessary for functional activity into separate model parameters. These parameters are tuned to simulate biologically plausible non-linear dynamics within and between neurons at multiple scales. For example, at the scale of single neurons, the Hodgkin-Huxley model is concerned with modeling neuronal spiking activity [86], and is based on parameterizing the flow of sodium and potassium ions across the cell membrane. At the next scale up, mean-field models focus on the collective activity patterns of co-located populations of neurons [87, 88]. Coupling multiple mean-field models together—where each model represents distinct neuronal populations—enables researchers to study how non-linear dynamics emerge from brain structure at the macroscale [87]. In turn, this approach gives rises to a wide range of complex dynamical behaviors, including synchronized oscillators [89], learning [90, 91], large-scale traveling brain waves [92], and structure-function coupling [93–95]. Broadly, NCT trades biophysical accuracy and the complexity of specific model behaviors for more power in designing and studying stimuli. For example, in lieu of studying state transitions that emerge from different models of associative memory [96] and context integration [97], NCT allows us to design specific stimuli to transition the model system to states that are known to be important for memory and cognitive control under specific constraints [49].

By contrast, graph-theoretic approaches instantiate relatively simple models of inter-nodal communication that rely on assumptions such as shortest-path routing, spatial proximity, random walks, and diffusion processes [4, 9]. While these assumptions are an oversimplification of brain dynamics—and are thus less biologically plausible—their simplicity confers greater analytic tractability and scalability, which are both desirable features when studying the human brain. This benefit compounds when the goal of a given study is to examine inter-individual differences, wherein dynamical models may be fit to thousands of participants. As such, despite their relative simplicity, graph-theoretic approaches have deepened our insights into large-scale brain organization [55, 56, 98–101], improved our understanding of the link between the brain and mental disorders [59, 102–104], and helped elucidate the link between structure and function [40, 44, 45, 105–107].

We consider NCT as situated between these two modeling approaches. As discussed in Section II, NCT is essentially a model of two parts, *dynamics* and *control*. For the former component, NCT models dynamics according to a diffusion process; thus, like graph theory, NCT makes simplifying assumptions of inter-nodal communication, which confers the advantages of analytic tractability and scalability. However, the second component, *control*, adds an additional layer of model parameterization that allows researchers to probe how the system might behave under different contexts (e.g., in response to task manipulations, cognitive control, or neurostimulation protocols). This added flexibility brings NCT closer to biophysical modeling, insofar as they both seek to understand how the dynamics of a system respond to external perturbation. Indeed, we have shown that NCT can be used to predict changes in the dynamics of coupled Wilson-Cowan oscillators following simulated stimulation [18], suggesting that NCT can explain some of the behaviors engendered by non-linear biophysical models.

## V. EXPERIMENTAL DESIGN

The goal of an NCT analysis, as it is conceptualized in this protocol, is to understand how the topology of the structural connectome supports and constrains spreading dynamics, and to what extent those dynamics can be controlled. Thus, core to this analysis is the acquisition of one or more structural connectomes from a model organism. These connectomes can be obtained in numerous ways that each depend on the model organism under study and the available imaging modality. In humans, for example, structural connectomes are typically extracted from diffusion-weighted imaging (DWI) sequences obtained using MRI. Tractography algorithms are applied to DWI scans to model the white matter pathways intersecting pairs of brain regions, which are then used to populate connectome edges with the number of those pathways (e.g., the streamline count) [4]. This example constitutes a weighted undirected connectome upon which NCT can be conducted. Note that while the application of NCT is not restricted to weighted undirected connectomes (edges can be weighted or unweighted, directed or undirected), the edge weight definition determines what types of questions can be addressed using this protocol (for example, see Ref. [48] for analysis of an unweighted directed *Caenorhabditis elegans* connectome).

Given that connectomes are central to the application of NCT, any artifacts present in the connectomes will be reflected in model outputs. For example, connectomes populated by DWI estimates of connectivity are known to contain false positives and false negatives, which may be partly mitigated by the use of thresholding techniques [108, 109]. In-scanner head motion is well known to spuriously impact these estimates of connectivity as well [110, 111]. Thus, the accurate generation and rigorous quality control of connectomes are both crucial considerations for experimental design. For human connectomes, we recommend researchers consult the extant literature on the processing and quality control of DWI scans [108, 112, 113] (see also https://qsiprep.readthedocs.io/).

Another consideration for connectome estimation is the brain parcellation used to define system nodes. If, as mentioned above, structural connectivity is determined by streamline count, then variations in the size of regions across the parcellation will bias connectome edge weights; larger brain regions will intersect with more white matter pathways and thus show higher connectivity to the remaining regions. As with any analysis of graph topology, this bias will effect the outputs of NCT; for example, larger regions may show higher *average controllability* just by virtue of being more directly connected to the system. It is for this reason that we recommend researchers reproduce their results using several different parcellation definitions and resolutions. Doing so ensures that their results are not driven by a specific parcellation choice.

Beyond the core requirement of a connectome, the flexibility of NCT makes it applicable to a broad range of experimental designs (see Section III); the most critical component is that researchers have hypotheses that pertain to studying the control of brain dynamics. However, in the case of *control energy*, where researchers will study the control signals, ***u***(*t*), there are some additional considerations. First, differences in brain states’ magnitude will impact *control energy*, potentially necessitating the normalization of state magnitude. For example, if researchers are examining transitions between patterns of brain activity (e.g., using functional data as in [20, 73]), then differences between states’ mean activity will impact *control energy*; assuming a common initial state, target states with higher activity will require more energy to transition to compared to target states with lower activity. This effect generalizes to binary brain states—where a brain state is encoded as a set of nodes being “on” while the rest of the nodes are “off”—as well. In this case, differences in state size (i.e., the number of “on” regions) constitute differences in state magnitude; transitioning to larger target states will require more energy. If there are differences in state magnitude, we recommend normalizing states before computing *control energy* (see Section IX A, step 3). Note, the need for this normalization will depend upon researchers’ analyses. For example, if researchers are studying individual differences in the energy associated with a single transition, then normalization may not be necessary so long as state definition is consistent across subjects. What is critical is that researchers consider what comparisons they want to make and whether variations in state definition would confound those comparisons.

A second consideration is how researchers define their control set, *B*. As discussed in Section II, the *N*×*m* control set defines the extent to which the nodes of the system can affect changes in its dynamics. In turn, the definition of *B* determines the dimensions of ***u***(*t*); the greater the number of control nodes, the more independent control signals will be generated. In our work, we have often deployed a *uniform full control set*, which means that all of the nodes of the system are designated as controllers (*full*) and all are given equivalent control over dynamics (*uniform*). In this case, *m* = *N*. Intuitively, this approach assumes that the entire brain is being controlled—either internally or externally—when completing a state transition. However, depending on a researcher’s hypotheses, this assumption may not be appropriate. Instead, researchers may want to define only a subset of nodes as controllers (e.g., [49, 50]), or assign variable weights to control nodes (e.g., [30–32, 51]), or both. Note that assigning variable control weights serves to give some nodes more control over system dynamics than others. In any case, it is critical that researchers check whether their designated control set was able to complete the associated state transition (see Section IX A, step 5); successful completion of a state transition is not guaranteed in the model, and completion is less likely when transitions are driven by a small control set.

## VI. EXPERTISE NEEDED TO IMPLEMENT THE PROTOCOL

We provide open-source and broadly accessible tools that implement *optimal control* and *average controllability* in a Python-based software package called *network control theory for python (nctpy)*. In *nctpy*, we provide a flexible implementation that enables researchers to make model assumptions that best fit their research question. As a result, while a full understanding of linear systems and optimal control theory are not required, the researcher must have enough expertise to make key modeling decisions that best represent the data, which we explicitly mark in the protocol.

The first piece of expertise needed is to understand the differences between (and implications of) discrete-time systems and continuous-time systems (see Section II). This difference is not merely a conceptual one, because discrete-time systems display a fundamentally different set of behaviors than continuous-time systems. That is, a discrete-time system is not simply a temporally coarse-grained version of a continuous-time system. Instead, each system exhibits different dynamics. As a simple example, consider a 1-dimensional, discrete-time system that evolves according to *x*(*t* + 1) = −*x*(*t*). Starting at 1, this system will alternate between -1 and 1 over time. There does not exist an equivalent continuous-time system that evolves according to 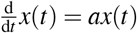 (where *a* is a real number) that oscillates in this way.

The second piece of expertise needed is to understand the nature of Pathway A (*control energy*, Section IX A) and Pathway B (*average controllability*, Section IX B) in order to interpret the outputs. Pathway A is solving an optimization problem. Specifically, first we provide a model of the dynamics (i.e. *A, B*), the initial and target states, and some optimization parameters (see Section IX A). Then, we solve for the control signals, ***u***(*t*), that minimize the cost. Hence, all interpretations of ***u***(*t*) should be made with the understanding that they were determined by the user-defined optimization parameters. Pathway B is not solving an optimization problem, and thus does not receive any optimization parameters. Rather, it measures the magnitude of the neural states over time as a result of an impulse stimulation. Because Pathways A and B use the same dynamics but output different quantities through different means, more expertise in linear systems and optimal control is needed to meaningfully compare and contrast the two pathways.

## VII. LIMITATIONS AND ONGOING DEVELOPMENT

Network control theory has the ability to flexibly accommodate many scientific questions and to generate concise knowledge from a simple model. However, NCT also possesses several limitations that are faced by many in the study of high-dimensional complex systems, such as numerical stability of algorithms, validation at the scale of microscopic states, approximations of complex interactions, and interpretation of model parameters.

### A. Numerical Stability of Optimization

One limitation is the numerical stability of Pathway A under certain parameter conditions, which arises from ill-conditioned matrices that are built while solving for the control signals. This issue occurs most frequently when using a relatively small control set—a small *m* in the *N*×*m* matrix *B*—to control a network with large *N*. It is intuitive that precisely controlling the initial and target states of the whole brain from only a few nodes is difficult. In light of this limitation, it is crucial that the researcher carefully study the generated trajectories of the neural activity to ensure that the desired initial and target states are reached, and that the numerical integrator does not generate a warning of numerically ill-conditioned matrices. In the event that the control set must be small for the purposes of the research question, one solution may be to extend the control set by heavily weighting the desired control nodes and lightly weighting the remaining nodes [14]. Another option is to use Pathway B to study the *average controllability* of the control set.

### B. Validation at the Scale of Microscopic States

A second limitation is the validation of the model at the level of individual neural states. Phrased another way: given a connectome, *A*, and stimulations ***u***(*t*) delivered to brain regions *B* starting at neural activations ***x***(0), does experimentally measured neural activity agree with the simulated trajectory ***x***(*t*)? The challenges associated with addressing this sort of question extend far beyond NCT and to a significant portion of systems and network neuroscience. Microstate validation between neural structure and activity is most evident in small systems of neural circuits [114], but how to perform similar validations for large-scale systems such as the human brain remains an open area of research. Challenges include (i) the multiple possible scales of constructing brain networks [2, 115]; (ii) differing measures of inter-areal connectivity [108, 116]; (iii) multiple definitions of simulated neural activity [87, 117–119]; and (iv) the diverse spatial and temporal resolutions at which we can record whole-brain activity [52]. Along this active area of research, recent work has demonstrated that linear models outperform non-linear and kernel-based models in both 1-step prediction and model complexity for both fMRI and EEG data [120], as well as correspondence between *control energy* and local metabolism [121].

Closely linked to ideas of validation is ongoing work on the implementation of the connectome, *A*. NCT stipulates a simple but mechanistic model for the evolution of neural states, ***x***(*t*), given a stimulus ***u***(*t*). Hence, the interpretation of the model outputs is bounded by the interpretability of the model inputs. In the majority of applications, the matrix *A* is taken to be the structural connectome, with the justification that regions must be able to transmit information along structural connections. However, the distribution of edge weights can vary significantly across different pre-processing pipelines, which implies equally significant variations in the strength of interactions between regions. Hence, it is important for the interpretation of the results to critically assess whether the edge from node *i* to node *j* of the implemented adjacency matrix is a reasonable measure of the strength of activity diffusion from node *i* to node *j*.

### C. Linear Dynamics

A third limitation is the assumption of linear dynamics, which enables the calculation of powerful measures such as optimal control trajectories, but hinders the biophysical realism of the framework. More sophisticated non-linear models capture complex dynamics from individual neurons [86] to neural populations [37], thereby enabling the study of fine-grained experimental behavior [122] and complex non-linear phenomena [123]. These models make fewer simplifying assumptions to capture non-linear behaviors of biological systems such as complex memory landscapes [124]. While prior work has shown that linear models outperform many classes of non-linear models in describing and predicting brain-wide neural activity [120], extensions of NCT to non-linear systems will enable greater flexibility to accommodate and explore the impact of non-linear biophysical constraints. While the theory of non-linear control is an active area of research [125], there are immediate applications of NCT to non-linear systems, and many exciting potential extensions of NCT to capture more biophysical realism.

Broadly speaking, the linear dynamics assumed by NCT can be thought of as being valid for a non-linear system within small deviations of an operating state [7]. Hence, the most immediate application of NCT to non-linear systems is to linearize the model about an operating point, such as the upright position of an inverted pendulum [126]. Along these lines, the next immediate generalization to NCT is to linear time-varying systems [47], where the model is linearized not about a point, but about a trajectory. While methods to implement control for linearized and time-varying systems are well-established in the control community, a biophysically meaningful implementation and interpretation of the parameters—namely *A*(*t*) and *B*(*t*)— remains an area of active work [127]. Another approximation that is particularly relevant for high-dimensional neural systems is at the limit of weakly coupled oscillators [128, 129], whereby a high-dimensional system of oscillators with weak interactions can be reduced to a low-dimensional phase response curve, allowing for the potential linearization of the system about phase-locked states.

In addition to linearizing dynamics about points and trajectories, NCT can also meaningfully be applied to non-linear dynamical systems that can be made linear through a non-linear change of variables. One such example is by using finite-dimensional *Koopman subspaces*, which allow for the recasting of non-linear systems with single fixed points as higher-dimensional linear systems [130], and closely-related methods in dynamic mode decomposition [131]. Further, advances in non-linear control enable us to probe important coarse-grained questions such as the control set necessary to push non-linear systems between attracting states [132]. Other control strategies take advantage of the ability of non-linear systems to access states that lie outside of their linearization [133].

## VIII. MATERIALS

As discussed above, we split our protocol into two pathways (Figure 3). The primary pathway of our protocol focuses on computing *control energy* (Pathway A, Section IX A). This pathway is illustrated in Figures 3A, 3B, and 3C. Briefly, Figure 3A outlines the inputs required to run our protocol, Figure 3B outlines the model outputs, and Figure 3C outlines some of the variations to model inputs that we have discussed thus far. The second pathway of our protocol focuses on computing nodes’ *average controllability* (Pathway B, Section IX B; Figure 3D).

**FIG. 3.**
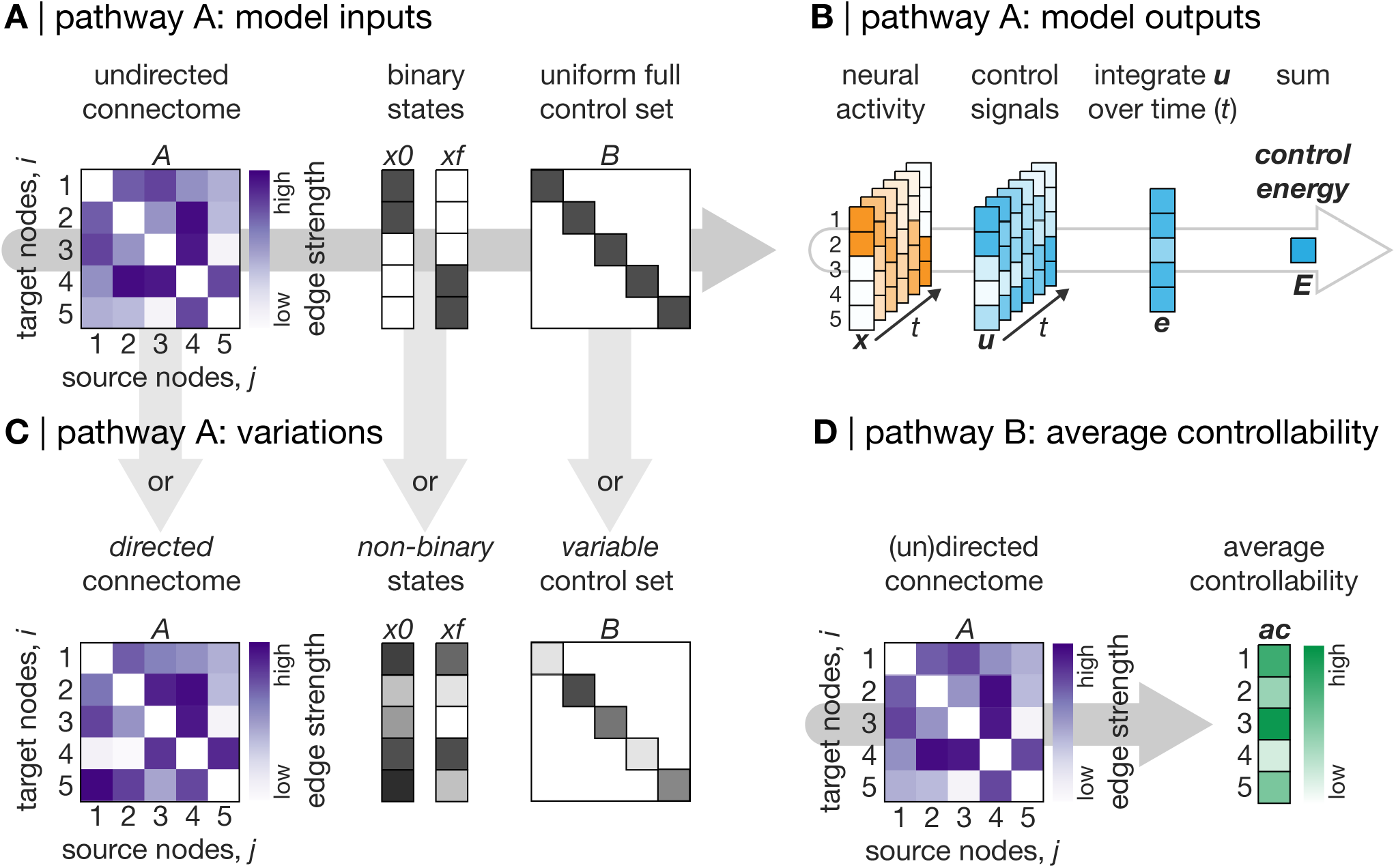
Schematic representation of the network control theory (NCT) protocol. Our protocol is split into two pathways. Primarily, our protocol focuses on modeling the *control energy* associated with user-defined control tasks. We refer to this part of our protocol as Pathway A (**A, B**, and **C**). Pathway A will be of interest to researchers who seek to study specific state transitions. We also outline a brief protocol for estimating nodes’ *average controllability*. We refer to this part of our protocol as Pathway B (**D**). Pathway B will be of interest to researchers who want to examine nodes’ general capacity to control system dynamics. **A**, Inputs required for Pathway A. To compute *control energy*, researchers must provide a structural connectome (*A*), an initial state (*x*0), and a target state (*xf*), and must also define a control set (*B*). **B**, Model outputs from Pathway A. Given these inputs, our protocol will output the state trajectory (neural activity, ***x***(*t*)) and the control signals (***u***(*t*)). Once inspected, the control signals can be integrated over time to obtain node-level energy (***e***), which in turn can be summed over nodes to get the *control energy* (***E***). **C**, Variations to Pathway A. Pathway A can handle a diverse range of inputs, including but not limited to undirected and directed connectomes (left), binary and non-binary brain states (middle), and control sets with uniform or variable weights (right). **D**, Pathway B: *average controllability*. Pathway B only requires a structural connectome (*A*) as input and will return the *average controllability* of each node. This metric quantifies the impulse response of the system from a given node. Higher *average controllability* indicates that a node is better positioned in the network to propagate dynamics.

### Equipment

- A computer with *Python* (tested on version 3.9) and `nctpy` installed alongside its dependencies. This protocol has been tested on Mac OS running on Intel Core i5/i7/i9 processors as well as on Apple Silicon. We have also tested this protocol on Linux Ubuntu running on Intel processors. RAM requirements will vary depending on researchers’ data and analyses, but we recommend at least 16 GB. Finally, we recommend installing `nctpy` inside a virtual environment managed by Anaconda (https://www.anaconda.com/). The following core dependencies are required to run `nctpy`:
  - numpy (https://numpy.org/). Tested on version 1.24.3.
  - scipy (https://scipy.org/). Tested on version 1.10.1.
  - tqdm (https://github.com/tqdm/tqdm). Tested on version 4.65.
  Additionally, there are some functions in `nctpy.plotting` and `nctpy.utils` that require the following:
  - statsmodels (https://www.statsmodels.org/). Tested on version 0.13.5.
  - matplotlib (https://matplotlib.org/). Tested on version 3.7.1.
  - seaborn (https://seaborn.pydata.org/). Tested on version 0.12.2.
  - nibabel (https://nipy.org/nibabel/). Tested on version 5.1.
  - nilearn (https://nilearn.github.io/). Tested on version 0.10.1.
  Finally, the following optional packages were used to run the analyses illustrated in this protocols paper:
  - (optional) pandas (https://pandas.pydata.org/). Tested on version 1.5.3.
  - (optional) scikit-learn (https://scikit-learn.org/). Tested on version 1.2.2.
  See https://github.com/BassettLab/nctpy for more details. Creating a *Python* environment using Anaconda and installing the above dependencies should take no longer than 30 minutes.

### Input Data

- CRITICAL **Adjacency matrix**, *A* **(used in all steps)**: The adjacency matrix, *A* (Figure 3A, left), is the primary input to our protocol. In *A*, the *N* nodes of the system are stored on the rows and columns, and the *N*×*N* edge values are stored in the entries. As discussed above, both the nodes and the edges of *A* can be defined in numerous ways and their definition depends on the acquired data as well as the research question. For example, the nodes of the system may be defined as single neurons in organisms such as the *Caenorhabditis elegans* [48, 134] or as brain regions of varying size and definition in organisms such as the mouse [76], Drosophila [50], macaque [135] and human [4]. The edges of *A* may be defined as either the directed or undirected structural connectivity between nodes (Figure 3C). Note, effective functional connectivity between nodes may also be used to define edges [15, 136, 137]. Critically, our model assumes that *A*_*ij*_ encodes the strength of diffusion of activity along the edge connecting *node j to node i*. In other words, our model assumes that the columns of *A* store the source nodes (i.e., projections from node *j*) while the rows store the target nodes (i.e., projections to node *i*). While this distinction is irrelevant for undirected connectomes where *A*_*ij*_ = *A*_*ji*_, it is crucial for directed connectomes. Thus, researchers must ensure that their directed *A* matrix conforms to the above assumptions. In any case, NCT can be applied to either participant-level or group-averaged connectomes. Although we focus on a group-averaged undirected structural connectome in this protocol, we also include an example of our protocol applied to a directed structural connectome.
- **Brain states**, *x*0 **and** *xf* **(used in steps 3-6)**: In order to analyze state transitions, researchers also need to provide a pair of brain states (Figure 3A, middle; initial state, *x*0; target state, *xf*) relevant to their hypotheses. Providing these states allows NCT to find the control signals, ***u***(*t*), that are required to transition between them and to summarize those control signals as *control energy* (Figure 3B); here, *control energy* estimates the amount of effort the model has to exert to complete the transition. Brain states can be defined in a number of ways. The simplest approach is to define each brain state as a binary vector (Figure 3A, middle), where nodes that are within a given state are assigned an arbitrary constant value (e.g., 1) and any remaining nodes are assigned a value of 0. In this setup, NCT is tasked with transitioning the brain between actuating different sets of node to a constant arbitrary level of neural activity. We commonly adopt this approach in our work. An alternative approach is to allow brain states to represent a variable pattern of activity (Figure 3C, middle). As mentioned above, Cornblath *et al*. [20] modeled the energy required to transition between brain states derived from clustering of fMRI data, while Braun *et al*. [73] used task activation maps extracted from an fMRI contrast. These approaches allowed the authors to generate state vectors that encode non-zero activity across all nodes of the system. In this protocol, we illustrate examples using both binary and non-binary brain states. Note that brain states are not required for Pathway B (Figure 3D).
- **Control set**, *B* **(used in steps 3-6)**: In addition to brain states, researchers also need to designate a control set; these are the nodes that NCT will use to complete state transitions. Intuitively, the control nodes are where time-varying control signals, ***u***(*t*), will be injected into the system in order to transition the system between states. Each column *k* of *B* indicates the impact of input *u*_*k*_ on the network nodes. Here, the simplest approach is to define a *uniform full control set* (Figure 3A, right), which means that all nodes of the system are designated as control nodes and that they are all given the same degree of control over system dynamics. Alternatively, as discussed above, researchers may also designate partial control sets or control sets with variable weights (Figure 3C, right). For the latter, variable weights can either be derived *a priori* and input directly into the model (e.g., [30–32]) or via data-driven optimization approaches (e.g., [51]). We illustrate examples of using such control sets in this protocol. Note that a control set is not required for Pathway B (Figure 3D).

### Example Dataset

- We primarily used undirected structural connectomes derived from DWI performed on the human brain. We obtained these connectomes from the Philadelphia Neurodevelopmental Cohort (PNC) [21, 22], a community-based study of brain development in youths aged 8 to 22 years. The neuroimaging sample of the PNC consists of 1,601 participants. From this original sample, we retained 253 typically developing participants who had no medical co-morbidity or radiological abnormalities, and who were not taking psychoactive medications at the time of assessment. Additionally, these participants’ T1-weighted, DWI, and rs-fMRI scans all passed stringent quality control procedures [113, 138, 139].
- Structural connectome reconstruction was performed using QSIprep 0.14.2 [112], which is based on Nipype 1.6.1 [140]. Connectomes were extracted using the 200-node variant of the Schaefer parcellation [115], ordered according to 7 canonical brain systems [66]. The strength of inter-regional connectivity was summarized using the number of streamlines that intersected each pair of parcels. Connectomes were averaged over subjects. This group-averaged connectome was thresholded by retaining the edges that were present in at least 60% of participants’ connectomes [109]. This process resulted in a final connectome with 98% edge density.
  - rs-fMRI was also obtained from the same 253 PNC participants [21]. These data were used to generate empirical brain activity states (see Figure 6). The eXtensible Connectivity Pipeline (XCP-D) [139, 141] was used to post-process the outputs of fMRIPrep version 20.2.3 [142]. XCP was built with Nipype 1.7.0 [140]. Processed rs-fMRI time series were extracted from the same 200-node parcellation mentioned above [115].
- We also studied a directed structural connectome obtained from the Allen Mouse Brain Connectivity Atlas.
  - Whole-brain structural connectomes were constructed with 2 ×10^5^ voxels at a spatial resolution of 100 µm. Voxels were assigned to regions (coarse structures) according to a 3-D Allen Mouse Brain Reference Atlas [23]. Isocortex was further divided into 6 systems (auditory, lateral, medial, prefrontal, somatomotor, and visual) based on prior work that applied community detection to identify stable modules [143]. Connection strengths were modeled for all source and target voxels using data from 428 anterograde tracing experiments in wild type C57BL/6J mice [144]. Normalized connection strengths were obtained by dividing the connection strengths by the source and target region sizes. Here, we retained only the 43 isocortical regions. This process resulted in a fully-connected directed structural connectome.

In all of the below code, we assume the existence of a *Python* environment with `nctpy` installed alongside its dependencies. First, we import all the functions we need to run our protocol:

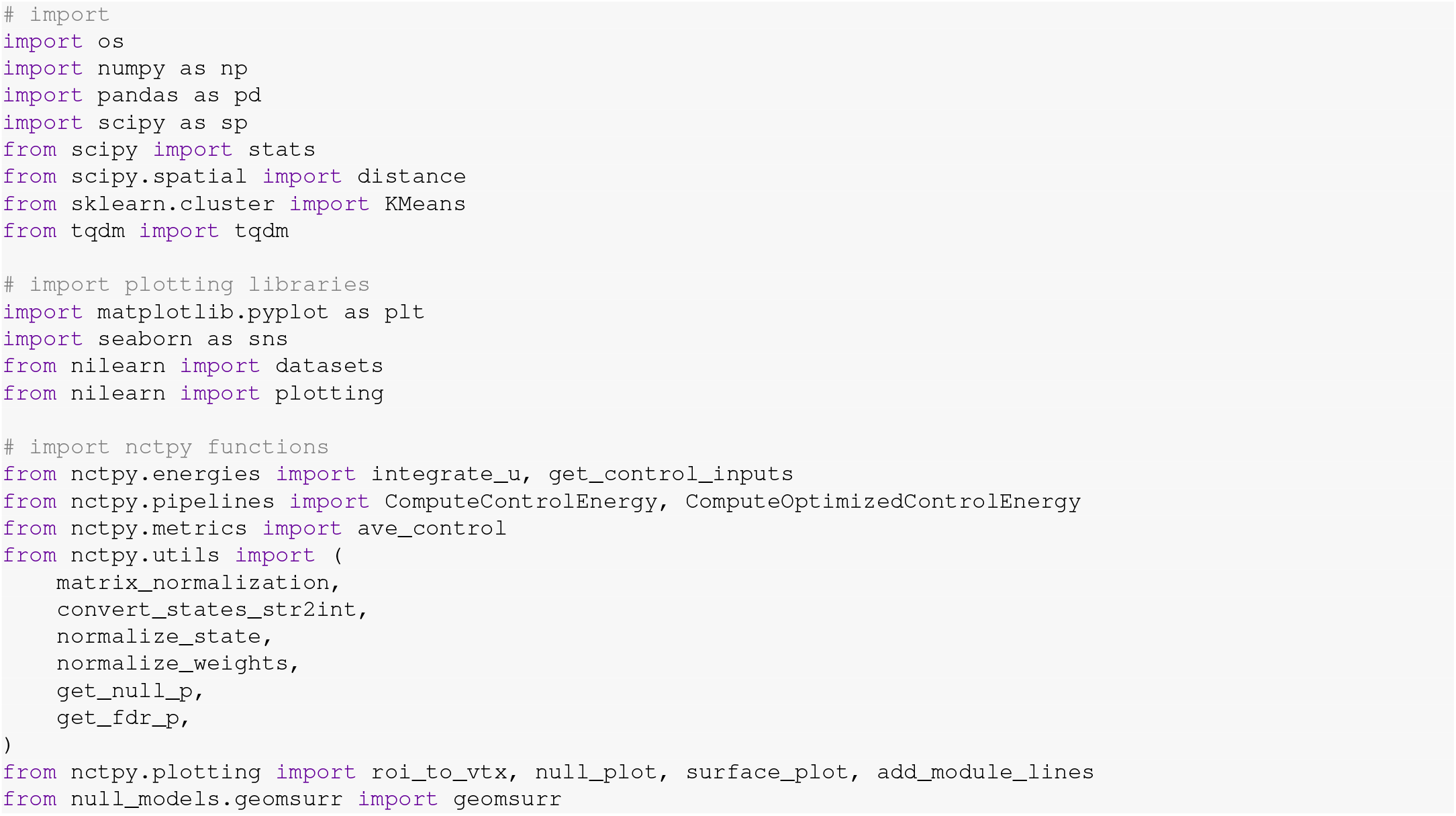

Note that depending on their goals, researchers may only need a subset of this import call. Next, we will load a structural connectome as our adjacency matrix:

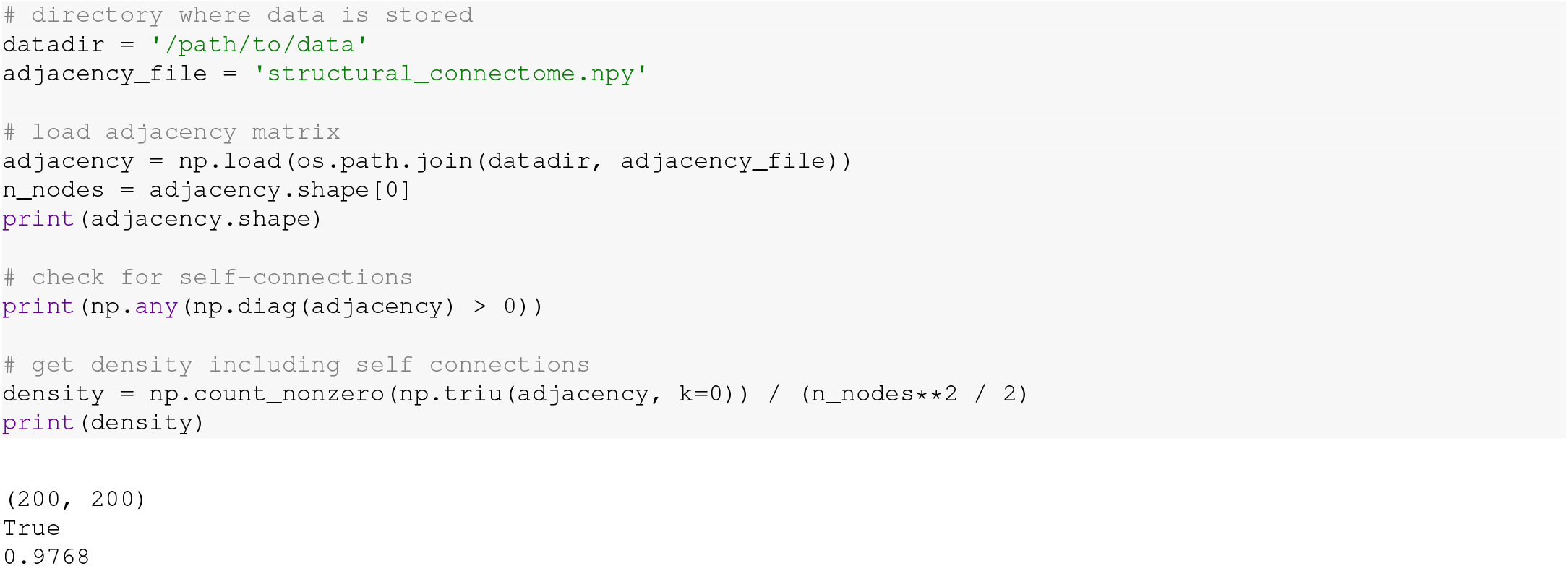

The above code demonstrates that our connectome comprises 200 nodes, includes self-connections (i.e., *A*_*ij*_ *>* 0), and has an edge density of 98%. See Figure S1 for *control energy* plotted as a function of edge density.

## IX. PROCEDURE

### Core Steps

#### Discrete-time versus continuous-time dynamical system

##### 1. CRITICAL Define a time system

The first step is to determine whether to model the linear dynamical system in discrete-or continuous-time. In a discrete-time system, the states of the system, ***x***(*t*), evolve forward in time according to a set of discrete steps (***x***(*t*) →***x***(*t* + 1)). In a continuous-time system, the states of the system are continuously changing in time 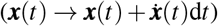. The choice of time system will depend upon the research question and affects all subsequent analyses owing to differences in the mathematical implementation of NCT under each system. We refer the reader to Karrer *et al*. [8], Kim *et al*. [7], Hespanha [47], and other texts in linear systems theory for extended discussion.

##### 2. CRITICAL Normalize adjacency matrix (Timing: *<* 1 second)

Once a time system has been determined, the first practical step in this protocol is to normalize the adjacency matrix, *A*. Normalizing *A* prior to modeling the dynamical system ensures stability and that the system will not grow to infinity over time. We include a function that normalizes *A* appropriately depending on the researcher’s chosen time system.

A) Normalizing for discrete-time systems:

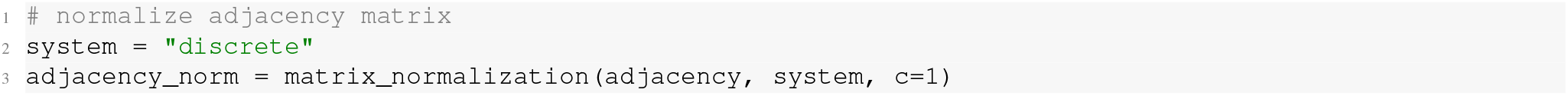

The above call will normalize *A* according to the following equation:

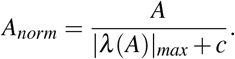

Here, |*λ* (*A*)|_*max*_ denotes the largest absolute eigenvalue of the system. Additionally, *c* is a user-defined input parameter that determines the rate of decay of system dynamics. We set *c* = 1 by default, which ensures that all modes of the system decay and thus that activity goes to zero over time (note, this is true of any positive *c* value). This normalization ensures that internal dynamics decay in a manner that is necessary for the stabilization of the system. Specifically, the largest absolute value of a matrix’s eigenvalues is called the *spectral radius*, and this normalization ensures that the spectral radius is less than 1: a condition known as *Schur stability*. Intuitively, a discrete-time system given by Eq. 1 with no input (i.e. ***u***(*t*) = 0) will evolve as ***x***(*n*) = *A*^*n*^***x***(0), and the most unstable eigenmode of the system will evolve as 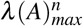. To ensure that this mode does not grow infinitely with *n*, it must have a magnitude less than 1.

B) Normalizing for continuous-time systems:

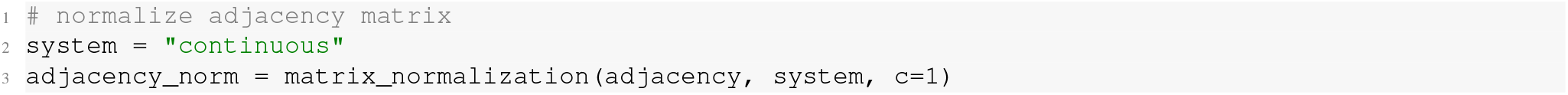

The above call will normalize *A* according to the following equation:

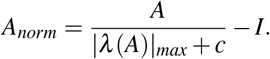

Here, *I* denotes the identity matrix of size *N*×*N*. As above, we normalize such that the spectral radius is less than one, but we take the additional step of subtracting the identity. This step exists because a continuous-time system given by Eq. 2 with no input will evolve as ***x***(*t*) = *e*^*At*^ ***x***(0), and the eigenmodes of the system as 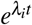. Hence, for the system to decay, all *λ*_*i*_ must have a negative real component, which is achieved through the subtraction of *I*.

Irrespective of the chosen time system, the above step outputs *A*_*norm*_, which contains the structural connectome as a normalized adjacency matrix that is ready for NCT analysis. Importantly, there are several different approaches to normalizing the matrix. Irrespective of approach, the key properties to ensure are the stability of the system as well as the preservation of as much structural information as possible. In all of the code and results shown below, *A*_*norm*_ was produced for a continuous-time system.

#### A. Protocol Pathway A: Control Energy

##### 3. Define a control task: binary brain states (Timing: *<* 1 second)

To calculate the *control energy* required to complete a state transition, researchers must first define a control task. A control task comprises an initial state, *x*0, a target state, *xf*, and a control set, *B*. In other words, a control task involves defining a specific set of control nodes that will be used by the model to transition from an initial state to a target state.

Here, we illustrate an example control task where NCT is used to transition between a pair of *binary brain states* controlled by a *uniform full control set*. As mentioned above, our *A*_*norm*_ is ordered according to 7 canonical brain systems [115]. We leverage this grouping to define a state transition between the visual system and the default mode network (DMN). To begin, we set up a vector, `states`, that stores integer values denoting which brain system each node belongs to. That is, `states == 0` represents nodes that belong to system 1, `states == 1` represents nodes that belong to system 2, *etcetera*. We create `states` from a list of strings that groups nodes into the aforementioned canonical brain systems (this file can be found here):

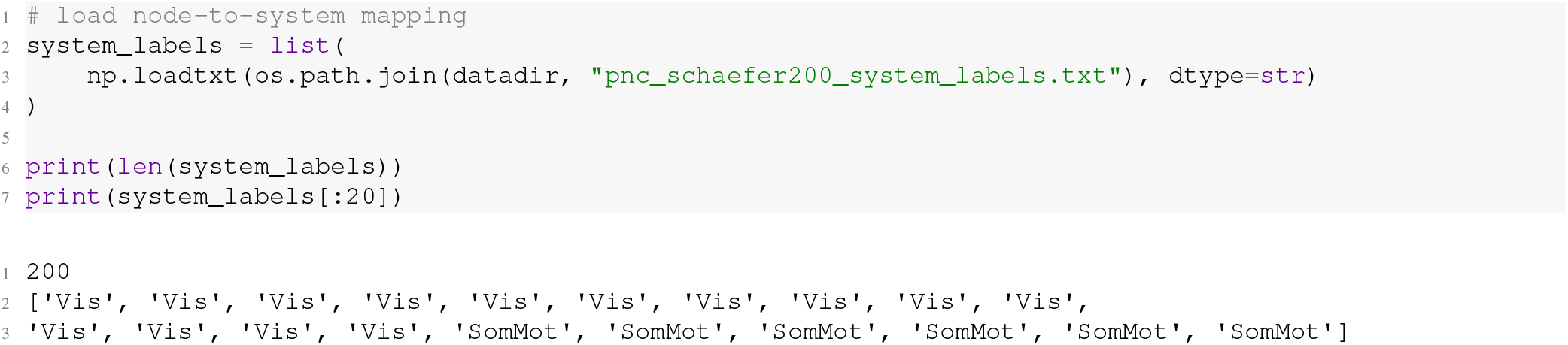

In `nctpy`, we include a function called `convert_states_str2int` that will convert this list of strings for us:

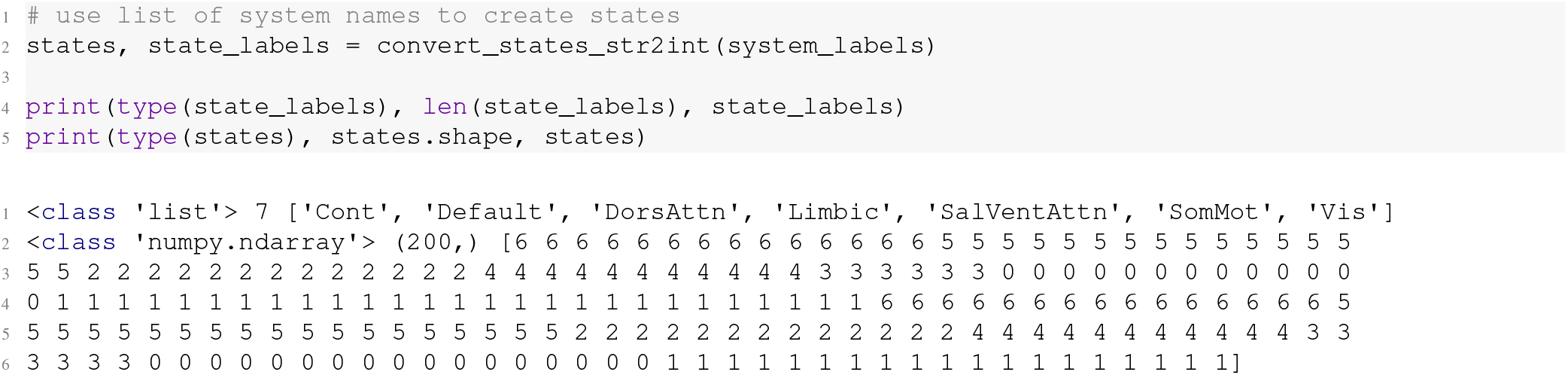

As can be seen from the above `print` commands, our `system_labels` variable comprises a string variable for every node in our system that denotes which brain system that node belongs to. `convert_states_str2int` takes that list of strings and returns an array of integers, `states`, with a corresponding list of labels, `state_labels`. Below, we extract *x*0 and *xf* using the integers that correspond to the visual system (`‘Vis’`) and the default mode system (`‘Default’`):

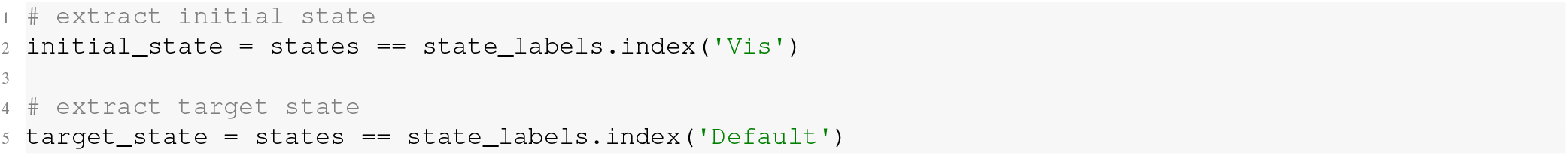

`initial_state` and `target_state` will be Boolean vectors ([**True, False**]), wherein `True` encodes the nodes that belong to a given state. Next, we normalize the state magnitude using our included function, `normalize_state`:

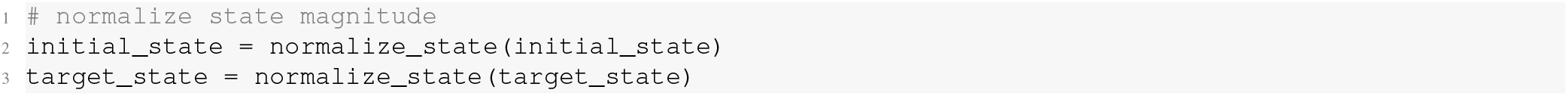

This process will convert `initial_state` and `target_state` from Boolean entries to floating point numbers that have been normalized using the Euclidean norm of the vector. This normalization constrains state magnitude to a unit sphere (see Section V for more details). Finally, we define our control set. Unlike the initial and target states, the control set is encoded along the diagonal of an *N*×*N* matrix, *B*. Here, we use a *uniform full control set*; thus, we can define our control set as the identity matrix:

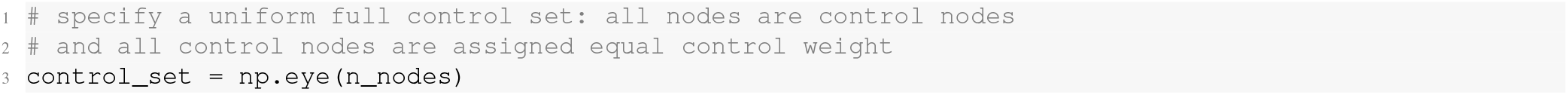

##### 4. Compute control signals and state trajectory (Timing: *<* 1 second)

Following the definition of a control task, the next step is to find the control signals, ***u***(*t*), that drive the system to transition between *x*0 and *xf*. ***u***(*t*) will be an *m*×*T* matrix of *m* time-varying signals injected into the control nodes over a specified time horizon, *T*. Here, owing to our use of a full control set, *m* = *N*. Critically, injecting these control signals into a system whose initial state is encoded by *x*0 should result in a system whose final state is encoded by *xf* at time *T*. Alongside the control signals, we also extract the state trajectory, ***x***(*t*). The state trajectory, which will be an *N*×*T* matrix, is the time-varying pattern of simulated neural activity that unfolds as the system traverses between *x*0 and *xf*. In this protocol, we find ***u***(*t*) and ***x***(*t*) using the `get_control_inputs` function:

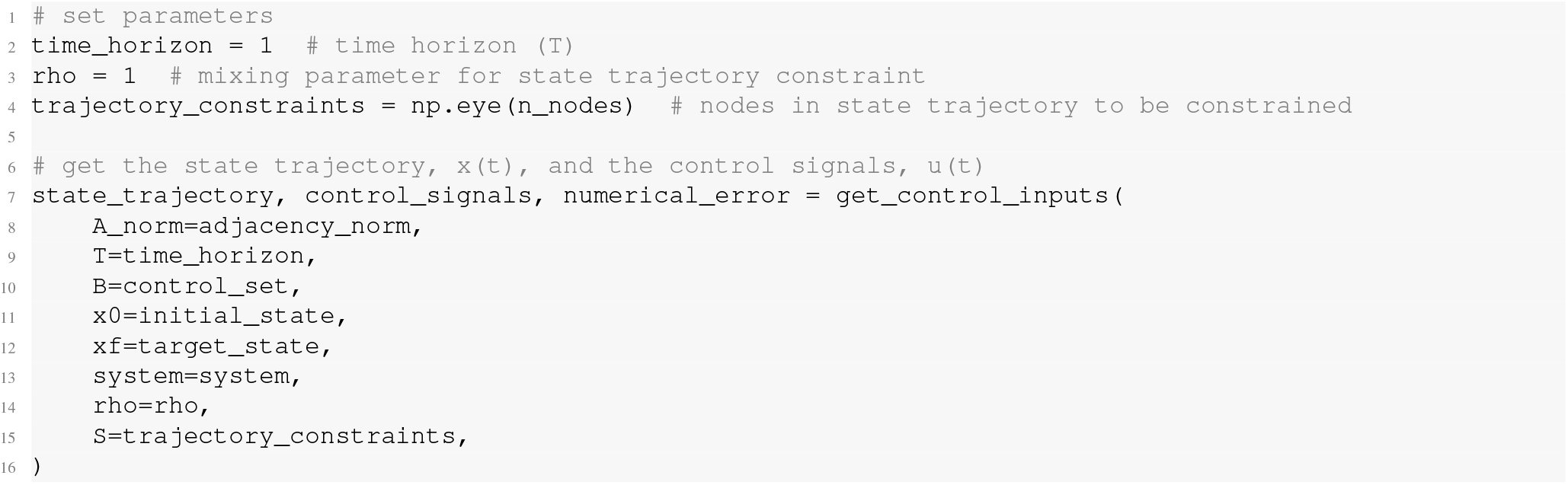

By default, we set `time_horizon=1`. Note that this value is arbitrary and does not correspond to any real-world time units (e.g., seconds). Importantly, `get_control_inputs` utilizes a cost function that includes both the magnitude of the control signals and the magnitude of the state trajectory. The input parameter `rho` allows researchers to tune the mixture of these two costs while finding the input ***u***(*t*) that achieves the state transition. Specifically, `rho=1` places equal cost over the magnitude of the control signals and the state trajectory. Reducing `rho` below 1 increases the extent to which the state trajectory adds to the cost function alongside the control signals. Conversely, increasing `rho` beyond 1 reduces the state trajectory contribution, thus increasing the relative prioritization of the control signals. Lastly, `S` takes in an *N*×*N* matrix whose diagonal elements define which nodes’ activity will be constrained in the state trajectory. In summary, `S` designates which nodes’ neural activity will be constrained while `rho` determines by how much it will be constrained, relative to the control signals. Here, by setting `rho=1` and `S=np.eye(n_nodes)`, we are implementing what we refer to as *optimal control* [11]. Alternatively, researchers may choose to constrain only a subset of the state trajectory by defining partial constraint sets. If `S` is set to an *N*×*N* matrix of zeros, then the state trajectory is completely unconstrained; we refer to this setup as *minimum control* [20, 51]. In this case, `rho` is ignored. See here for a notebook outlining different use cases of `get_control_inputs`.

In addition to `state_trajectory` and `control_signals`, `get_control_inputs` also outputs `numerical_error`, which stores two forms of numerical error. The first error is the *inversion* error, which measures the conditioning of the optimization problem. If this error is small, then solving for the control signals was well-conditioned (see Section VII). The second error is the *reconstruction* error, which is a measure of the distance between *xf* and ***x***(*T*). If this error is small, then the state transition was successfully completed; that is, the neural activity at the end of the simulation was equivalent to the neural activity encoded by *xf*. We consider errors *<* 1^*−*8^ as adequately small:

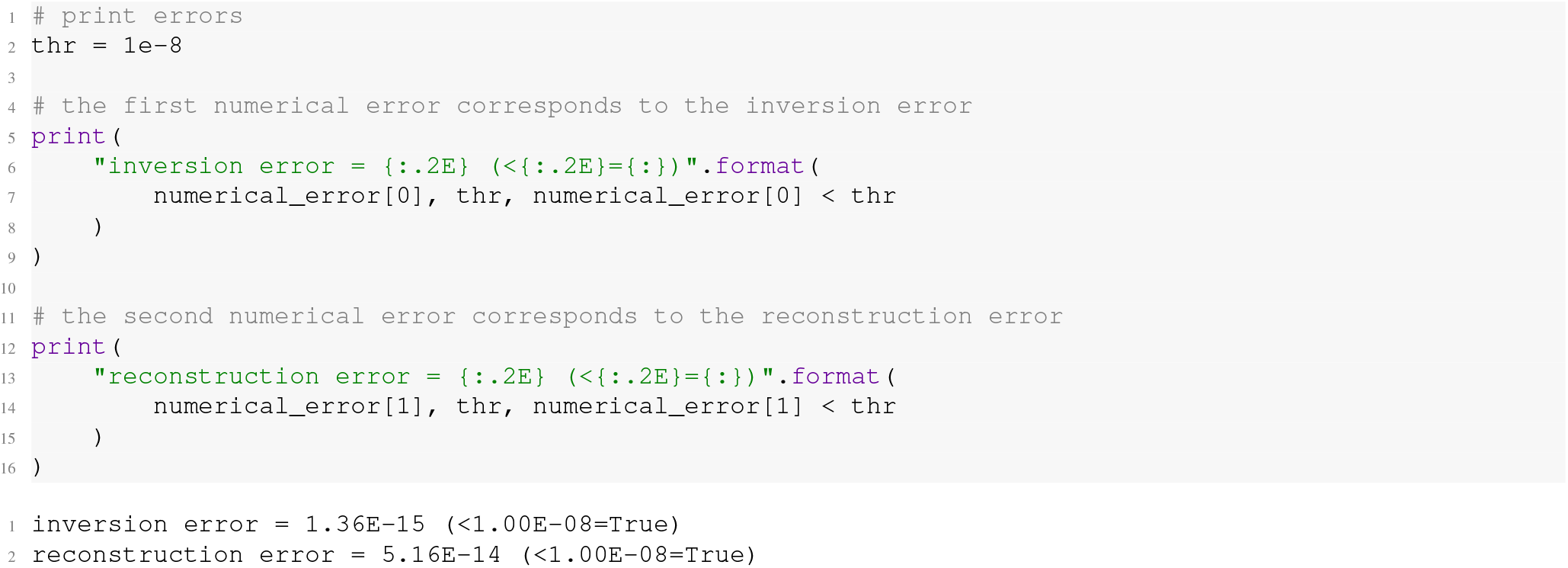

##### 5. Visualize state trajectory and control signals (Timing: *<* 1 second)

Once ***x***(*t*) and ***u***(*t*) have been derived, they should be visualized before computing *control energy*. Visualization provides intuition regarding how the model is behaving and is helpful for confirming that the state transition was completed successfully. As noted in Section V, completion of a state transition is not guaranteed by the model, and an incomplete transition may necessitate revising either the control set (e.g., to provide more control over the system if a partial control set was used) or the time horizon (e.g., to provide more time for the model to complete the transition). We suggest the following simple plot:

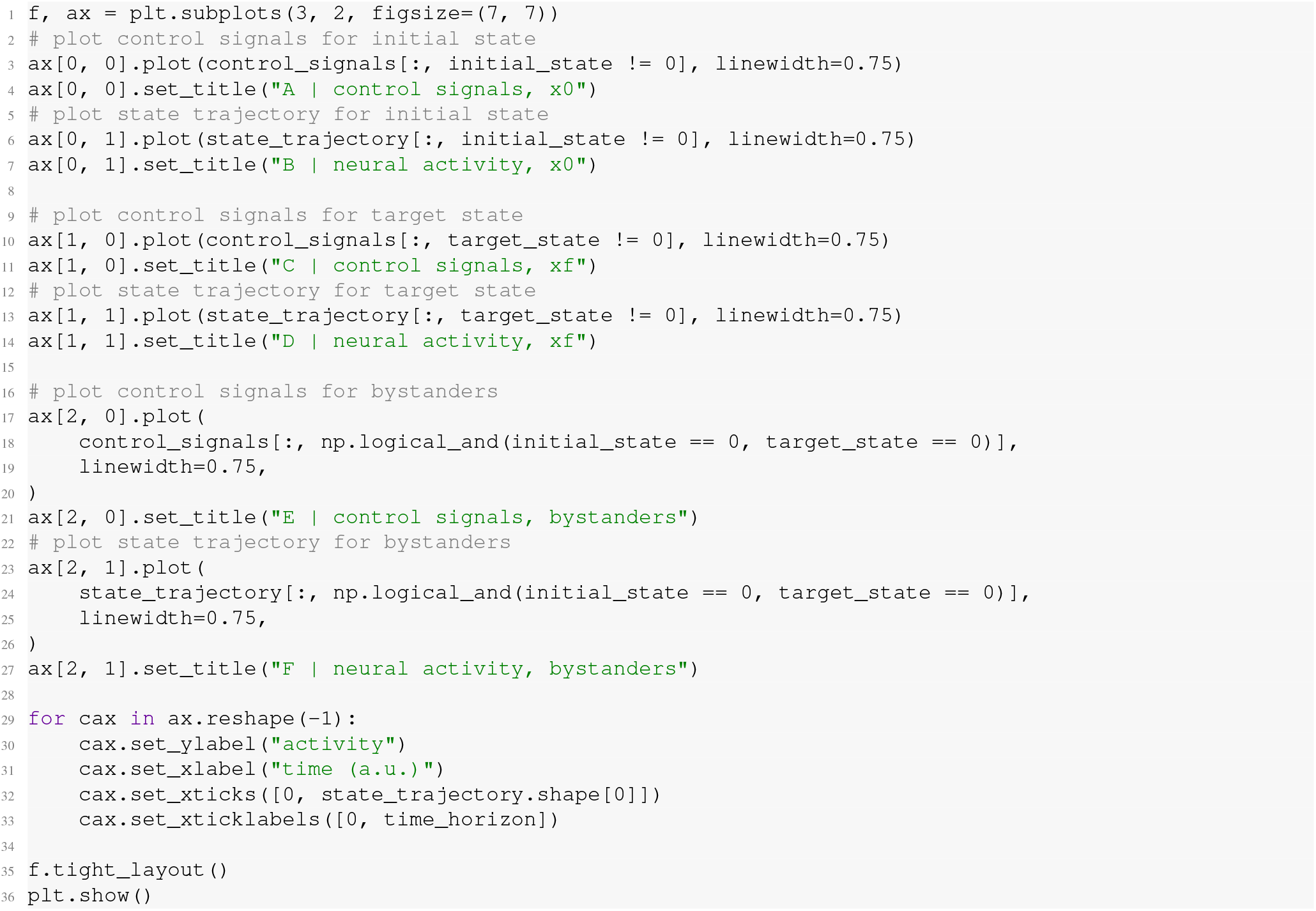

This plot (Figure 4) shows the control signals (left column) alongside the state trajectory (i.e., neural activity; right column) separately for nodes within the initial (top row) and the target state (middle row), as well as the bystanders (bottom row). Note, we define bystanders as nodes that are outside both the initial and target states. We choose this division of nodes as it provides several simple intuitions about model behavior. First, we can see that the model is driving negative time-varying control signals into the nodes of the initial state (Figure 4A), which drives their activity to 0 over time (Figure 4B). Second, we can see that the model is driving positive time-varying control signals into the nodes of the target state (Figure 4C), which drives their activity from 0 to ∼0.15 over time (Figure 4D). Note that ∼0.15 represents the maximum neural activity following state normalization for the states presented here; this maximum activity may vary depending on state definition. Finally, we can see that diverse time-varying control signals are being injected into the bystander nodes (Figure 4E), which are in turn guiding changes in these regions’ activity (Figure 4F). In other words, Figure 4 shows that the model is performing a combination of “turning off” the initial state, “turning on” the target state, as well as guiding diffusing activity toward the target state via the bystanders. Figure 4 also provides a simple visual way to check whether the state transition completed successfully; at the end of the simulation, it is apparent that activity in the target state is maximal while activity in the initial state and bystanders is 0, which accords with our definition of *xf*. This behavior explains the low reconstruction error shown above. Additionally, this plot allows researchers to visualize how model behavior varies under different control sets (see Section 6 below as well as Figures S2, S3, S4, and S5) and time horizons (see Figures S6, S7, and S8). Note that while we view Figure 4 as the simplest way to plot initial model outputs, it is only one of many options. Researchers may choose to plot ***x***(*t*) and ***u***(*t*) as heatmaps or on the brain’s surface, which would facilitate visualization of spatial patterning (see step 3a in Section 6 for an example).

**FIG. 4.**
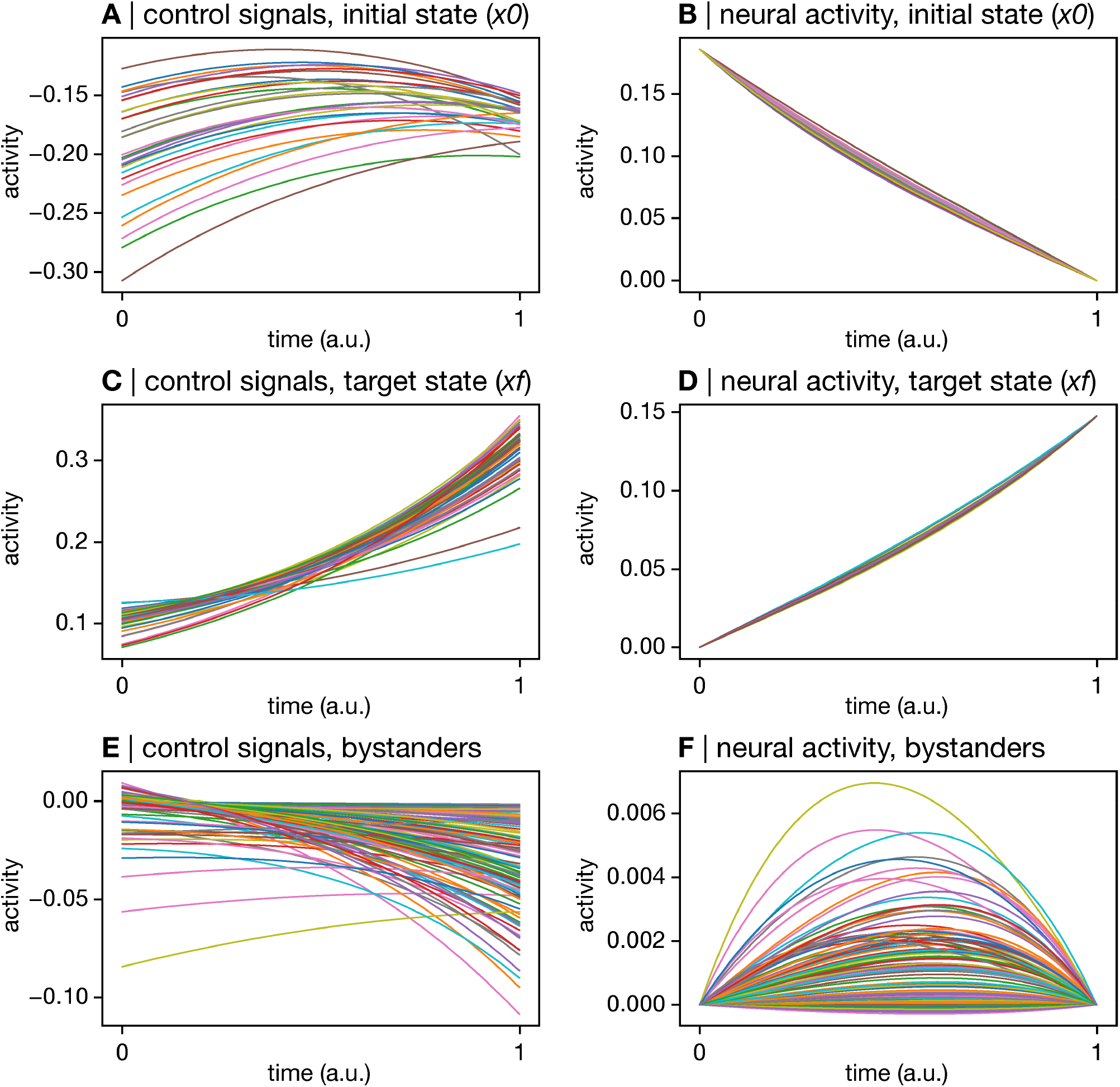
Visualize the control signals and the state trajectory. For a given state transition, the control signals (***u***(*t*), left column) and the state trajectory (***x***(*t*), right column) should be visualized. Here, owing to our use of binary brain states, we group this visualization by nodes in the initial state (*x*0, top row), the target state (*xf*, middle row), and the remaining nodes (bystanders, bottom row). This plot provides intuition on model behavior by showing the kinds of control signals that are driving specific changes in neural activity. The top row shows that the model is driving negative time-varying control signals into the nodes of the initial state (**A**), which drives their activity to 0 over time (**B**). The middle row shows that the model is driving positive time-varying control signals into the nodes of the target state (**C**), which drives their activity from 0 to ∼0.15 over time (**D**). The bottom row shows that the model is driving diverse time-varying control signals into the bystander nodes (**E**), which are in turn guiding changes in these regions’ activity (**F**).

##### 6. Compute control energy (Timing: *<* 1 second)

The final step is to summarize the control signals into *control energy*. This is done by numerically integrating the control signals over time. In this protocol, we use Simpson’s rule—an extension of the trapezoidal rule that fits a polynomial through neighboring sets of points—to achieve this integration, yielding a vector of node-level energy:

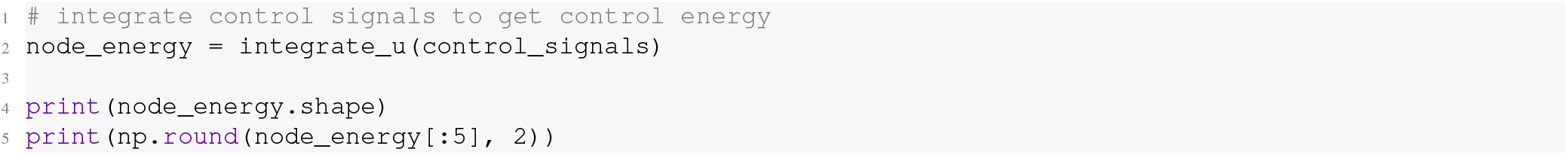
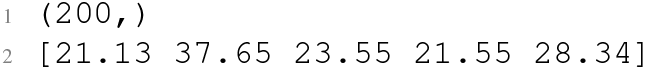

Finally, these node-level energies can be summed to produce a single estimate of *control energy*

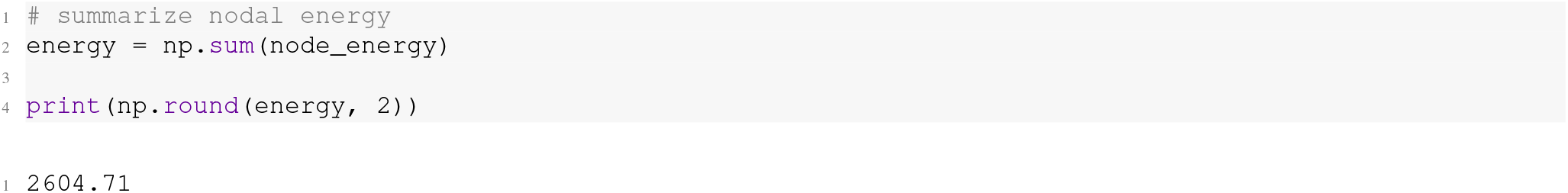

#### Wrapping Pathway A for ease of use

Steps 1-6 outline how to extract the *control energy* for a single control task, which we defined as completing a state transition between the visual system and the default mode system using control signals delivered to all system nodes (to view the above steps in a single notebook, see here). Alternatively, researchers may want to examine many control tasks within the context of a single study. Thus, in `nctpy` we include a *Python* class called `ComputeControlEnergy` that wraps all of the above steps (excluding step 5) and applies them over a list of control tasks. In addition to an adjacency matrix, `ComputeControlEnergy` expects a dictionary as input wherein each entry is a control task that includes: (i) an initial state, *x*0; (ii) a target stage, *xf* ; (iii) a control set, *B*; (iv) a matrix of state trajectory constraints, *S*; and (v) a constraint parameter, *rho*. For example, using the `states` variable defined above, this dictionary could be created as follows:

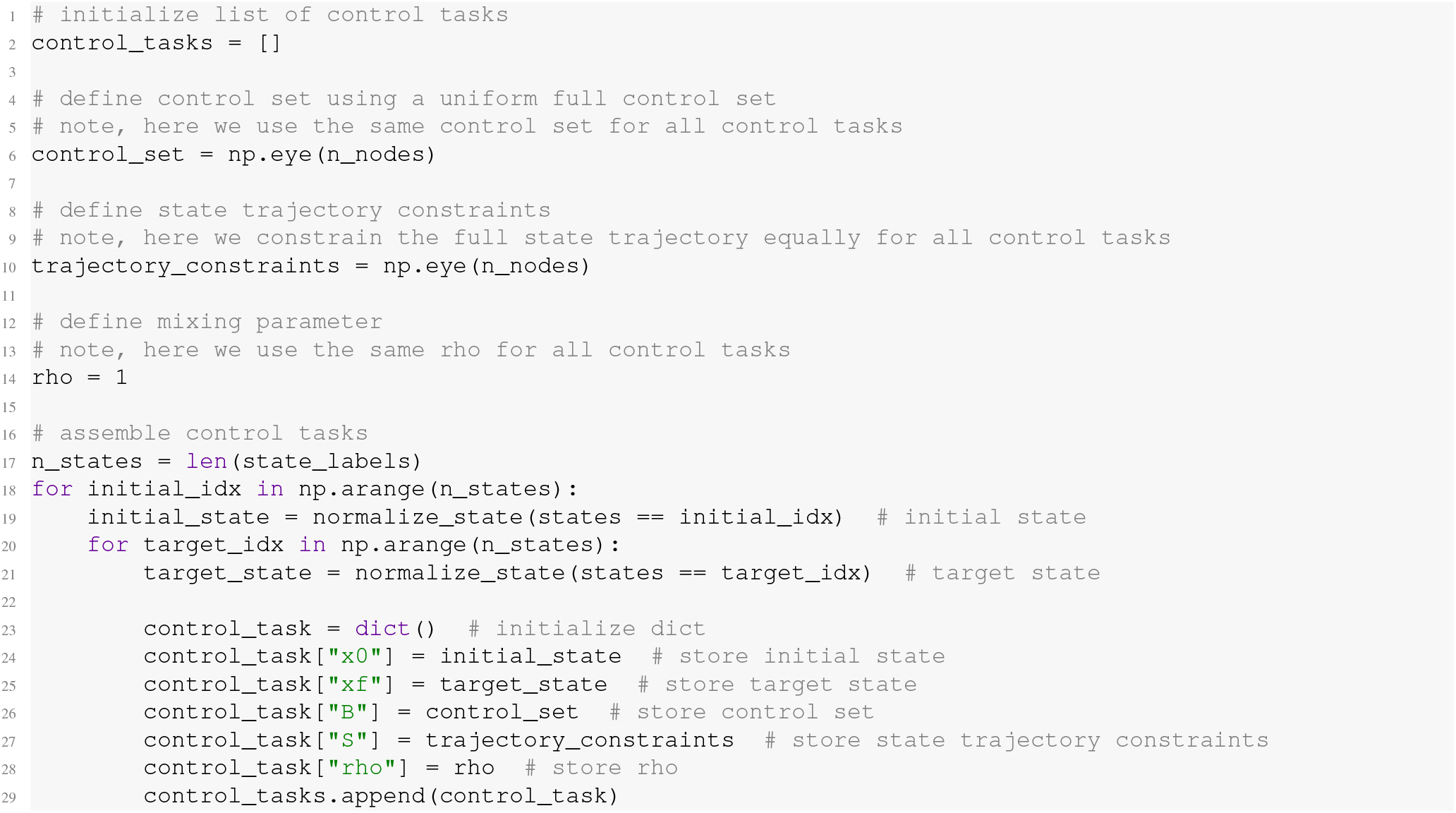

Note that for simplicity, the above section of code assumes the same control set (*B*), trajectory constraints (*S*), and *ρ* for all control tasks, but researchers may vary these parameters over transitions according to their needs. Next, we initialize and run `ComputeControlEnergy`:

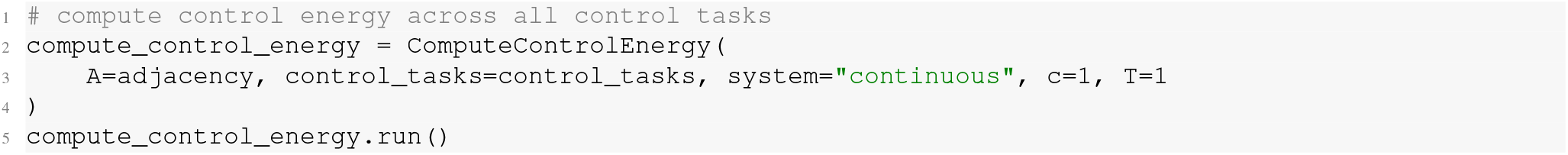

`ComputeControlEnergy` will perform matrix normalization internally, and hence here we input *A* rather than *A*_*norm*_. Apart from the `control_tasks` variable, `ComputeControlEnergy` also requires that the user specify the time system (`system=‘continuous’`), normalization constant (`c=1`), and time horizon (`T=1`) as input arguments. Once instantiated, `ComputeControlEnergy.run()` will run steps 1-6 (excluding step 5). Once completed, a single estimate of *control energy* per control task will be stored in `ComputeControlEnergy.E`. Note that `ComputeControlEnergy` will not output the state trajectory, control signals, or node-level energy. These energy values can be trivially reshaped into a matrix and visualized (Figure 5):

**FIG. 5.**
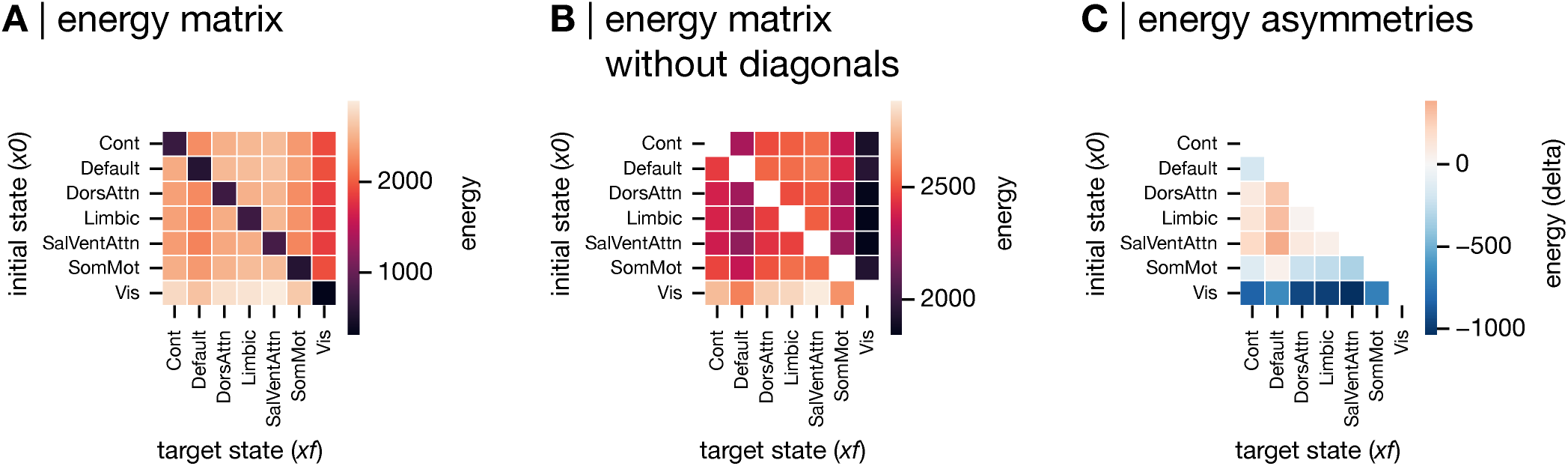
Visualize control energy. In cases where multiple state transitions are considered, we recommend visualizing *control energy* using the following heatmaps. **A**, Full energy matrix. The full energy matrix shows energy for all state transitions, including those where the initial and target state are the same (diagonal elements). This form of energy is referred to as persistence energy and is interpreted as the amount of effort required to maintain neural activity in a given state. Persistence energy is typically lower than the energy associated with transitioning between different states. **B**, Between-state energy matrix. In order to visualize variance in *control energy* for transitions between states, we recommend plotting the energy matrix without the diagonal as well. **C**, Energy asymmetry matrix. Finally, we recommend subtracting the transpose of the energy matrix and visualizing the lower (or upper) triangle of the ensuing asymmetry matrix. Doing so allows researchers to see the asymmetries present in the *control energy*.

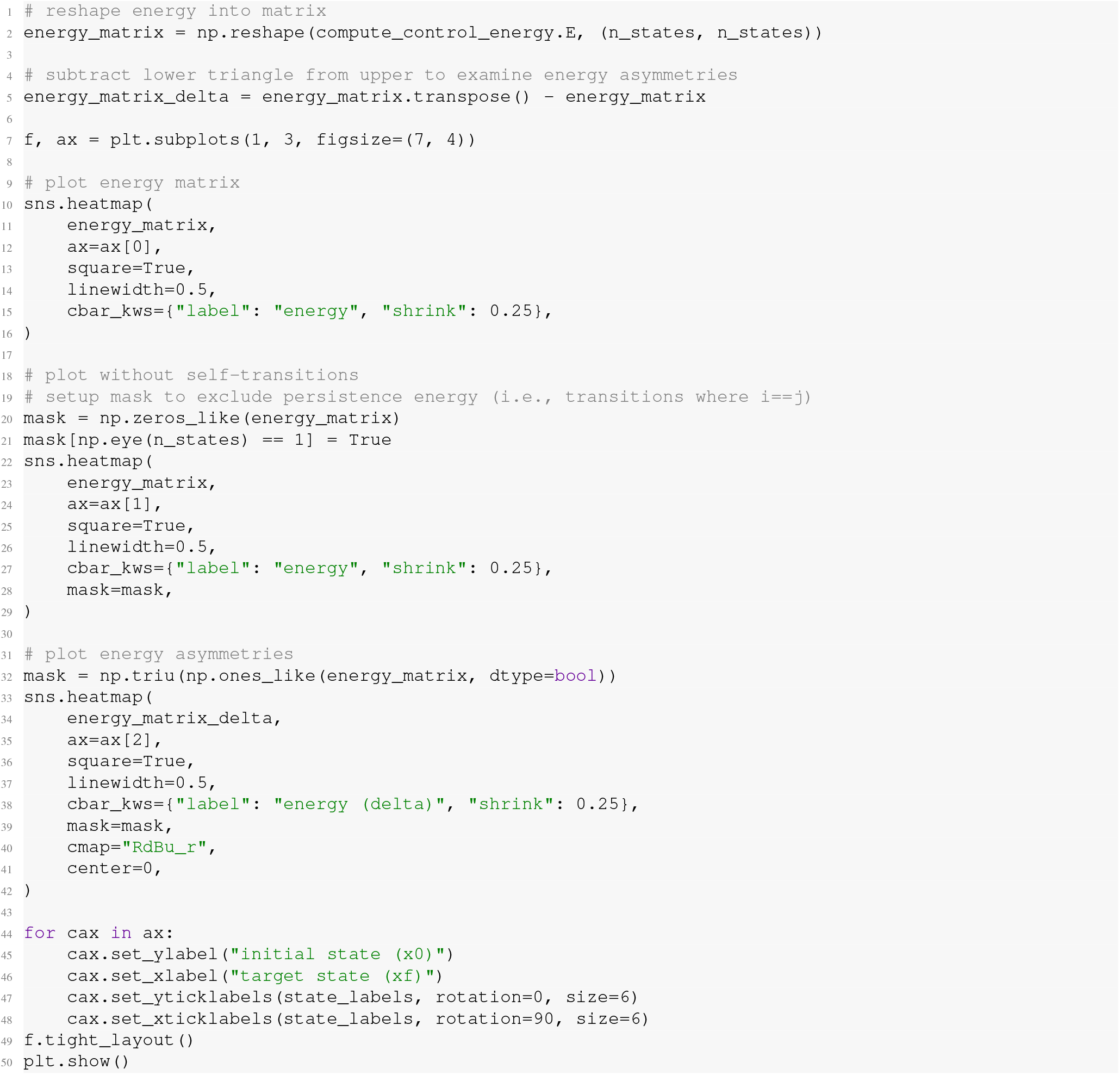

In Figure 5A, the diagonals represent the energy associated with control tasks that have identical initial and target states. We refer to these energies as persistence energy [20, 73, 145], which we interpret as the amount of *control energy* required to maintain a given brain state (see Figure S9 for a persistence energy example of Figure 4). Persistence energy is typically lower than the energy for control tasks wherein the initial and target states differ (i.e., the off-diagonal elements of `energy_matrix`). Thus, in order to better visualize the variance in energy across state transitions, we recommend also plotting `energy_matrix` excluding the diagonal elements (Figure 5B). Additionally, as the energy associated with transitioning from *x*0 to *xf* is not necessarily equivalent to that associated with the reverse direction, researchers may subtract the upper and lower triangles of `energy_matrix` to examine energy asymmetries (Figure 5C). Indeed, we have done this in our previous work (see [51, 73]).

### Variations to Pathway A

All of the above constitutes our complete protocol for calculating *control energy*. However, there are multiple variations to the above protocol that researchers may wish to consider depending on their research goals. In this section, we illustrate a selection of these variants that are likely to be of broad interest to the field of neuroscience (Figure 3C). These variations include (A) studying non-binary brain states; (B) implementing partial and non-uniform control sets; and (C) examining directed structural connectomes.

#### Non-binary brain states

In Section IX A, we illustrated a state transition between the visual system and the default mode system using a binary definition of brain states extracted from a canonical system-level grouping of brain regions [66]. Below, we provide an example of using non-binary brain states instead. This example draws on aforementioned work from Cornblath *et al*. [20], who modeled the energy required to transition between clusters of rs-fMRI activity.

3A) *Define a control task: non-binary brain states*. To define non-binary brain activity states, we cluster rs-fMRI data along the time dimension to extract co-activation states [20]. To achieve this goal, we first load the rs-fMRI data for 253 participants extracted from the same parcellation that defined our structural connectome. Then, we concatenate these time series end-to-end across subjects:

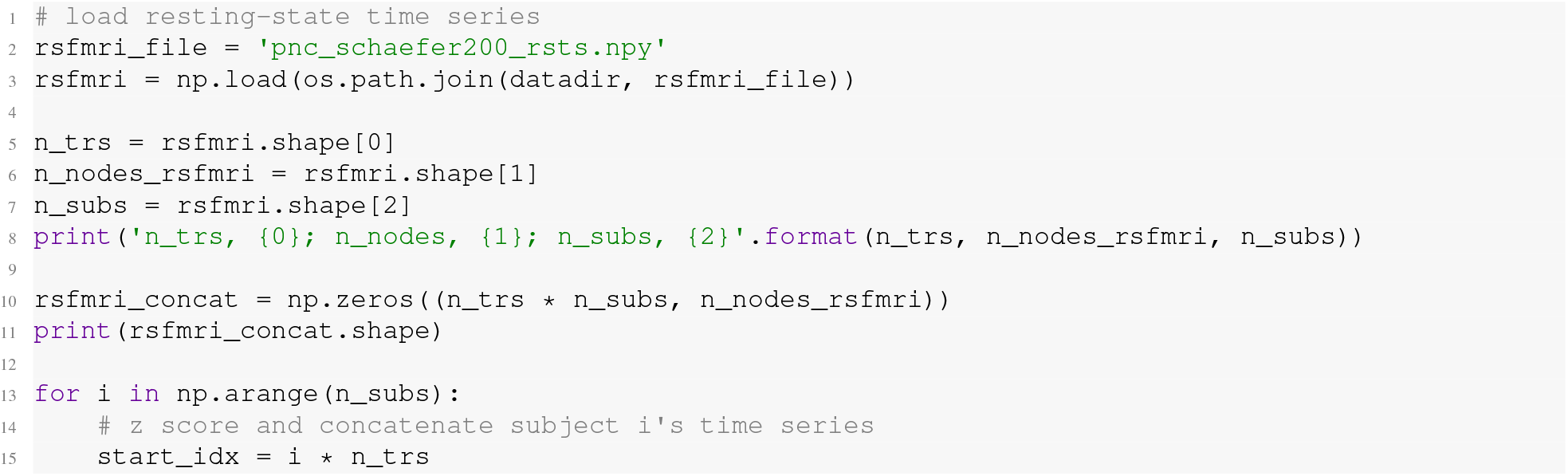
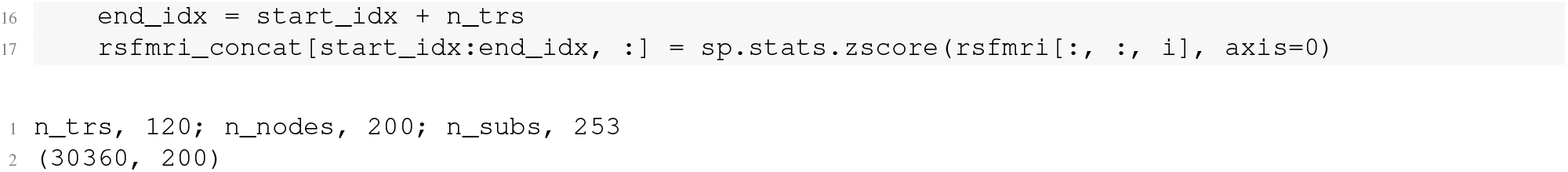

This process assigns empirical time series to each of the system nodes comprising 30,360 time points (120 TRs by 253 participants) of rs-fMRI data. Next, we cluster these data in time using K-means and visualize the corresponding co-activation states on the cortical surface. To generate this plot, we include a function called `surface_plot` that utilizes tools from Nilearn (https://nilearn.github.io/stable/index.html):

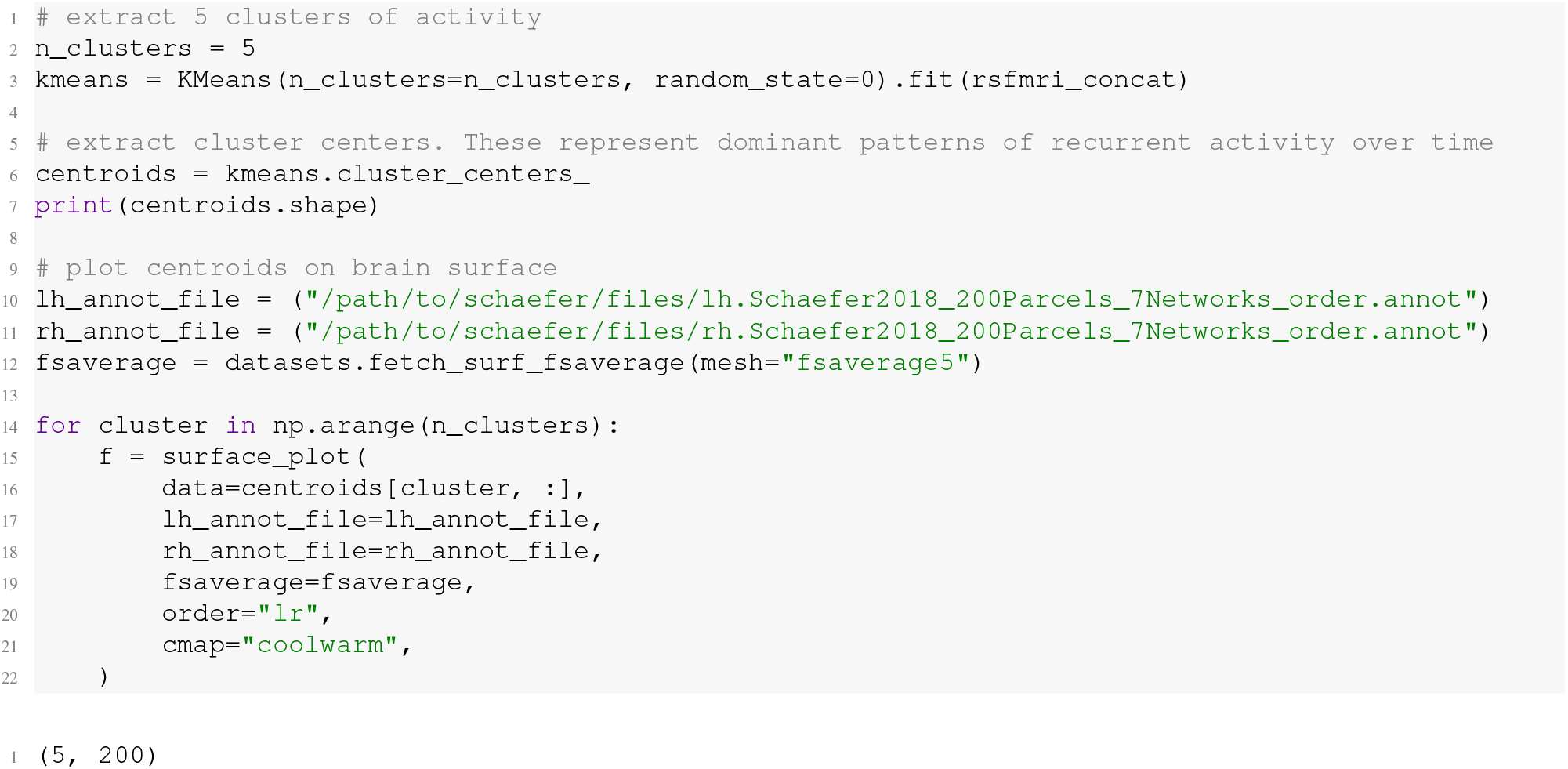

Figure 6 illustrates the centroids for 2 of the 5 clusters we extracted using K-means. As we are focused on illustrating a single state transition for the purposes of this protocol, the remaining 3 centroids are not shown (see Cornblath *et al* [20] for detailed discussion of all 5 clusters). These centroids represent patterns of activity that recur throughout our concatenated time series. The spatial patterning of these 2 centroids indicate activity concentrated in visual cortex (Figure 6A) and the default mode system (Figure 6B), respectively. Using the same functions outlined above, we can extract this pair of centroids as brain states and recompute the control energy:

**FIG. 6.**
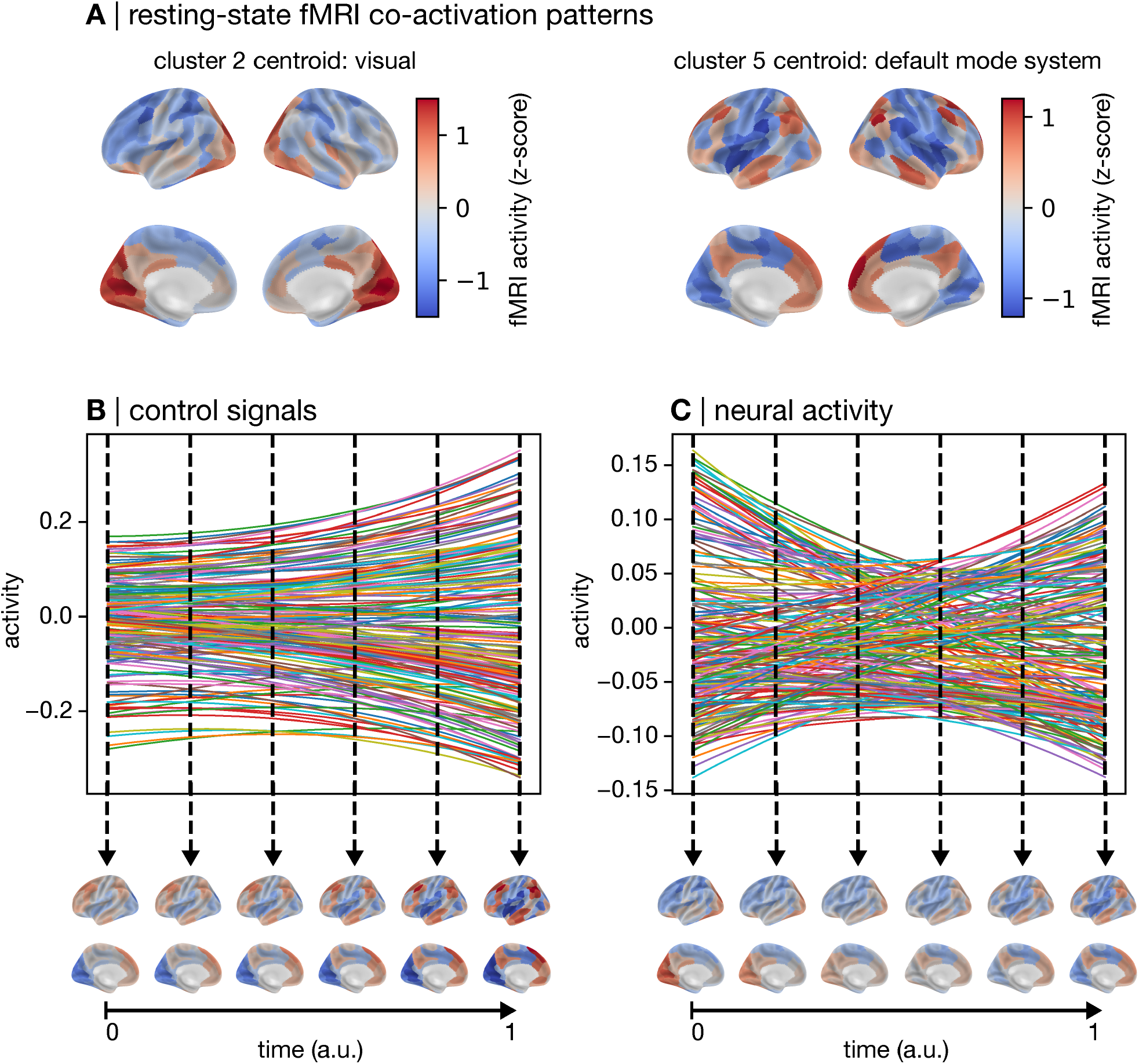
Control signals and state trajectory for non-binary brain states derived from resting-state fMRI. We applied K-means clustering to resting-state fMRI time-series to extract patterns of co-activation. These patterns were used as non-binary brain states for network control theory (NCT) analysis. **A**, Resting-state fMRI clusters that represent visual system activity (left) and default mode activity (right). The control signals and the state trajectory were modeled by assigning the visual system to the initial state and the default mode system to the target state. **B**, Control signals visualized using line plots (top) and on the cortical surface for select time points (bottom). **C**, State trajectory (neural activity) visualized using line plots (top) and on the cortical surface for select time points (bottom).

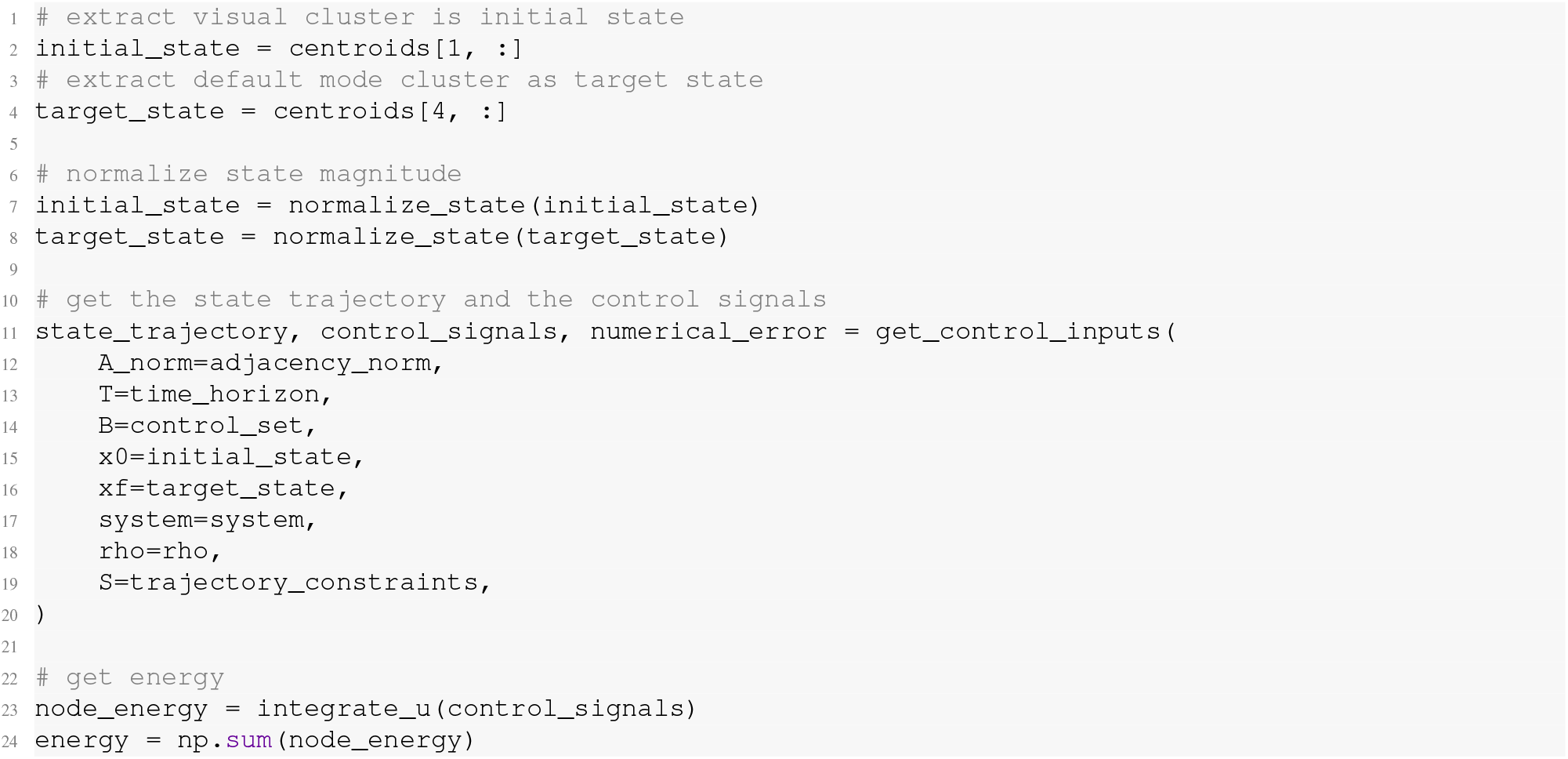

As stated above in step 5, the temporal unfolding of ***x***(*t*) and ***u***(*t*) can be visualized using line plots (Figure 6B top, C top). Alternatively, to illustrate the spatial patterning of the state transition, ***x***(*t*) and u(t) can be visualized on the cortical surface for specific time points (Figure 6B bottom, C bottom):

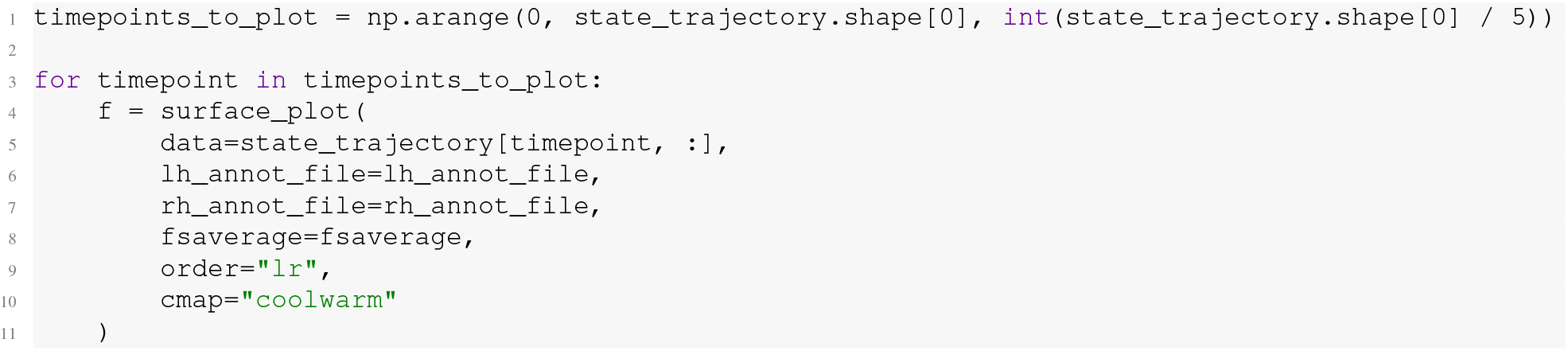

#### Partial and variable control sets

In Section IX A, we illustrated a state transition controlled via a *uniform full control set*. Such a control set amounts to assigning all nodes of the system the same degree of control over system dynamics. Below, we provide examples of using alternatives to this regime that involve using variable (instead of uniform) and partial (instead of full) control sets.

3B) *Define a control task: uniform* ***partial*** *control set*. Defining a *uniform partial control set* is straight forward. Instead of assigning the *N*×*N* identity matrix to *B*, we select specific diagonal elements to assign the value of 1, and assign 0 to the remaining diagonal (and non-diagonal) elements. Here, we illustrate the example of selecting the bystander regions (see Figure 4E, F) as our control set:

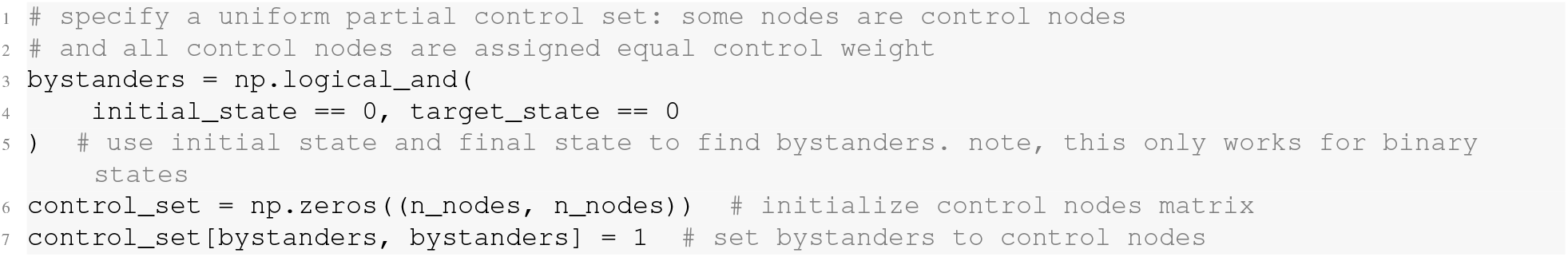

Figure S2 shows the results of generating ***x***(*t*) and ***u***(*t*) under the above control set. Note that the above code only works for binary brain states, because in this case bystanders can be trivially defined as nodes with no activity in either the initial or target states. This fact does not imply that partial control sets cannot be used for non-binary brain states. In Figure S2, we observe that the control signals are 0 for both *x*0 and *xf*, indicating that they received no control signals. Additionally, the control signals for the bystanders, as well as the neural activity of all nodes, has changed substantially compared to Figure 4. Notably, the control signals are several orders of magnitude greater for this *uniform partial control set* compared to the *uniform full control set* used above. In turn, although this state transition completes successfully, the energy we observe here is also several orders of magnitude greater (energy = 6.64 ×10^9^). See Figures S3, S4, and S5 for more examples of *uniform partial control sets*, including some for which the state transition does not complete. While the code implementation of a *uniform partial control set* is straight forward, researchers must ensure that their control set is large enough to achieve numeric stability (see Section VII).

3C) *Define a control task:* ***a priori variable*** *full control set*. Instead of assigning all nodes of the system the same degree of control over dynamics, researchers may want to make statements about which nodes should have more or less control according to their hypotheses. One way that this can be achieved is by using node-level annotation maps [146, 147] to assign control weights. For a single annotation map—which we assume is stored in `‘neuromap.npy’`—assigning weights can be achieved in the following way:

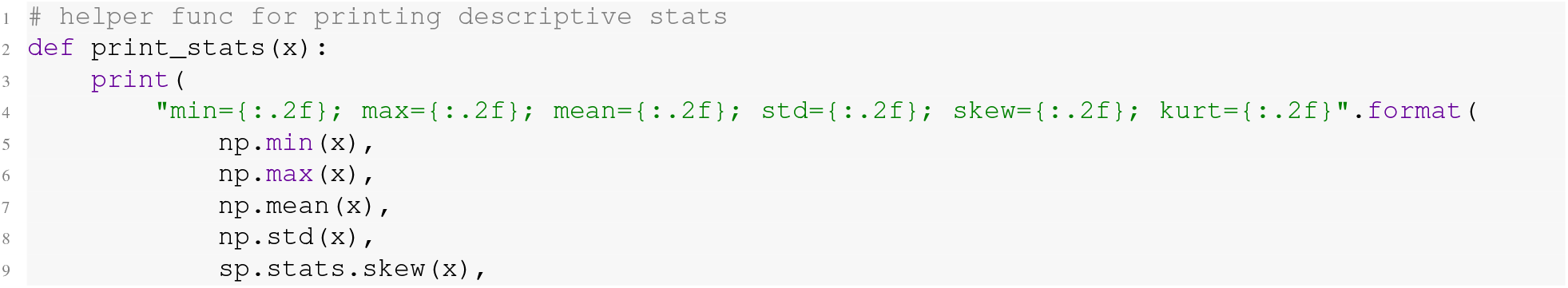
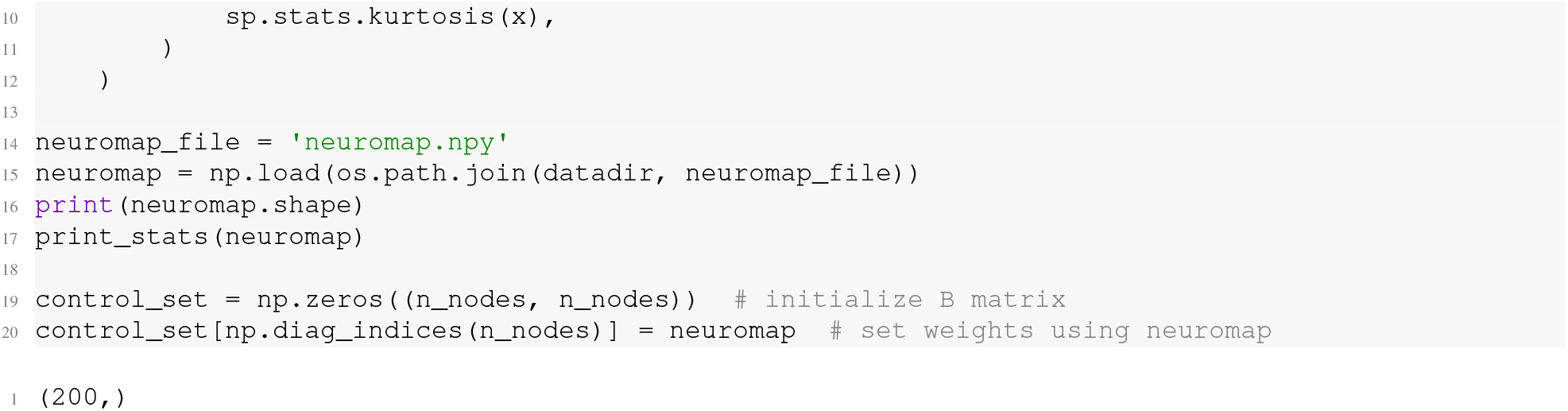
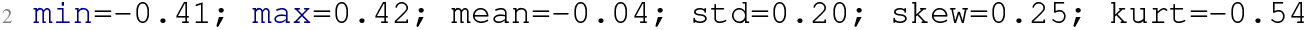

The above code demonstrates that our annotations, and therefore our control set, include both positive and negative values. The sign of a given weight will determine how a positive control signal delivered to that node influences its neural state; a positive weight will cause a positive change in neural activity, while a negative weight will cause a negative change. As control signals can also carry positive and negative values over time (see Figure 4), this behavior has no impact on *control energy*; a positive control signal driven into a node using a positive control weight will cause the same change in neural state as a negative control signal driven via a negative control weight. However, it does impact the interpretation of ***u***(*t*). Thus, to simplify the interpretation of the control weights, and of ***u***(*t*), we suggest adding a constant (1) as well as the absolute minimum value to the annotation map:

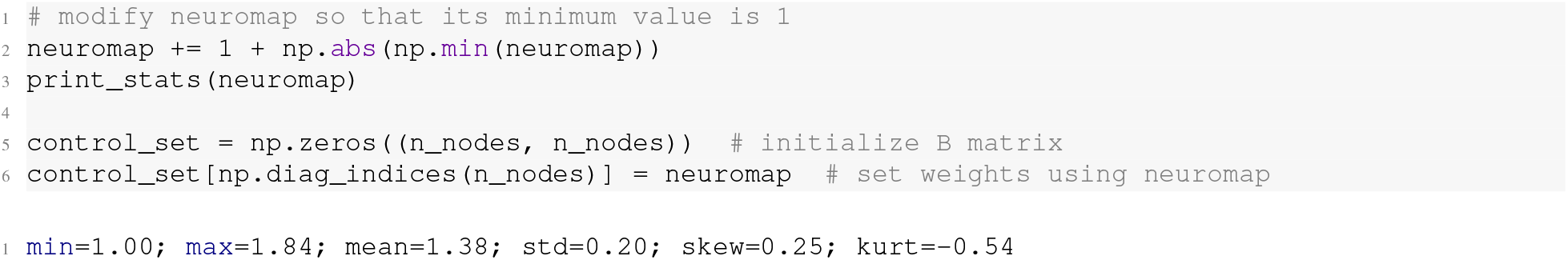

This process will create a set of weights with a minimum value of 1 and, in this case, a maximum value of 1.84 (see Figure S10 for a plot ***x***(*t*) and ***u***(*t*) derived from this control set). This fact simplifies our interpretation. For example, because all weights are positive, we can say that the highest weight node has 1.84 times more control over system dynamics compared to the lowest weight node. Additionally, whether a control signal drives a positive or negative change in neural state now depends solely on its own sign at a given point in time.

A recent study by Singleton *et al*. [30] utilized an *a priori variable full control set* by assigning control weights according to a range of seretonin receptor maps. Singleton *et al*. [30] found that the *control energy* associated with their state transition was lowest when using a 5-HT2a receptor map compared to 5-HT1a, 5-HT1b, 5-HT4, and 5-HTT maps. This result suggests that, compared to other receptors, the spatial patterning of 5-HT2a receptors yielded the most efficient state transition (i.e., by reducing *control energy* the most). The authors subsequently replicated their results using N,N-Dimethyltryptamine [31]. However, to draw this conclusion, researchers need to mindful of the following caveat. In our model, increasing the total amount of control necessarily reduces energy. This relation exists because the task of completing a given state transition is easier for the model when any node is granted a greater degree of control over dynamics than it had previously, leading to smaller amplitude control signals and thus lower energy. For example, if we compared energy between our *uniform full control set* and our above *variable full control set*, energy would be trivially lower for the latter. This difference occurs because all weights on the diagonal of *B* are 1 in our *uniform full control set*, while all but one of the weights in our *variable full control set* are *>* 1. This issue is pertinent for researchers who want to compare different *variable full control sets* (as in [30–32]), because any comparison of *control energy* across two different annotation maps needs to account for differences in those maps’ distributions. The simplest solution to this problem is to take the rank of each annotation map and then rescale the ranks to be between 1 and 2. `normalize_weights` performs this normalization:

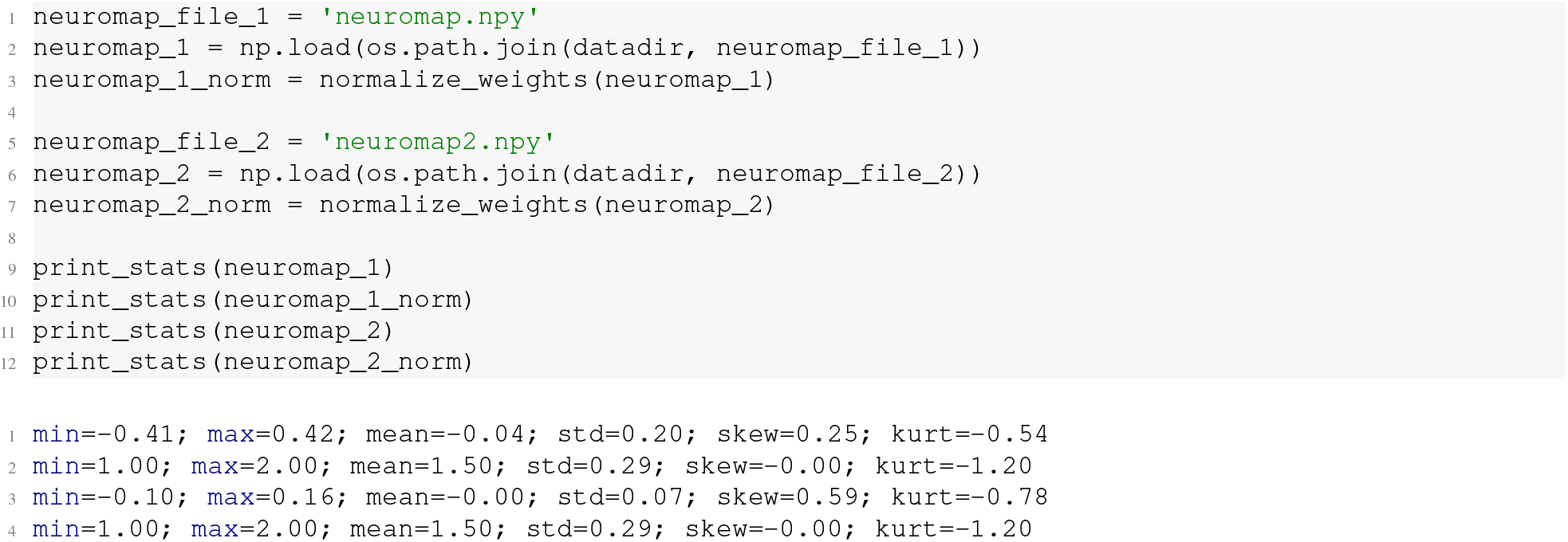

Once annotation maps have been normalized in this manner, differences in energy can only be attributed to differences in the (rank) spatial patterning between maps. This independence occurs because both annotation maps now conform to a uniform distribution. Note, energy derived from this normalization approach will still be lower than our *uniform full control set*.

3D) *Define a control task:* ***data-driven variable*** *full control set*. Instead of assigning variable weights *a priori*, researchers may assign them in a data-driven manner. In our recent work, we developed an approach for achieving this goal using gradient descent (see [51] for more details). Briefly, starting with a *uniform full control set*, this approach involves perturbing control nodes’ weight one at a time by a constant arbitrary amount and measuring the corresponding change in energy. This process results in *N* estimates of *perturbed control energy* for a given state transition. As mentioned above, each of these perturbations will necessarily reduce *control energy* compared to the baseline *uniform full control set*, creating negative Δs. In turn, differences in Δ magnitude encode the relative importance of each node to completing a specific state transition; nodes with more negative energy Δs are more important:

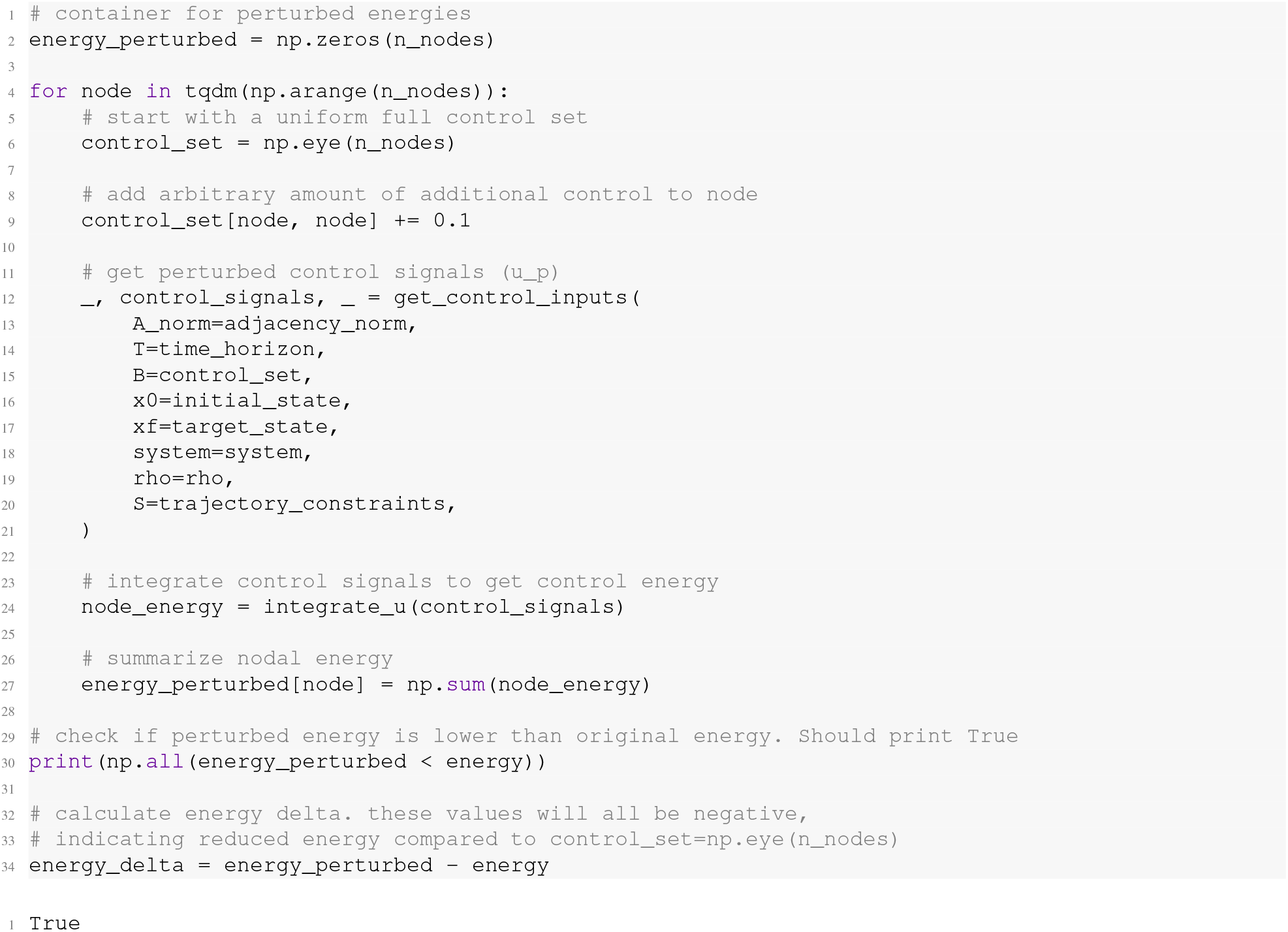

Once estimated, these Δs can be used as weights on *B* to obtain an optimized version of *control energy*. Note that here we draw a distinction between *optimal* and *optimized*. The former refers to constraining the magnitudes of ***x***(*t*) and ***u***(*t*) in the optimization problem, whereas the latter refers to finding the weights that create the most efficient transition, irrespective of this constraint:

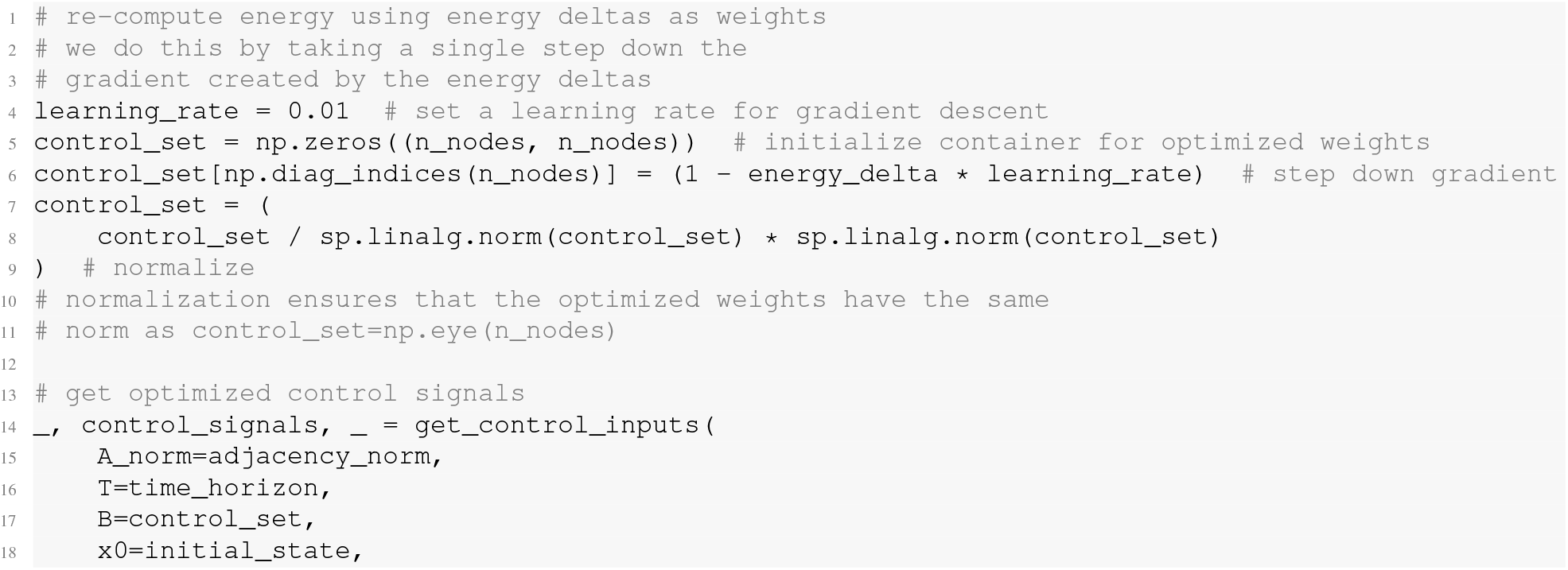
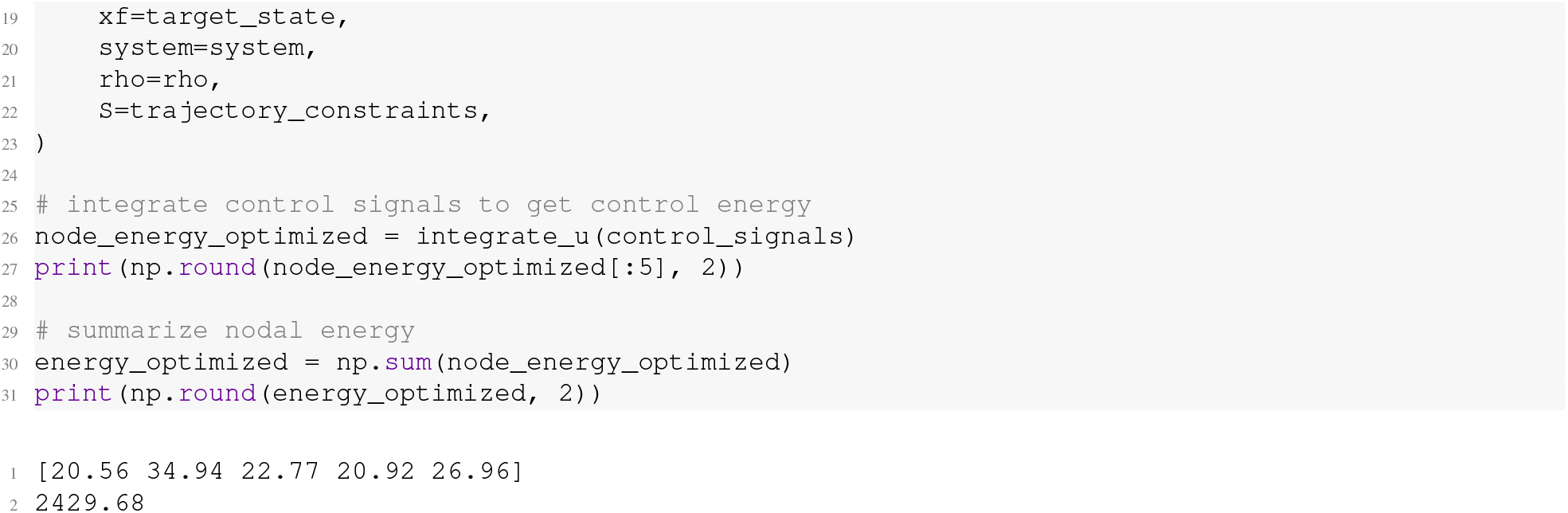

The above code uses `energy_delta` to define a gradient that we step down one time using a learning rate of 0.01. Stepping down this gradient yields a set of optimized weights, *B*_*o*_, that we then use to re-estimate *control energy*. As expected, this new estimate of energy (2429.68) is lower than the energy derived from our *uniform full control set* (2604.71). Thus, we have found a set of control weights that optimize (i.e., reduce) our *control energy* in a data-driven way. Additionally, setting up this algorithm using gradient descent allows researchers to optimize energy over multiple steps, wherein each new set of optimized weights is calculated from the previous set. In `nctpy` we include a *Python* class called `ComputeOptimizedControlEnergy` that wraps the above optimization steps and allows researchers to define their own learning rate and number of gradient steps:

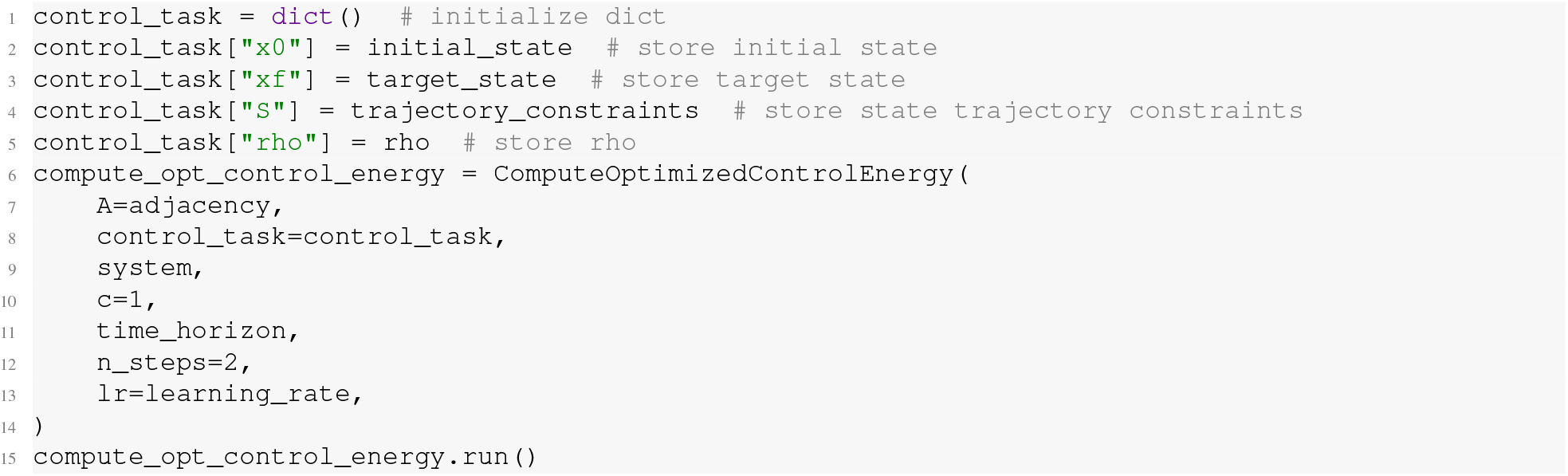

`ComputeOptimizedControlEnergy` is similar to `ComputeControlEnergy` but has some notable differences. Like `ComputeControlEnergy`, `ComputeOptimizedControlEnergy` will perform matrix normalization internally, thus we input *A* rather than *A*_*norm*_. Additionally, `ComputeOptimizedControlEnergy` requires that users specify the time system (`system=‘continuous’`), normalization constant (`c=1`), and time horizon (`T=1`) as input arguments. The differences are as follows. First, `ComputeOptimizedControlEnergy` only accepts a single control task dictionary, rather than a list of tasks. Second, `ComputeOptimizedControlEnergy` requires that users also specify the number of gradient steps (`n_steps=2`) and the learning rate (`lr=0.01`) as inputs. Once instantiated, `ComputeOptimizedControlEnergy.run()` will run the above optimization steps. Once completed, optimized energy at each gradient step will be stored in `ComputeControlEnergy.E_opt` as a vector of length *N*_*s*_, where *N*_*s*_ is the number of steps. The corresponding optimized control weights will be stored in `ComputeControlEnergy.B_opt` as an *N*_*s*_×*m* matrix.

#### Directed structural connectome

In Section IX A, we performed NCT analysis using an undirected structural connectome derived from the human brain. However, our protocol is designed to work with directed connectomes as well. Thus, as a final variation on Pathway A, we present results from a directed connectome obtained in the mouse brain. As discussed in Section VIII, our model assumes that *A*_*ij*_ encodes the edge connecting *node j to node i*. Provided that this assumption is met, Pathway A can be run without modification. Using the Allen Mouse Brain Connectivity Atlas [24, 25, 148], we extracted the ipsilateral directed connectivity from 43 regions in the mouse isocortex. These 43 regions were grouped into 6 systems: auditory, lateral, medial, prefrontal, somatomotor, and visual. Following [148], we combined the prefrontal and medial systems, as well as 3 regions from the somatomotor system, to create the mouse default mode system. Then, using `ComputeControlEnergy`, we estimated the energy required to transition from the default mode to the lateral, visual, and auditory systems and back again. Similar to the undirected human connectome (see Figure 5), this process yielded energy asymmetries (Figure 7A). We repeated this process using a symmetric version of the mouse connectome (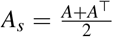; Figure 7B) and examined how energy asymmetries differed between the directed and undirected cases (Figure 7C); see here for *Python* code. For both the directed (Figure 7A) and undirected connectomes (Figure 7B), we found that *control energy* was lower when transitioning to the default mode compared to from the default mode. Critically, these observed energy asymmetries were larger for the directed connectome compared to the undirected connectome (Figure 7C); the lateral energy asymmetry was larger by 131 units (20% larger), the visual asymmetry was larger by 16 units (9% larger), and the auditory asymmetry was larger by 92 units (35% larger). Thus, the presence of directed edges increased the energy asymmetries observed in our model.

**FIG. 7.**
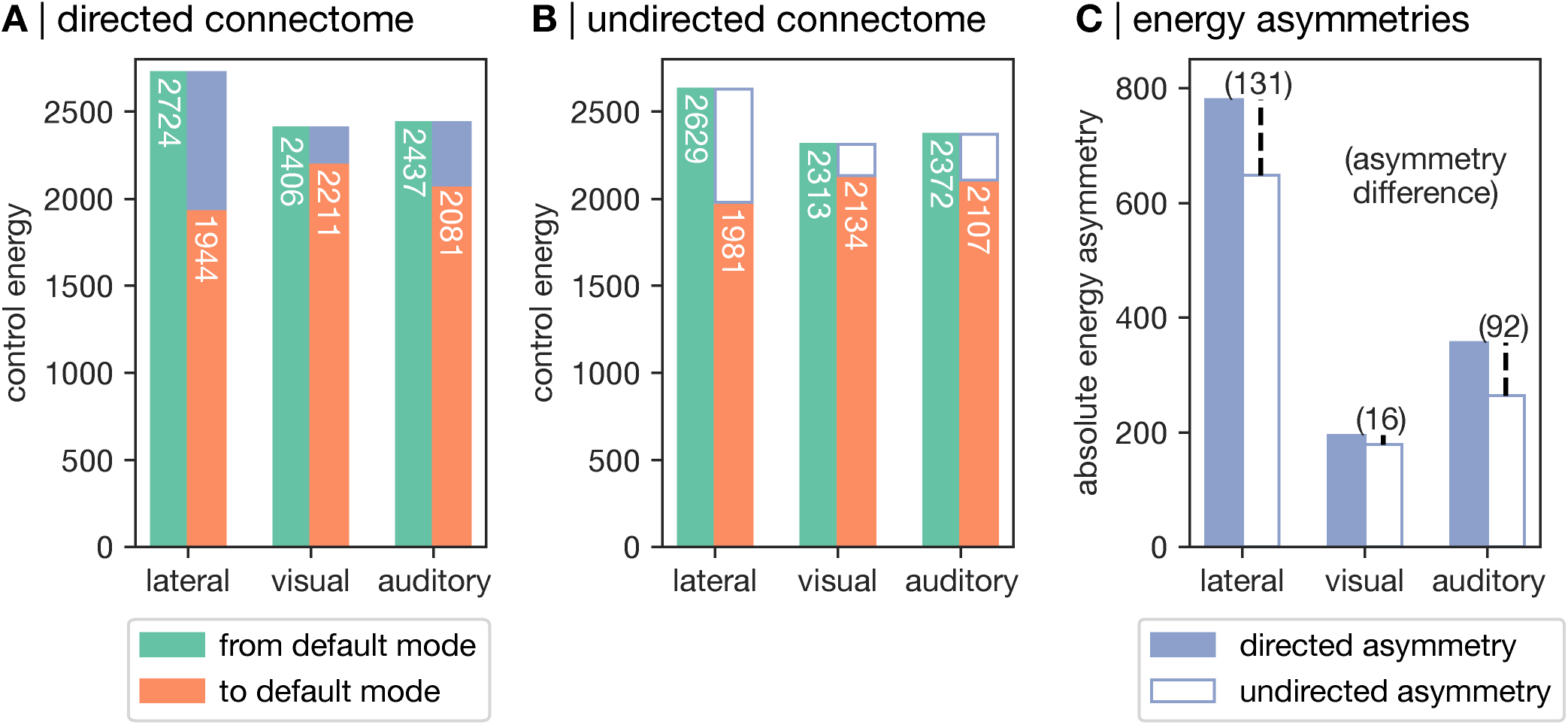
Energy asymmetries are larger for the directed than the undirected mouse connectome. Our protocol can be applied to connectomes with directed edges as well as undirected edges. In the directed case, our protocol assumes that *A*_*i j*_ encodes the edge connecting *node j to node i*. Here, we estimated *control energy* using the Allen Mouse Brain Connectivity Atlas. **A**, *Control energy* associated with transitioning from the default mode (DMN) to the lateral, visual, and auditory systems (green) and back again (orange) in the directed mouse connectome. *Control energy* associated with transitioning to the default mode was lower than the reverse direction, indicating a clear asymmetry (blue). **B**, *Control energy* estimated in the undirected mouse connectome. We recomputed energy using a symmetrized version of the mouse connectome (*As* = *A* + *A*^⊤^) and observed the same set of energy asymmetries. **C**, Differences in energy asymmetries between the directed and undirected mouse connectome. We observed that the size of the energy asymmetries were larger for the directed than the undirected mouse connectome.

### B. Protocol Pathway B: Average Controllability

Pathway A is the primary component of our protocol. Implementing these steps assumes that researchers are interested in studying a specific set of state transitions defined in accordance with their research questions and hypotheses. In the absence of such hypotheses, researchers may instead wish to examine nodes’ general capacity to control a broad range of unspecified state transitions. To support these types of hypotheses, we present a complementary pathway to our protocol that yields estimates of *average controllability*, where higher values indicate that a region is better positioned in the network to control dynamics (see section II):

1. **Compute average controllability** (Timing: discrete time, *<* 1 second for 200 nodes; continuous time, 10 −20 seconds for 200 nodes).

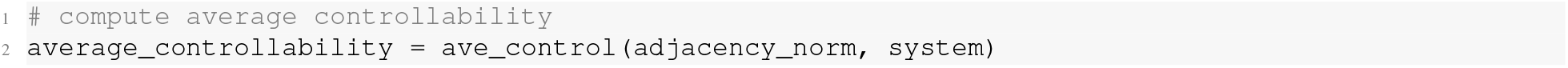

`average_controllability` will be a vector containing the *average controllability* of each node of the system.

2 **Visualize average controllability**. As *average controllability* is a regional metric, we can simply plot its distribution of values on the surface of the cortex (Figure 8).

**FIG. 8.**
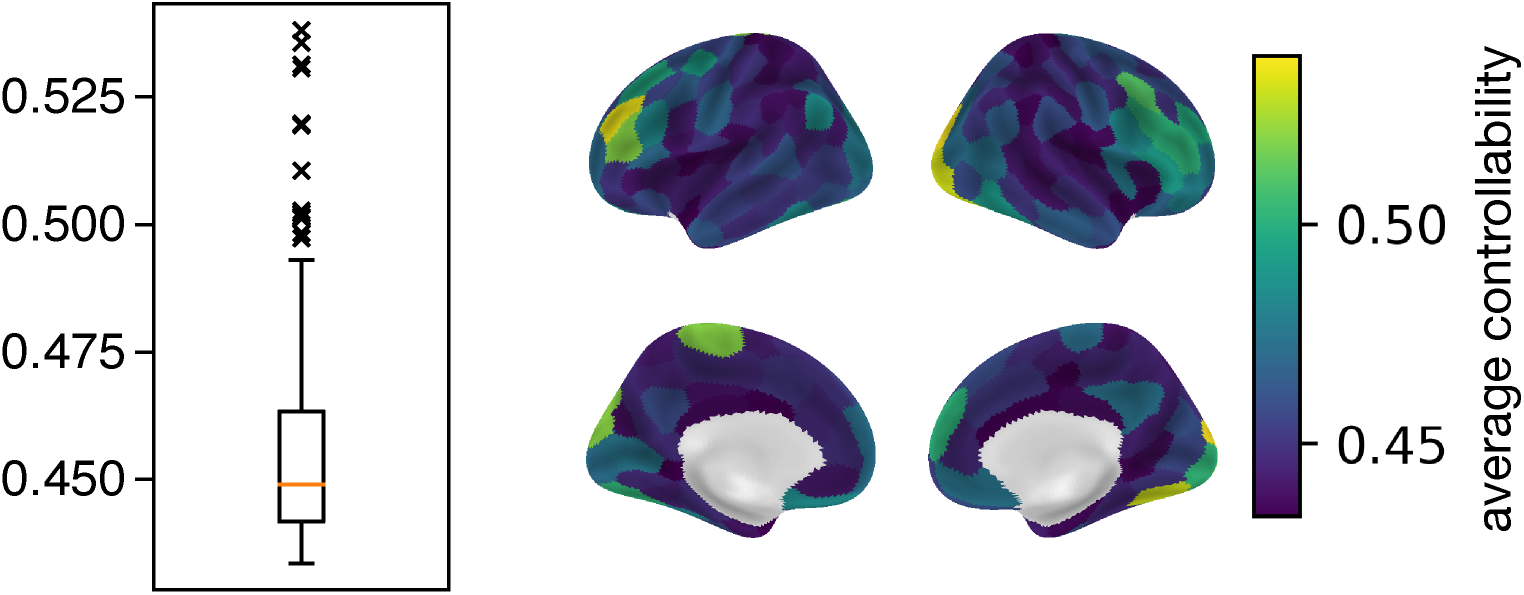
Average controllability. Each system node receives an impulse of equal magnitude. Nodes with higher *average controllability* are able to broadcast that impulse throughout the system to a greater extent compared to nodes with lower *average controllability*. Thus, nodes with high *average controllability* are better positioned within the network to control dynamics. Distribution of *average controllability* values are displayed using a box plot (left) as well as projected onto the cortical surface (right). In the box plot, the orange line represents the median, the box spans the middle 50% of the data, the whiskers span 1.5 times the interquartile range on either side, and the crosses represent outliers beyond this limit.

## X. ANTICIPATED RESULTS

The final outputs of our protocol will depend on whether researchers choose to follow Pathway A (see Section IX A) or Pathway B (see Section IX B). For the former, the output will be one estimate of *control energy* per control task, or one estimate per brain region per task if energy was not summarized across regions. This value will be positive and can be thought of as the amount of effort the model has to exert in order to complete a specific control task; higher energy corresponds to greater effort. For the latter, output will be one estimate of *average controllability* per brain region; a regional map of control over system dynamics. These regional values will also be positive. Greater *average controllability* indicates that regions are better positioned within the network’s topology to broadcast an impulse, and as such may better orchestrate control of brain dynamics.

What can researchers do with these outputs? The answers to this question are diverse and depend heavily on the researcher’s goals. As we discussed in Section III, we have used NCT to investigate a range of research questions that spanned from examining the influence of topology [10, 49, 50], to predicting state transitions observed in functional data [14, 73], to studying individual differences, including psychosis symptoms [69], executive function [71], and sex effects [19]. Providing detailed guidance on each of these applications is beyond the scope of this protocol. However, to conclude this protocol, we outline the use of null network models as an initial analysis that we believe is an essential step irrespective of researchers’ study goals.

### 1. Null network models

Null models allow researchers to examine the extent to which different aspects of topology explain NCT model outputs. As discussed in Ref. [149], these null models take different forms, including edge rewiring, generative models of surrogate networks, and spatially-preserved node permutation. Of these different forms, the appropriate choice will depend on the research question. Here, in order to understand the extent to which topology informs *control energy*, we focus on null models that rewire an empirical adjacency matrix (see Ref. [69] for an example of spatially-preserved node permutation used to compare maps of *average controllability* with other properties of network topology). In this case, a null network model involves randomly swapping the edges of the adjacency matrix *n* times subject to certain constraints—for example, while preserving the spatial embedding of the nodes as well as the degree or strength distribution [149, 150]—and recalculating energy upon each rewired matrix. This process generates an empirical null distribution that observed *control energy* can be compared against. For instance, *control energy* that is lower than expected under the null suggests that NCT was able to leverage properties of network topology, beyond those preserved by the null model, to complete a given state transition. In turn, by deploying a range of null models that each preserve different topological properties, researchers can systematically probe the aspects of topology that explain their observed outputs.

Using the undirected human connectome, we provide an example of the above approach using two of our binary state transitions: the default mode to visual transition and the default mode to ventral attention (VAN) transition. We chose these transitions as they represent two control tasks with strong but opposing energy asymmetries (see Figure 5). For each transition, we recompute the *control energy* for each direction under two null models. The first preserves the spatial embedding of the nodes as well as the strength distribution (strength-preserving). The second preserves spatial embedding and the strength sequence of the nodes (sequence-preserving). That is, the sequence-preserving null preserves the strength of each node as it was in the original connectome. By contrast, the strength-preserving null only preserves the distribution of strength across the network; the strength of each node is allowed to change. The sequence-preserving null is a more stringent test than the strength-preserving null as it preserves how strength is embedded in the connectome [150]. We begin by loading the coordinates of our nodes in 3 dimensions:

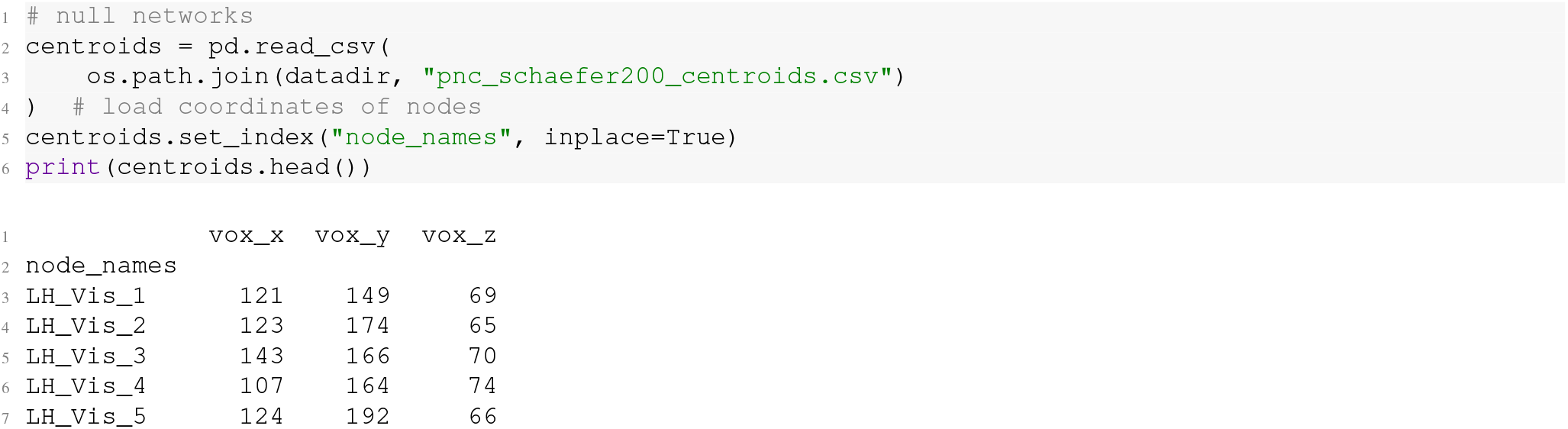

Then, we use those coordinates to define a distance matrix that encodes the physical distance between node pairs:

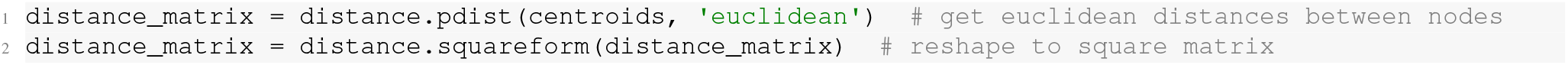

Finally, we define our control task and compute our nulls using the included function, `geomsurr` [150]:

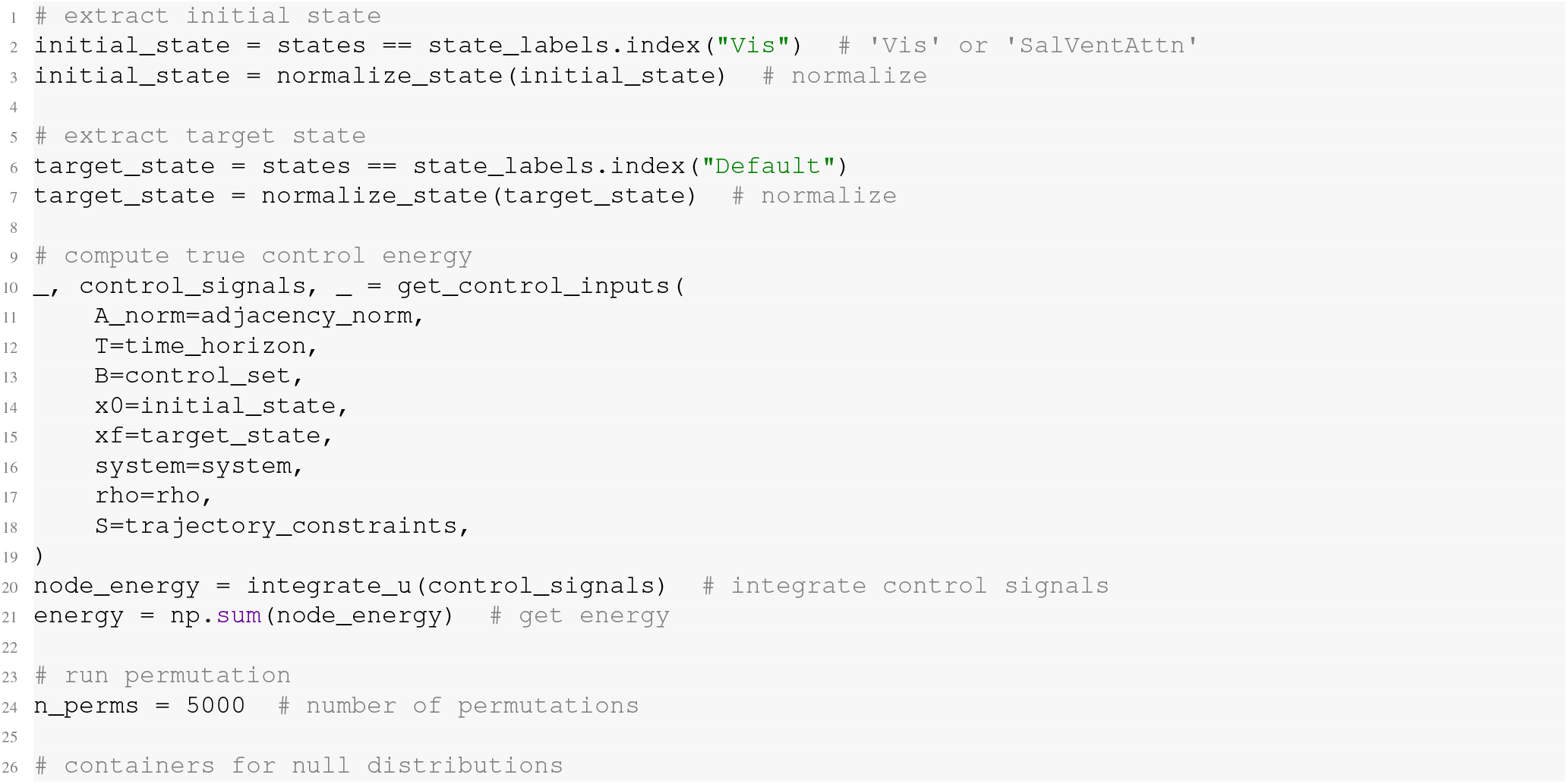
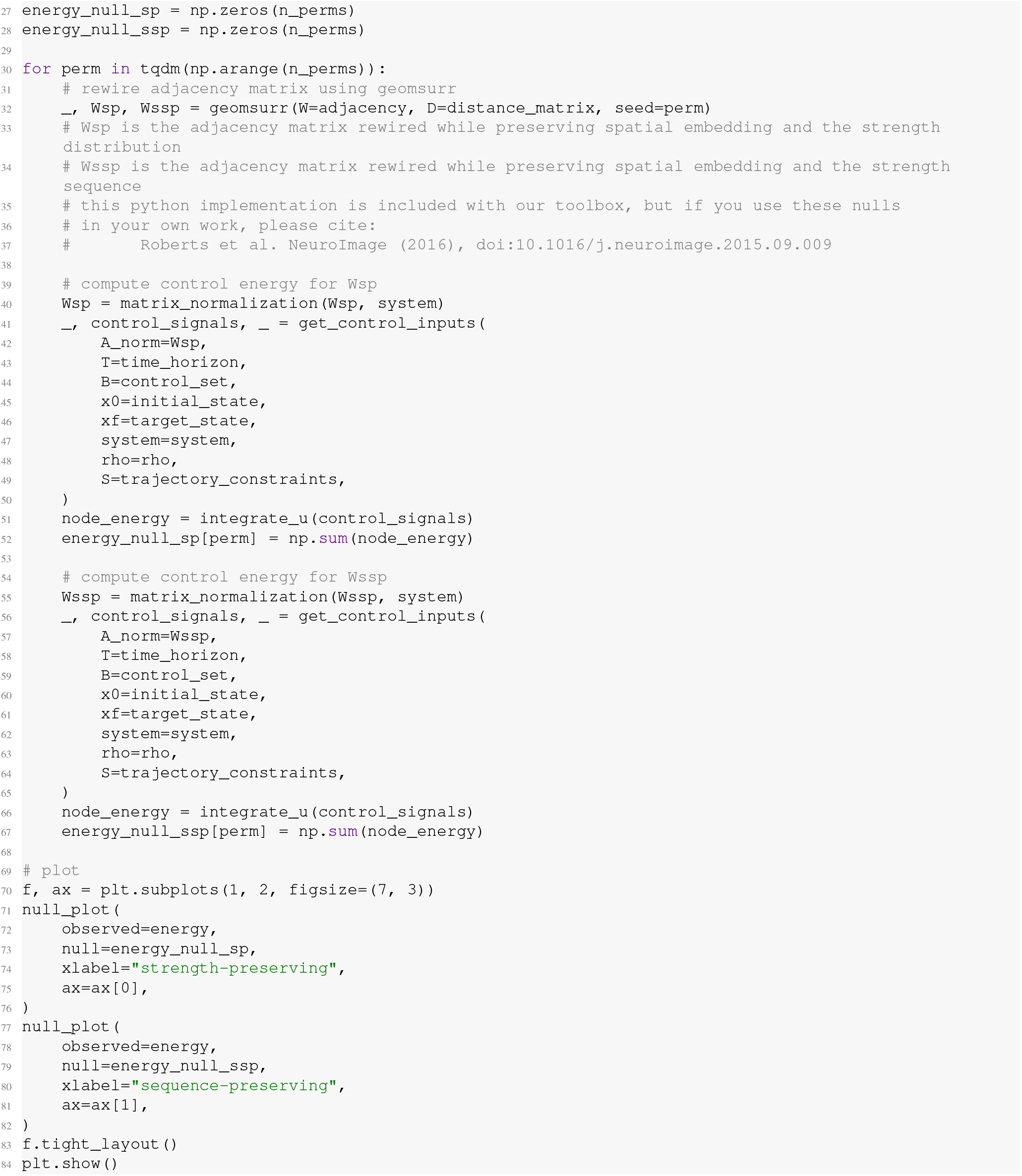

Figure 9 displays the energy associated with transitioning from the visual cortex to the DMN (Figure 9A) and back again (Figure 9B), as well from the VAN to the DMN (Figure 9C) and back again (Figure 9D). In each panel of Figure 9, the strength-preserving null is shown on the left and the sequence-preserving null is shown on the right. These results provide several insights. First, as mentioned above, both transitions show energy asymmetries but in opposite directions. The energy associated with transitioning from visual cortex to the DMN is larger (energy = 2605) compared to the reverse direction (energy = 1947). By contrast, the energy associated with transitioning from the VAN to the DMN is lower (energy = 2218) compared to the reverse direction (energy = 2601). Note that the former result represents an exception to the general finding that energy is lower when transitioning to the DMN than from it (see Figure 5). This pattern of findings is consistent with what we observe in the mouse connectome (Figure 7).

**FIG. 9.**
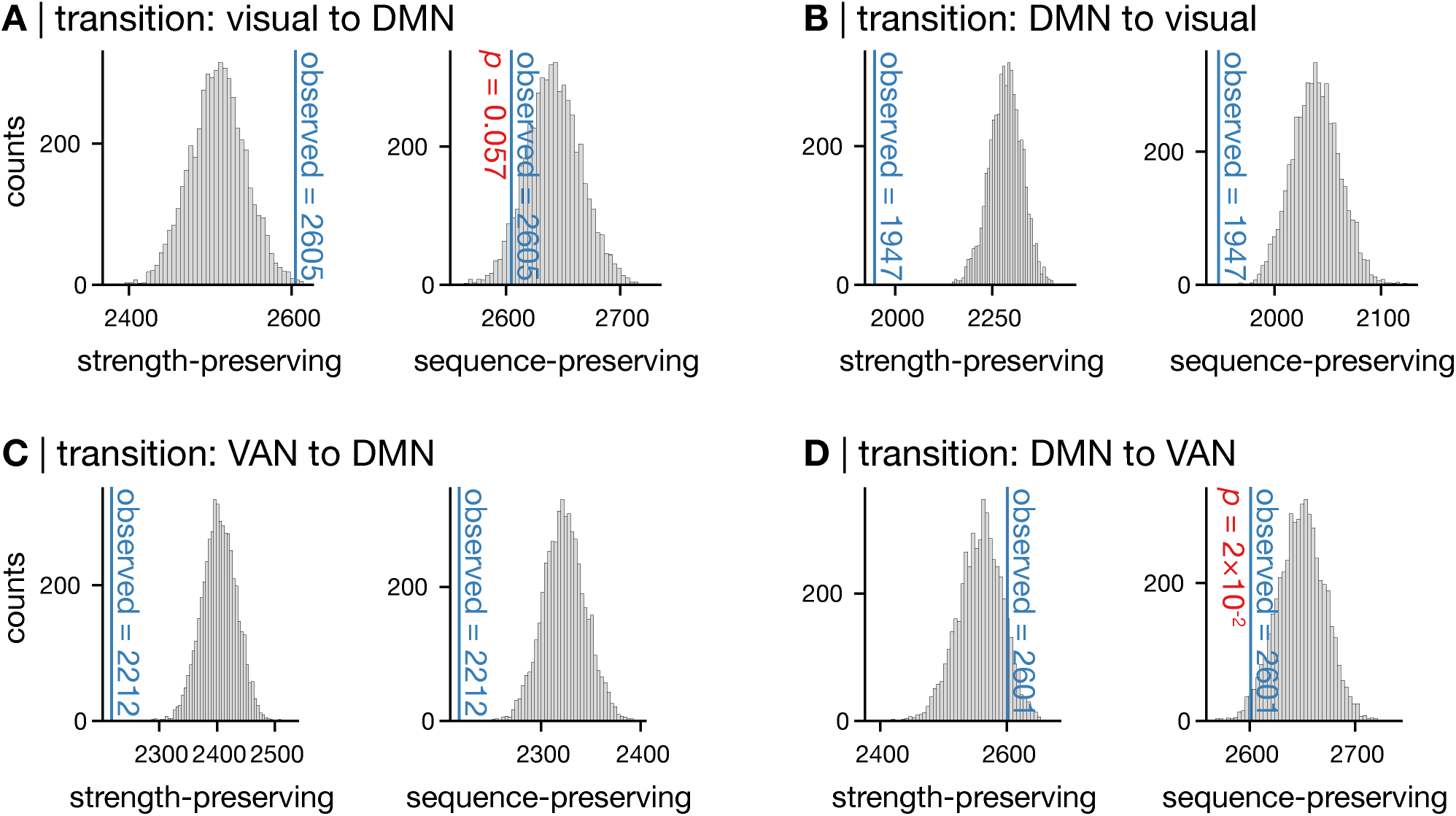
Null network models uncover the topological properties that are important for control energy. For a given state transition, we recompute *control energy* 5,000 times. Each time we randomly rewire the edges of the adjacency matrix subject to certain constraints. This process generates an empirical null distribution for *control energy*. Here, we generate two null distributions per state transition; one that preserves the spatial embedding of system nodes as well as the strength distribution (strength-preserving), and another that preserves spatial embedding and the strength sequence of the nodes (sequence-preserving). We computed these nulls for the visual-to-DMN transition (**A**), the DMN-to-visual transition (**B**), the VAN-to-DMN transition (**C**), and the DMN-to-VAN transition (**D**). Collectively, these results demonstrate that *control energy* for some transitions is likely driven by strength (e.g., visual-to-DMN and DMN-to-VAN) while others may be driven by higher-order topology (e.g., DMN-to-visual and VAN-to-DMN). See main text for extended discussion.

Second, when we preserve the strength distribution in our null model, we observe larger-than-expected energy when transitioning from the visual cortex to the DMN (Figure 9A, left). This result appears counter intuitive until we consider the differences in strength between these brain states. In the structural connectome used here, the mean strength of the nodes within the visual state is 67,773, while the mean strength of the nodes in the DMN is 59,455, and the mean strength of the remaining nodes is 53,761. Thus, the connectivity strength between the visual state and the rest of the brain is higher than the connectivity strength between the DMN and the rest of the brain. When we preserve only the strength distribution in the null, this difference in strength is not maintained, which results in relatively high-strength nodes being redistributed throughout the brain. In turn, this redistribution results in reduced *control energy* in the null distribution for the visual-to-DMN transition. This finding suggests that the high strength nodes of the visual system broadcast activity in a way that necessitates high amounts of *control energy* to guide those dynamics toward the DMN. By contrast, when we preserve the strength sequence (Figure 9A, right) we also preserve the between-state difference in strength, which yields a null that is much closer to the observed energy. Together, the results in Figure 9A demonstrate that differences in connectivity strength between brain states drives the energy observed for the visual-to-DMN transition.

Third, the above line of reasoning does not hold when we consider the transition from DMN back to visual cortex (Figure 9B). Here, we observe lower-than-expected energy under both the strength-preserving and the sequence-preserving nulls. This result demonstrates that the aforementioned difference in strength between the visual state and the DMN is not what drives observed energy. In turn, this result suggests that higher-order topological properties of the connectome may support the efficient completion of the DMN-to-visual transition. Finally, Figure 9C and D show that all of the above results and interpretations vary as a function of states.

The above findings—that the spatial embedding of node strength drives energy for one transition direction but not the other—underscores the utility of probing NCT outputs using null network models. That is, through comparing a pair of null network models, we obtained evidence for how certain aspects of network topology (i.e., strength) contribute to different state transitions.

Null network models can also be applied to *average controllability*:

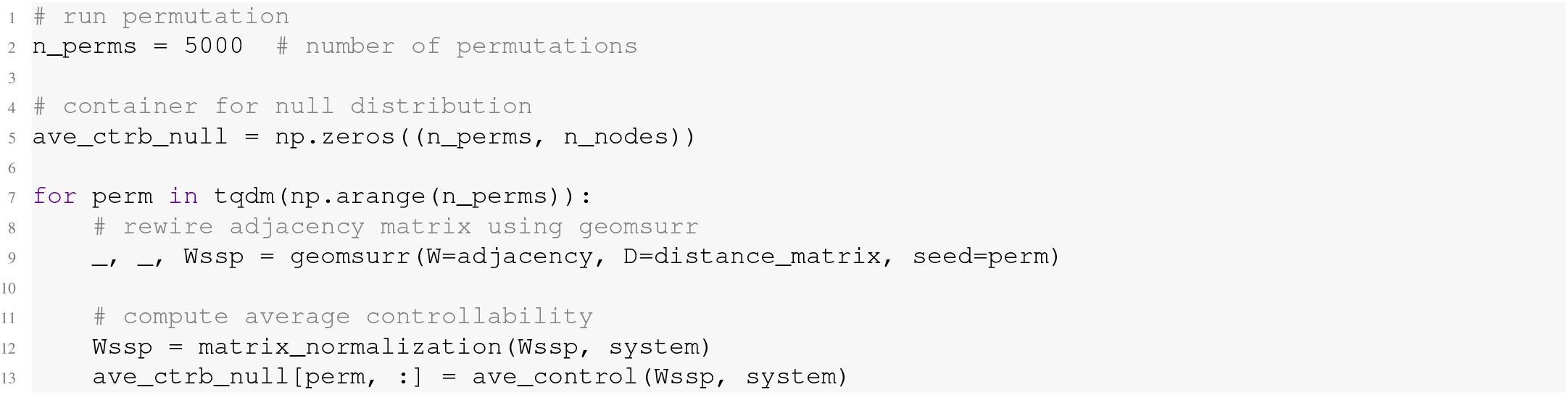

The above code will generate an empirical null distribution for *average controllability* at each system node. In turn, this procedure will yield *N* null distributions for a given null network model, the visualization of which is impractical. As such, researchers may instead assign *p*-values to the observed *average controllability* values using `get_null_p`:

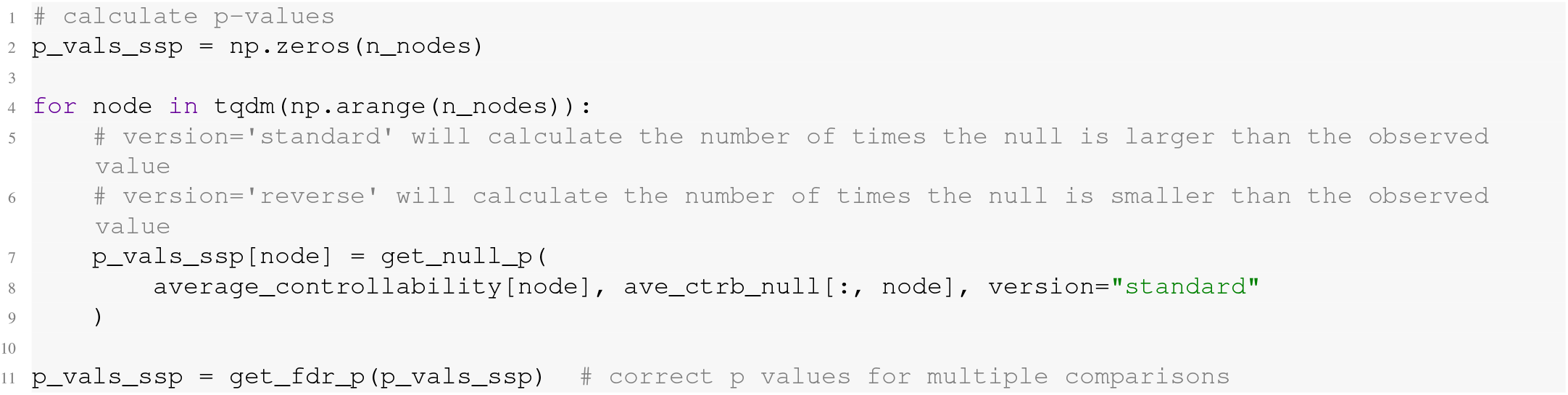

The above code will yield FDR-corrected *p*-values denoting the proportion of times that a nodes’ *average controllability* was larger than expected under the null. As touched upon above, this result only tells half the story, and there may be reasons for *average controllability* to be smaller than expected under the null. As such, we recommend running both `version=‘standard’` and `version=‘reverse’` to test both tails of the null distribution.

## XI. TIMING

As noted throughout the protocol, the timing of each step is relatively short, often not exceeding 1 second per step. We note two clarifications. First, these time estimates are only for a single execution of each step as shown in the protocol. In reality, these steps will likely need to be executed many times over to achieve researchers’ goals. For example, a given study may need to compute control energy for multiple control tasks across multiple subjects, which will increase run time. This time will increase further if null network models are used, wherein each step may be run thousands of times for a single control task. However, in these instances, protocol steps can be trivially parallelized using High Performance Computing (HPC), which will reduce run time. Second, timing will vary as a function of researchers’ data processing. For example, in this protocol, we performed analysis on a structural connectome comprising 200 nodes. Increasing parcellation resolution will increase run time.

## Funding

National Institute of Mental Health grant K99MH127296 (LP). The content is solely the responsibility of the authors and does not necessarily represent the official views of the National Institutes of Health.

NARSAD Young Investigator Grant 28995 from the Brain & Behavior Research Foundation (LP)

National Institute of Mental Health grant R21MH106799 (DSB and TDS)

National Institute of Mental Health grant R01MH113550 (DSB and TDS)

National Institute of Mental Health grant RF1MH116920 (DSB and TDS)

Swartz Foundation (DSB)

John D. and Catherine T. MacArthur Foundation (DSB)

National Institute of Mental Health grant R01MH120482 (TDS)

National Institute of Mental Health grant R01MH107703 (TDS)

National Institute of Mental Health grant R01MH112847 (TDS and RTS)

National Institute of Mental Health grant R37MH125829 (TDS)

National Institute of Mental Health grant R01EB022573 (TDS)

National Institute of Mental Health grant R01MH107235 (RCG)

National Institute of Mental Health grant R01MH119219 (RCG and REG)

Penn-CHOP Lifespan Brain Institute

National Science Foundation grant DGE-1321851 (JZK)

National Institute of Mental Health grant RC2MH089983 (Philadelphia Neurodevelopmental Cohort)

National Institute of Mental Health grant RC2MH089924 (Philadelphia Neurodevelopmental Cohort)

## Author contributions

Conceptualization: L.P., J.Z.K., T.D.S., and D.S.B.

Methodology: L.P., J.Z.K., J.S., and D.S.B.

Software: L.P., J.Z.K., and J.S.

Formal analysis: L.P., and J.Z.K.

Visualization: L.P., and J.Z.K.

Data curation: J.K.B., M.C., S.C., R.E.G., R.C.G., R.T.S., D.Z., and T.D.S.

Writing—original draft: L.P., and J.Z.K.

Writing—reviewing and editing: L.P., J.Z.K, J.S., J.K.B., M.C., S.C., R.E.G., R.C.G., F.P., R.T.S., D.Z., T.D.S, and D.S.B.

## Competing interests

R.T.S. receives consulting compensation from Octave Bioscience and compensation for reviewership duties from the American Medical Association. All other authors declare no competing interests.

## Data availability

The PNC data are publicly available in the Database of Genotypes and Phenotypes: accession number: phs00607.v3.p2; https://www.ncbi.nlm.nih.gov/projects/gap/cgi-bin/study.cgi?study_id=phs000607.v3.p2

## Code availability

All analysis code is freely available at https://github.com/BassettLab/nctpy/

## XII. EXTENDED DATA

**FIG. S1.**
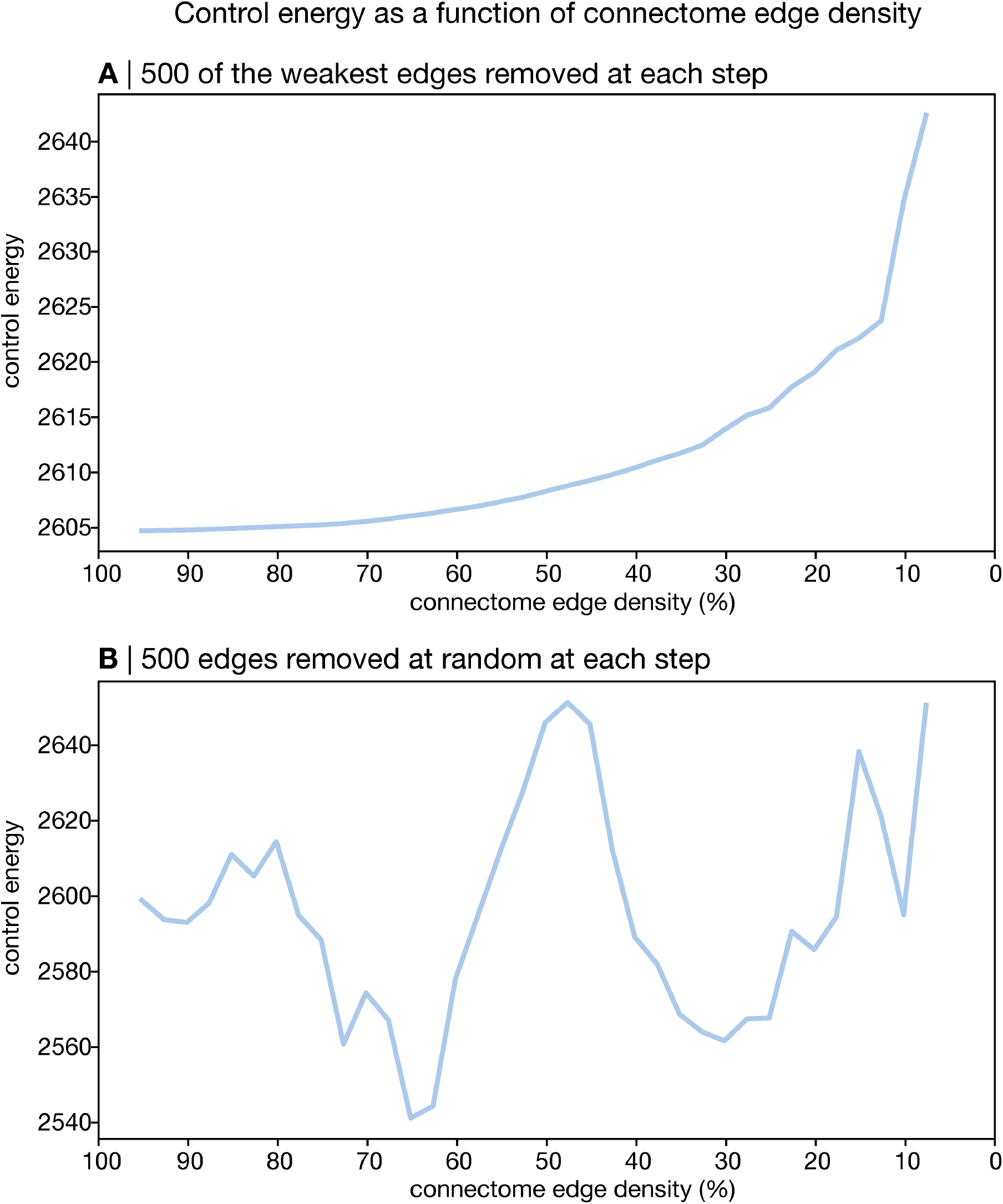
Control energy as a function of connectome edge density. In the undirected human connectome, we iteratively set 500 of the weakest edges (**A**) or 500 random edges (**B**) to 0, stopping once edge density fell below 10%. *Control energy* was recomputed at each iteration and is shown here. This plot illustrates the fact that energy varies as a function of edge density and is primarily driven by the strongest edges in the network.

**FIG. S2.**
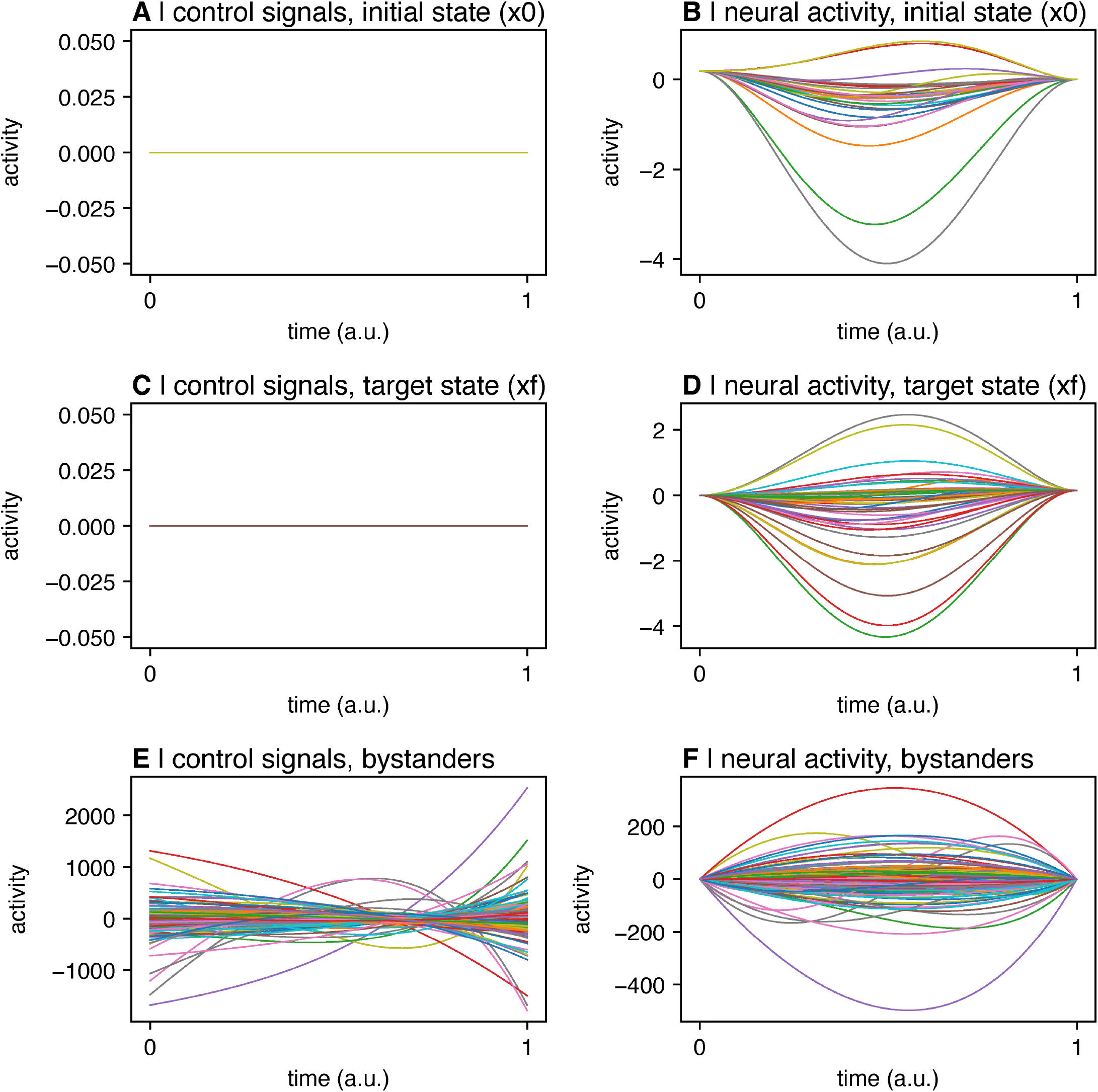
Control signals and state trajectory. *Only bystanders are set as control nodes*. T=1. Initial state = visual system. Target state = default mode system. Inversion error = 2.09 × 10^*−*10^. Reconstruction error = 1.75 × 10^*−*8^. Energy = 6.64 × 10^9^.

**FIG. S3.**
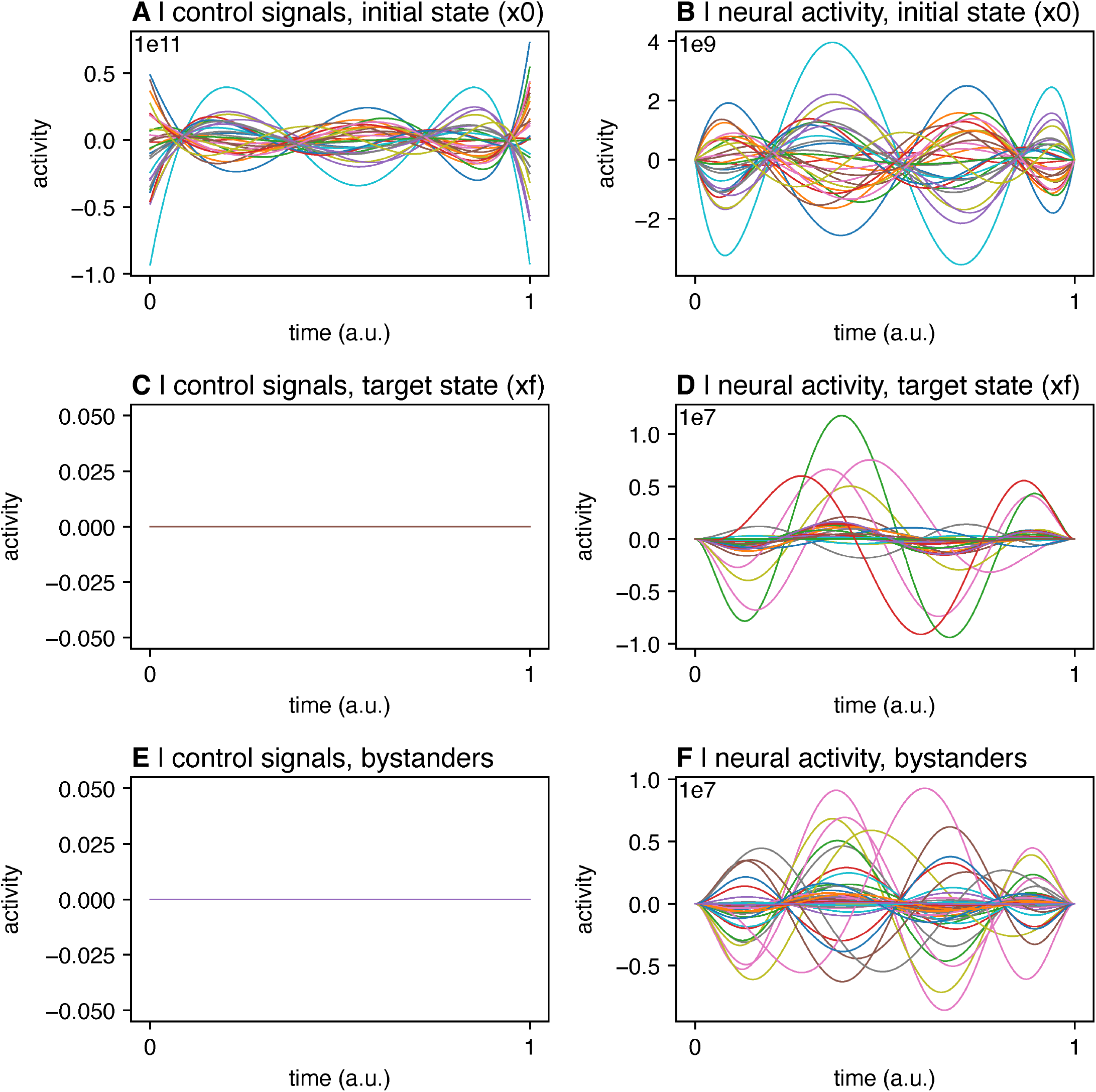
Control signals and state trajectory. *Only nodes in the initial state are set as control nodes*. T=1. Initial state = visual system. Target state = default mode system. Inversion error = 1.37 ×10^3^. Reconstruction error = 2.04 ×10^5^. Energy = 3.68 ×10^24^. Note that this state transition does not complete successfully.

**FIG. S4.**
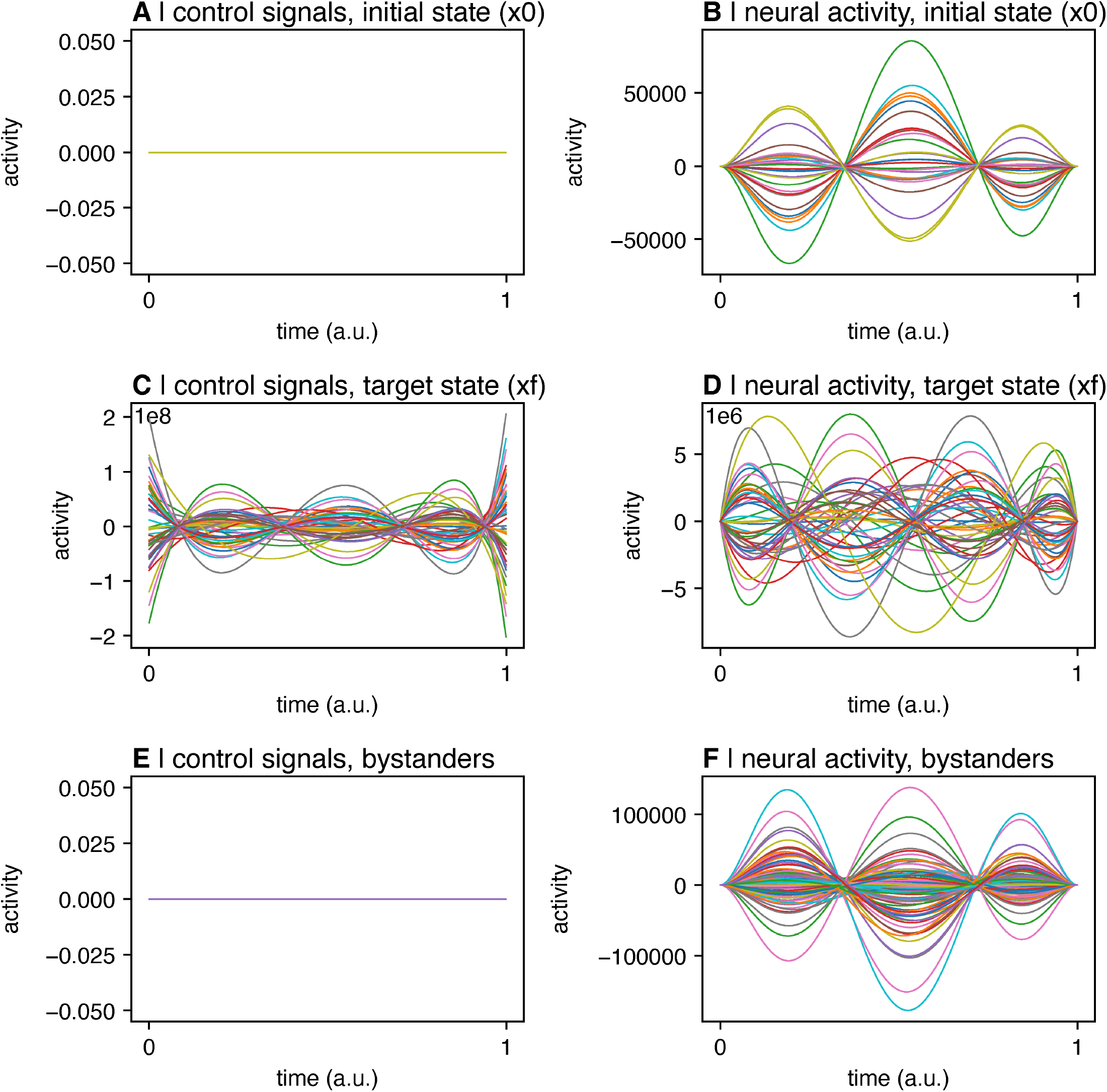
Control signals and state trajectory. *Only nodes in the target state are set as control nodes*. T=1. Initial state = visual system. Target state = default mode system. Inversion error = 1.50. Reconstruction error = 1.12 ×10^2^. Energy = 3.38 ×10^19^. Note that this state transition does not complete successfully.

**FIG. S5.**
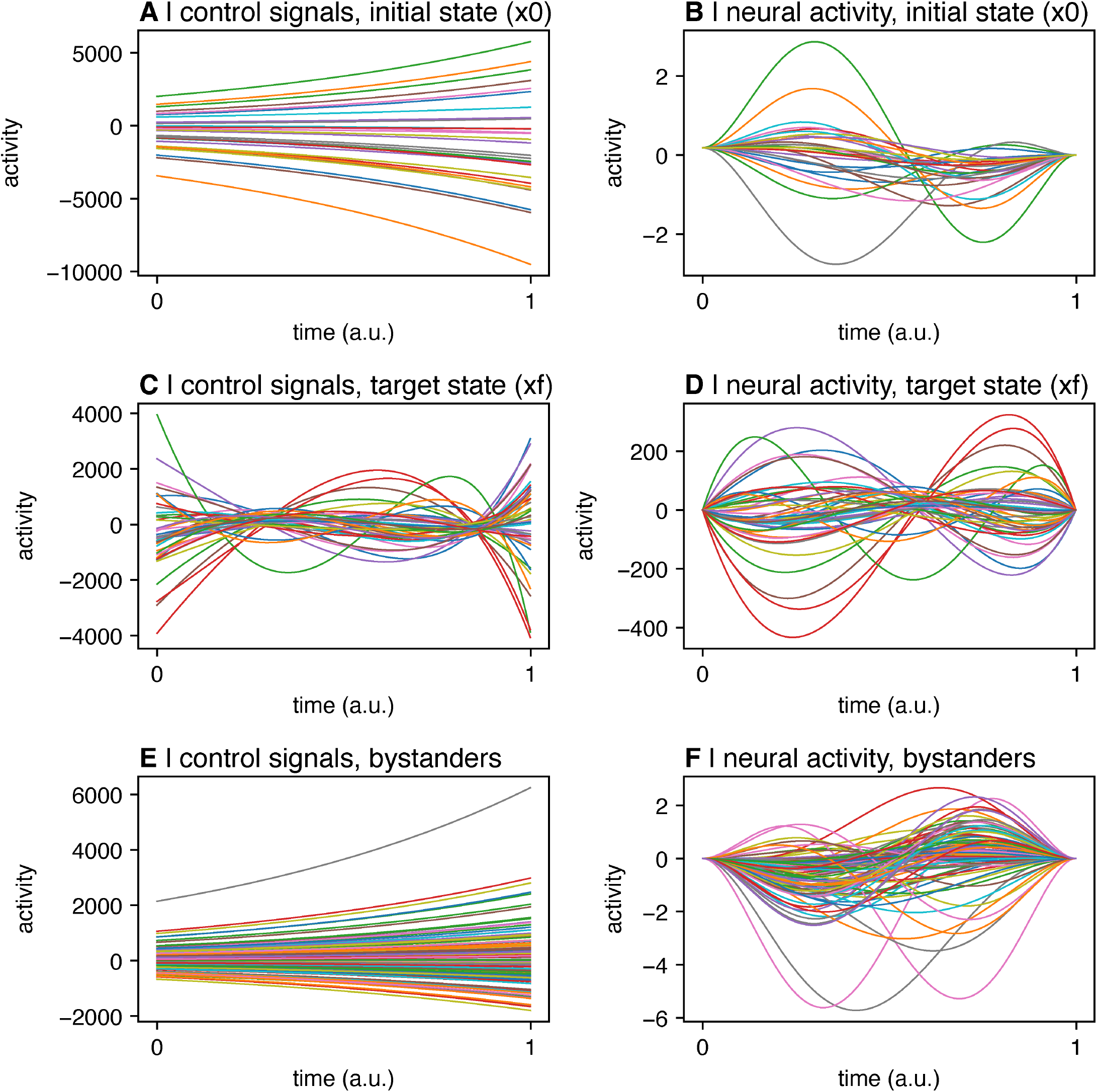
Control signals and state trajectory. *Nodes in the target state are set as control nodes with control weights of 1, whereas the remaining nodes are given a small amount of control (*1 ×10^*−*5^). T=1. Initial state = visual system. Target state = default mode system. Inversion error = 2.00 ×10^*−*8^. Reconstruction error = 1.16 ×10^*−*6^. Energy = 2.31 ×10^11^. Unlike the scenario depicted in Figure S4, this state transition completes successfully.

**FIG. S6.**
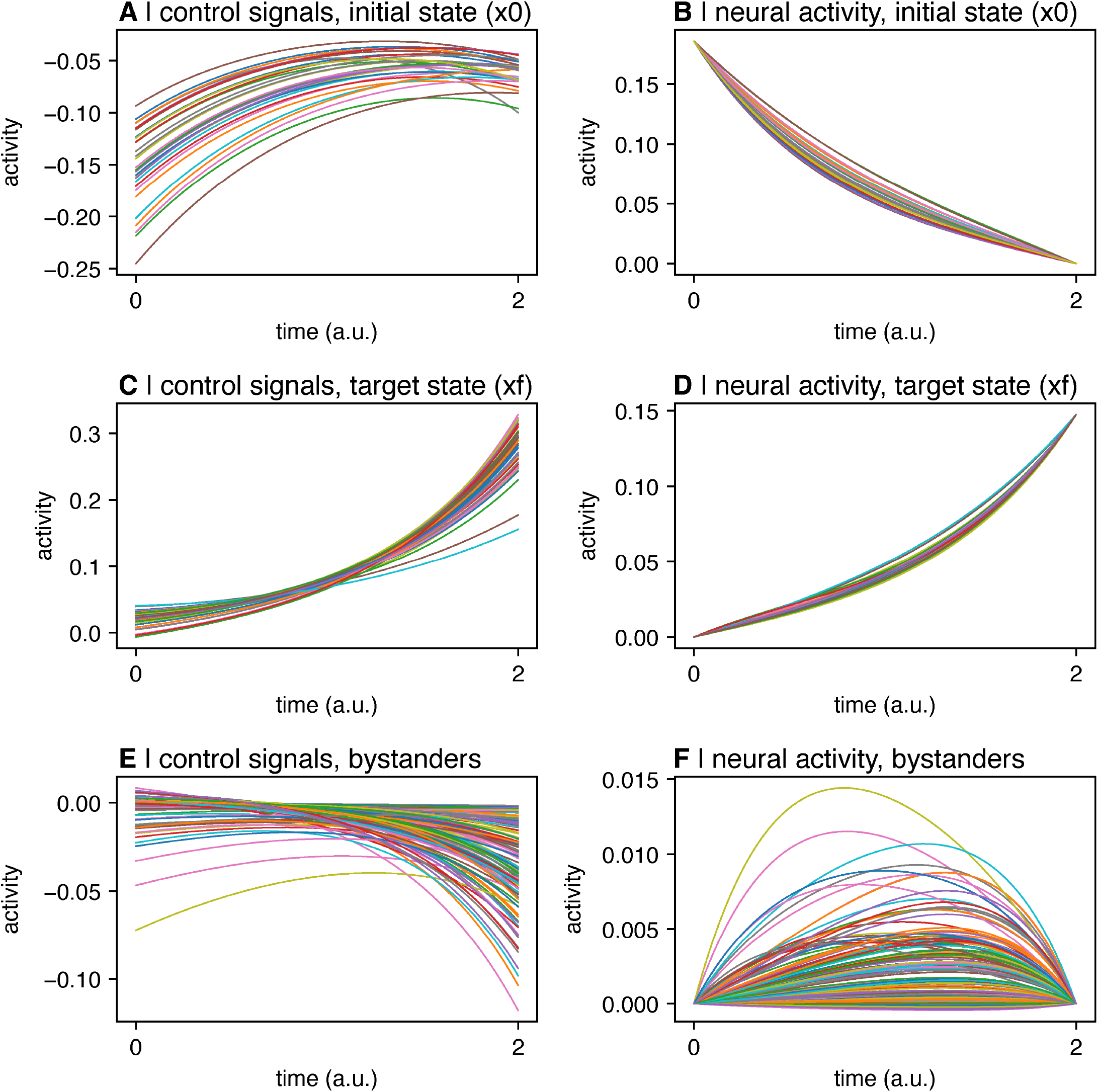
Control signals and state trajectory. Uniform full control set. *T=2*. Initial state = visual system. Target state = default mode system. Inversion error = 3.24 × 10^*−*15^. Reconstruction error = 1.55 × 10^*−*13^. Energy = 1904.

**FIG. S7.**
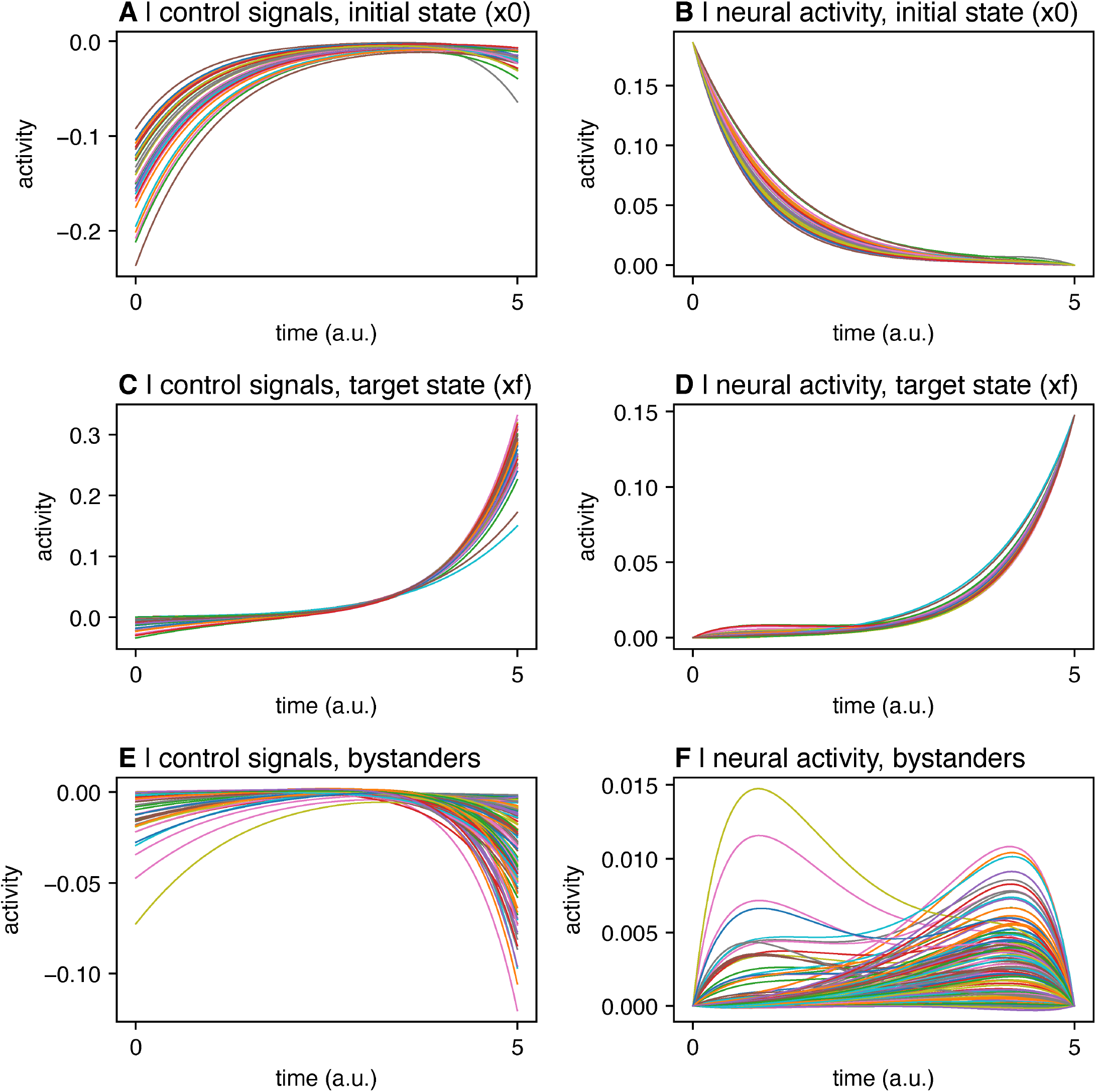
Control signals and state trajectory. Uniform full control set. *T=5*. Initial state = visual system. Target state = default mode system. Inversion error = 1.33 × 10^*−*13^. Reconstruction error = 1.03 × 10^*−*11^. Energy = 1797.

**FIG. S8.**
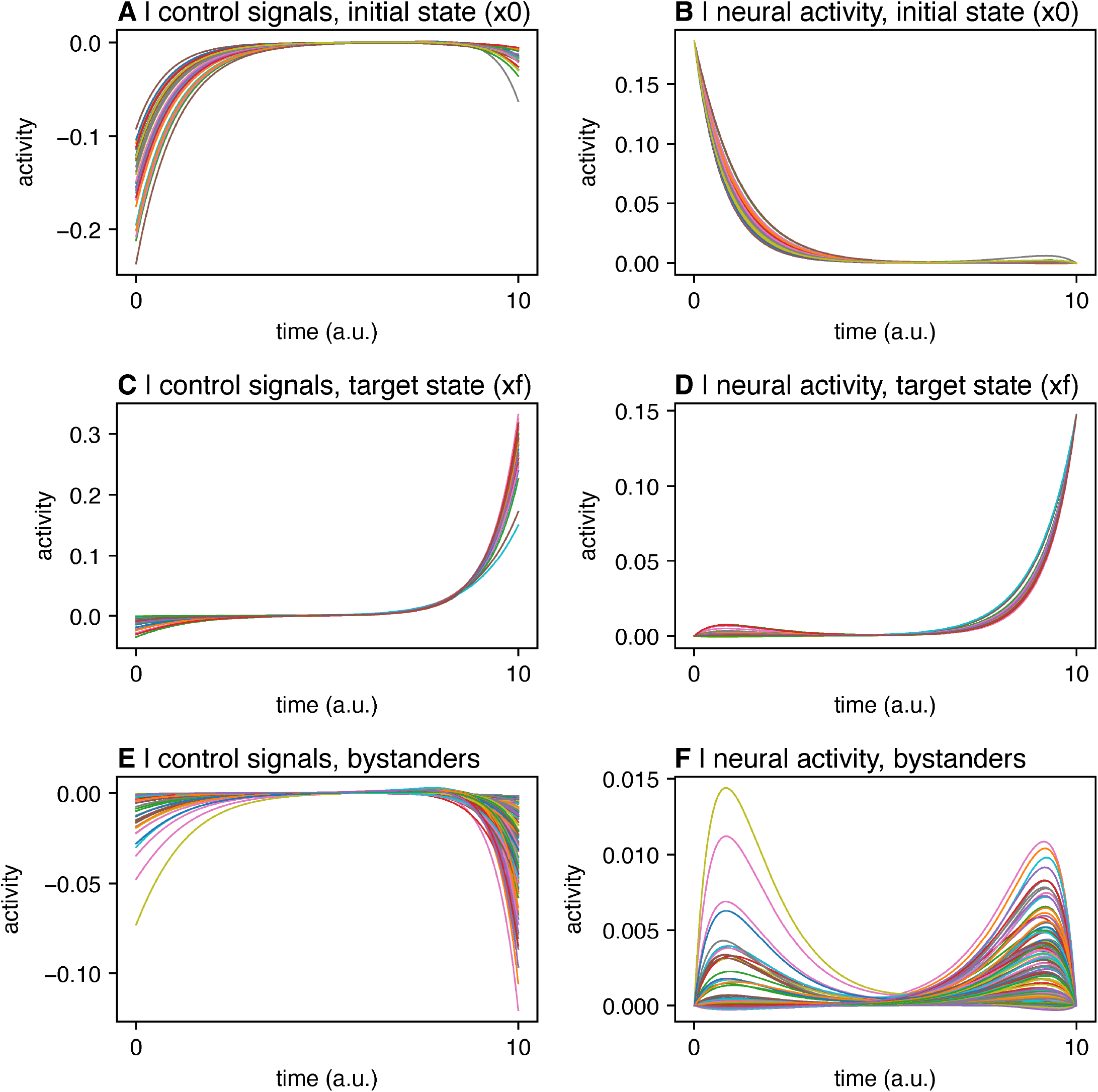
Control signals and state trajectory. Uniform full control set. *T=10*. Initial state = visual system. Target state = default mode system. Inversion error = 1.48 × 10^*−*10^. Reconstruction error = 1.71 × 10^*−*8^. Energy = 1801.

**FIG. S9.**
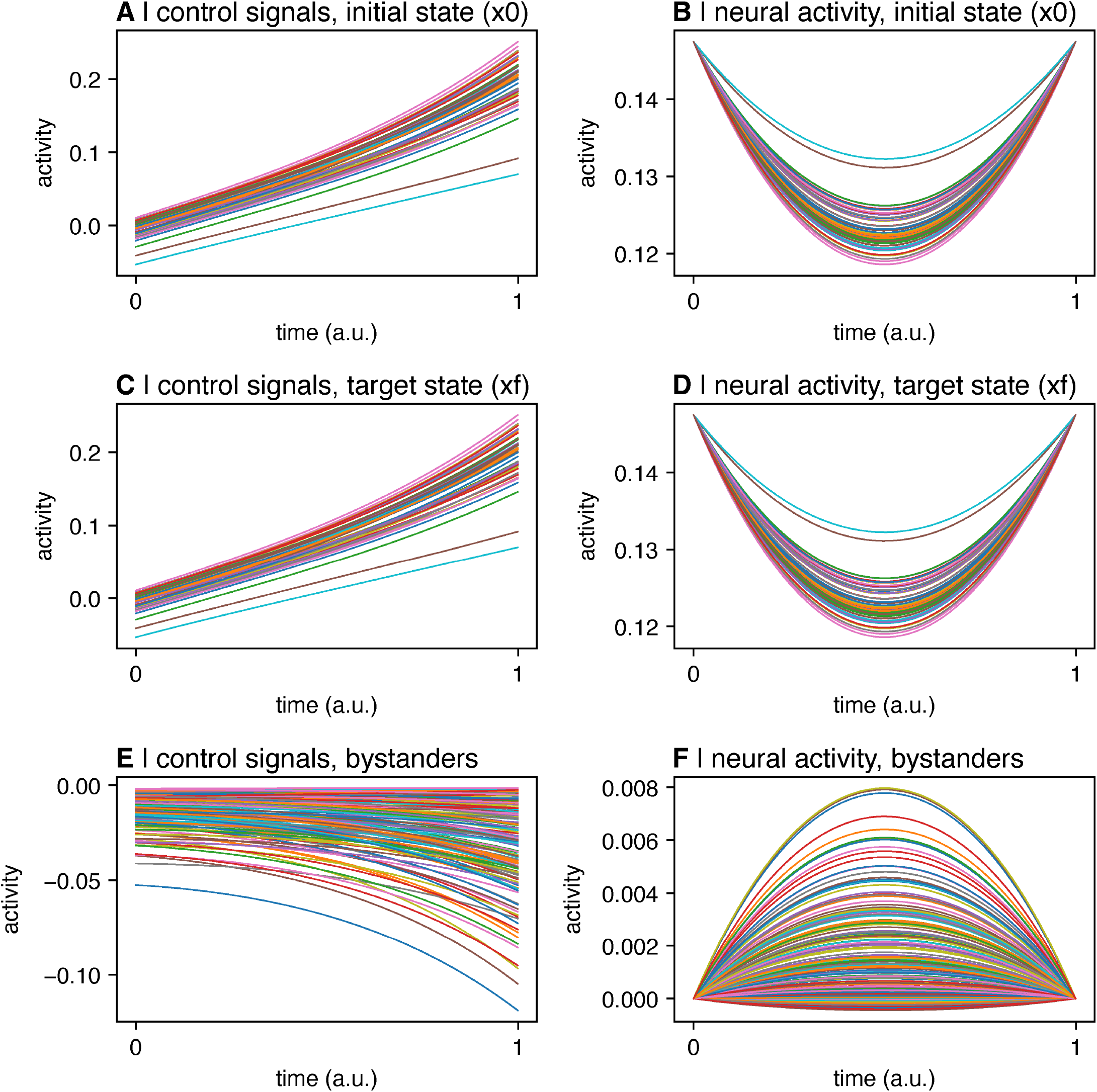
Control signals and state trajectory. Uniform full control set. T=1. *Initial state = default mode system. Target state = default mode system*. Inversion error = 2.20 × 10^*−*16^. Reconstruction error = 3.95 × 10^*−*14^. Energy = 571.

**FIG. S10.**
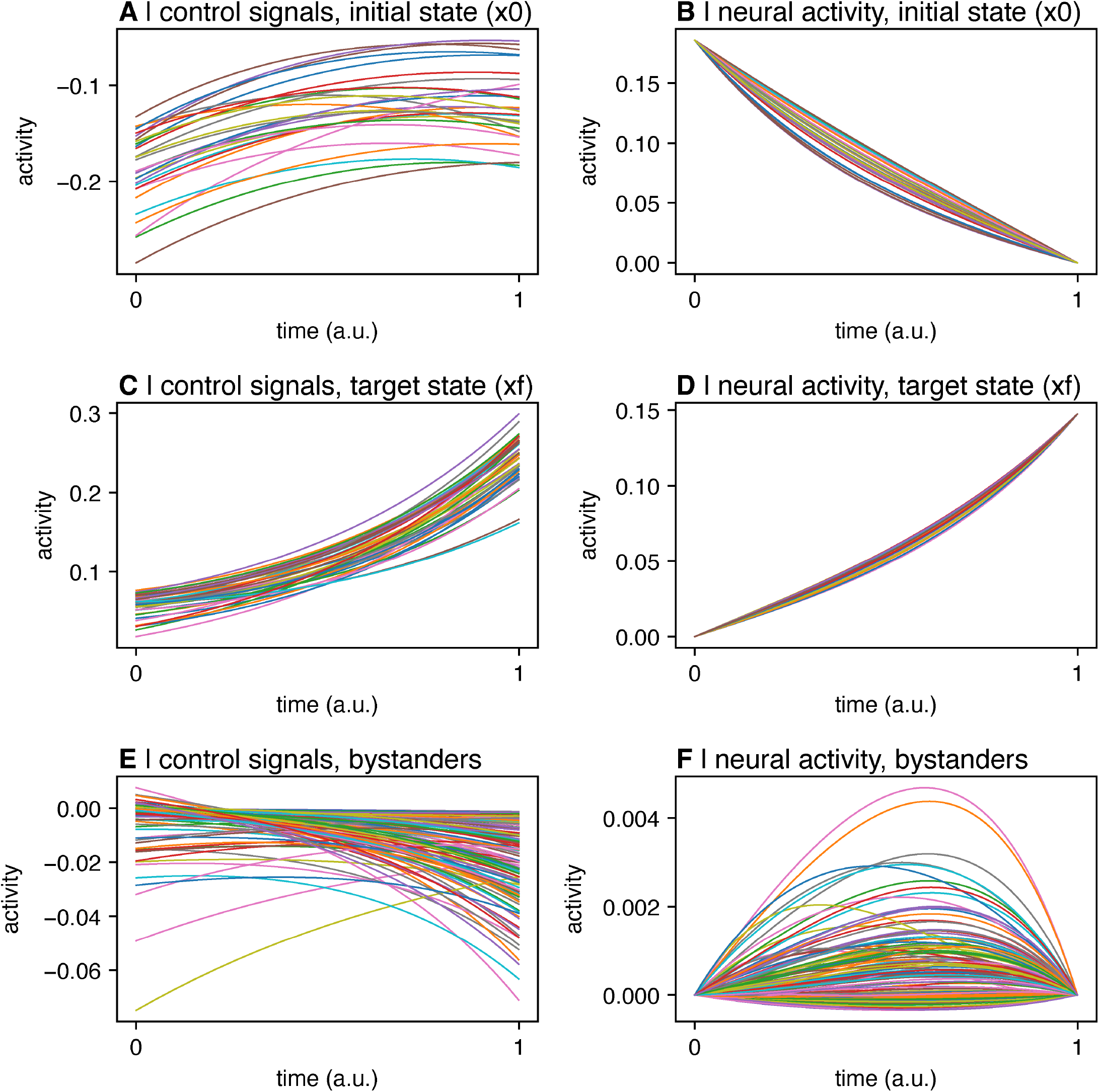
Control signals and state trajectory. *Annotation map control set*. T=1. Initial state = visual system. Target state = default mode system. Inversion error = 1.87 × 10^*−*15^. Reconstruction error = 6.07 × 10^*−*14^. Energy = 1455.

## Notes

https://github.com/BassettLab/nctpy

